# Conglomeration of highly antigenic nucleoproteins to inaugurate a heterosubtypic next generation vaccine candidate against Arenaviridae family

**DOI:** 10.1101/2019.12.29.885731

**Authors:** Kazi Faizul Azim, Tahera Lasker, Rahima Akter, Mantasha Mahmud Hia, Omar Faruk Bhuiyan, Mahmudul Hasan, Md. Nazmul Hossain

## Abstract

Arenaviral infections often resulting in lethal Hemorrhagic Fevers (HF) affect primarily African and South American regions. To date, there is no FDA-approved licensed vaccine against arenaviruses and treatments have been limited to supportive therapies. Hence, the study was employed to design a highly immunogenic heterosubtypic vaccine candidate against Arenaviridae family. The whole proteomes of Lassa virus (LASV), Lymphocytic Chorio Meningitis Virus (LCMV), Lujo virus and Guanarito virus were retrieved from NCBI database and assessed to determine the most antigenic viral proteins. Only the conserved sequences were used for T cell and B cell epitope prediction to ensure protective response against a wide range of viral strains. For each virus, nucleoproteins were identified as most antigenic which generated a plethora of antigenic epitopes. The proposed epitopes were highly conserved (up to 100%) and showed high cumulative population coverage. Moreover, results revealed that among the top epitopes, T cell epitope GWPYIGSRS were conserved in Argentine mammarenavirus (Junin virus) and Brazilian mammarenavirus (Sabia virus), while B cell epitope NLLYKICLSG were conserved in Bolivian mammarenavirus (Machupo virus) and Brazilian mammarenavirus (Sabia virus), indicating the possibility of final vaccine constructs to confer broad range immunity in the host. A total 3 constructs were designed by the combination of top epitopes from each protein along with suitable adjuvant and linkers. Different physicochemical properties revealed the superiority of construct V1 in terms of safety and efficacy. Docking analysis of the refined vaccine structure with different MHC molecules and human immune receptors were also biologically significant. The vaccine receptor complex (V1-TLR3) showed minimal deformability at molecular level. Moreover, construct V1 was compatible for insertion into pET28a(+) vector and heterologous cloning in *E. coli* srain K12. However, the results were based on different sequence analysis and various immune databases. Further wet lab based studies using model animals are highly recommended for the experimental validation of the designed vaccine candidates.

## Introduction

Arenaviral infections, primarily affecting South American and African regions, have traditionally been neglected as tropical diseases (Brisse and Ly, 2019). Although most infections are mild, transmission occurs through rodents to humans, sometimes resulting in severe Hemorrhagic Fevers (HF) with high death rates (Salvato, 2012). They are divided into 2 categories, Old World (eastern hemisphere) and New World (western hemisphere) arenavirus. Old World arenaviruses and the diseases they cause include Lassa virus (Lassa fever), Lujo virus, and Lymphocytic Chorio Meningitis Virus (LCMV) (meningitis, encephalitis, congenital fetal infection, severe disease with multiple organ failure). New World arenaviruses include Junin (Argentine hemorrhagic fever), Machupo (Bolivian hemorrhagic fever), Guanarito (Venezuelan hemorrhagic fever), Sabia (Brazilian hemorrhagic fever) and Chapare virus (Shao et al., 2015). Though, arenaviral outbreaks have been restricted to certain geographic areas, known cases of exportation of arenaviruses from endemic regions and socioeconomic challenges to control the rodent reservoirs locally raised serious concerns about the potential for larger outbreaks in the future. Currently, there are no FDA-approved vaccines for arenaviruses and treatments have been limited to supportive therapy and use of non-specific nucleoside analogs like Ribavirin (Brisse and Ly, 2019; Shoemaker et al., 2015). Though, investigational vaccines exist for Argentine hemorrhagic fever (Ambrosio et al., 2011) and Lassa fever (Carrion et al., 2007), no preventive strategies have been available to treat diseases caused by LCMV, Lujo and Gunarito virus (Daniel et al., 2012; Cheng et al., 2015).

Lassa fever is an animal-borne, or zoonotic, acute viral illness which is endemic in parts of West Africa including Sierra Leone, Liberia, Guinea and Nigeria. Neighboring countries are also at risk, as the animal vector for Lassa virus, “multimammate rat” (*Mastomys natalensis*) is distributed throughout the region. It was estimated that this virus infects roughly 300,000 to 500,000 individuals per year yielding approximately 5,000 deaths (Ogbu et al., 2007; Houlihan et al., 2017). In some areas of Sierra Leone and Liberia, it is known that 10-16% of people admitted to hospitals annually have Lassa fever, demonstrating the serious impact the disease has on the region. Case fatality rates in infected patients can reach more than 80% (Houlihan et al., 2017). Making a correct diagnosis of Lassa fever is made difficult by the wide spectrum of clinical effects that manifest, ranging from asymptomatic to multi-organ system failure and death. Common symptoms associated with cases of LASV are fever, sore throat, red eyes, headache, weakness, retrosternal pain, facial edema, generalized abdominal pain, epistaxis and haemoptysis. In severe cases bleeding from mucousal membranes such as the mouth can also be observed. The only available drug, ribavirin is only effective if administered early in infection (within the first 6 days after disease onset). A fundamental understanding of the mechanisms of antibody-mediated neutralization of Lassa virus may have significant implications for the generation epitope-targeted vaccines. There is no approved vaccine for humans against LASV as of 2019 (Yun et al., 2012).

LCMV is another rodent-borne (*Mus musculus)* prototypic virus of Arenaviridae family that can cause substantial neurological problems, including meningitis, encephalitis, and neurologic birth defects, particularly among prenatal and immune compromised humans (Daniel et al., 2012). LCMV Infections have been reported in Europe, America, Australia, Japan, and may occur wherever infected rodent hosts of the virus are found. Several serologic studies conducted in urban areas have shown that the prevalence of LCMV antibodies in human populations range from 2% to 5% (Wright et al., 1997). A meta-analysis of all reported cases of congenital LCMV infection revealed a mortality rate of 35% by 21 months of age (Bonthius et al., 2007; Wright et al., 1997). Most of the survivors have severe neuro-developmental disorders, including microcephaly, poor somatic growth, profound vision impairment, severe seizure disorders, spastic weakness, and substantial mental retardation (Bonthius et al., 2007; Larsen et al., 2001). An effective antiviral therapy for LCMV infection has not yet been developed and still, there is no vaccine to prevent LCMV infection (Daniel et al., 2012). Although Ribavirin has had mixed success in the treatment of severe infections, but is limited to off-label use and can cause muscular toxicity (Mendenhall et al., 2011). Recently, Lujo virus was isolated as a newly discovered novel arenavirus associated with a VHF outbreak in southern Africa in 2008. It was found to cause a fulminant viral hemorrhagic fever (LUHF) syndrome characterized by nonspecific symptoms such as fever, malaise, myalgias, sore throat, nausea, vomiting and non-bloody diarrhea followed with variable retrosternal or epigastric pain, usually progressing to bleeding, shock and multiorgan failure (Sewlall et al., 2017). This virus has been associated with an outbreak of five cases in September and October 2008 in cities namely Lusaka (Zambia) and Johannesburg (Republic of South Africa). The case fatality rate was 80% (Paweska et al., 2014). In essence, the distribution and prevalence of LUHF in humans and rodents is unknown, as are the ecology, distribution, and mode of transmission from reservoir host to humans. Ribavirin’s effectiveness against Lujo virus remains unknown as well (Paweska et al., 2009).

Guanarito (GTO) virus is the etiological agent of Venezuelan haemorrhagic fever, a rodent borne zoonosis which is endemic in the Northern America (Manzione et al., 1998). Infections are characterized by having the onset of pulmonary congestion and edema, renal and cortical necrosis and haemorrhage in different sites like mucous membranes, major internal organs, digestive and urinary tracts (Craighead, 2000). The short tailed cane mouse, *Zygodontomys brevicauda* (a grassland rodent) acts as a reservoir of Guanarito virus (Fulhorst et al., 1999). Rodent to human transmission may occur via inhalation of virus in aerosolized droplets of secretions or excretions from infected rodents or via contact with the virus through mucosal or cutaneous routes (Ter Meulen et al., 1996). Venezuelan hemorrhagic fever was first recognized as a distinct clinical entity in 1989 during an outbreak of hemorrhagic fever that began in the Municipality of Guanarito in southern Portuguesa (Salas et al., 1991, Tesh et al., 1994). Results of an epidemiologic study of 165 cases of Venezuelan haemorrhagic fever indicated that the disease is seasonal and the number of affected people peaks in november to january. The overall fatality rate was 33.3 % among the 165 cases despite hospitalization and vigorous supportive care (Manzione, 1998). Although successful in many cases traditional vaccines are associated with several demerits (Stratton et al., 2003; Hasan et al., 2019a). Developing vaccines for the organisms not grown in culture is very costly and the yield of vaccines is very low. There is also danger of non-virulent organisms getting converted to virulent ones (Hasson et al., 2015; Kaufmann et al., 2014). Vaccinations by such organisms may cause the disease itself. Recombinant vaccines produced using immunoinformatic approaches, on the contrary, offer some advantages while reducing the time and cost for production (Azim et al., 2019). Hence, the study was conducted to design a highly antigenic polyvalent vaccine against the viruses of Arenaviridae family which are responsible for severe hemorrhagic fevers in human.

## Materials and Methods

In the present study, reverse vaccinology approach was employed to design a novel multiepitope subunit vaccine against the most deadly viruses of Arenaviridae family requiring urgent need for effective medications and preventive measures. The flow chart summarizing the entire protocol of *in silico* strategy for developing a chimeric polyvalent vaccine has been illustrated in Fig. 1.

**Fig. 1:**
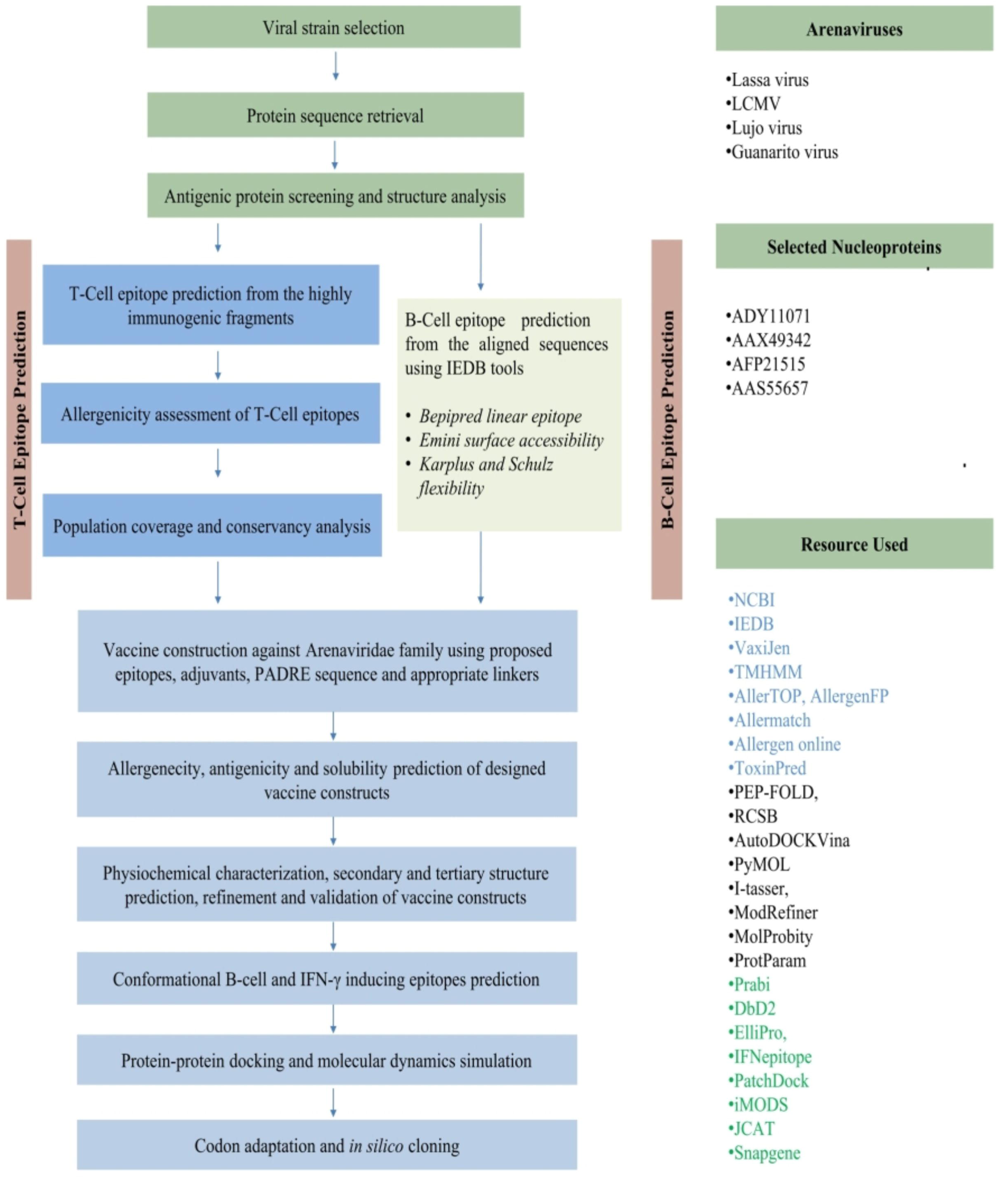
Schematic presentation of the procedures used for multi-epitope vaccine development against Arenaviridae family.

### Retrieval of viral proteomes and antigenic protein selection

NCBI server was used for the selection of viral strains and the whole proteomes of LASV, LCMV, Lujo virus and Guanarito virus were retrieved from the database (https://www.ncbi.nlm.nih.gov/genome/). Study of viral genus, family, host, disease, transmission, and genome were performed by using ViralZone (https://viralzone.ExPASy.org/). The most potent immunogenic proteins were identified individually for all the viruses after determining the antigenicity score via VaxiJen v2.0 server (Doytchinova & Flower, 2007). Different physiochemical parameters of the proteins were analyzed through ProtParam tool (Gasteiger et al., 2003).

### Retrieval of homologous protein sets and identification of conserved regions

The selected proteins from each virus were used as query and homologous sequences were retrieved using BLASTp tool from NCBI. Multiple sequence alignment (MSA) was further performed to find out the common fragments for each set of proteins using CLUSTAL Omega (Sievers et al., 2011) along with 1000 bootstrap value and other default parameters to fabricate the alignment.

### Antigenicity prediction and transmembrane topology analysis of the conserved fragments

The conserved regions from the selected proteins were again screened to demonstrate their antigenicity via VaxiJen v2.0 (Doytchinova & Flower, 2007). The fragments were subjected TMHMM v0.2 server (Krogh et al., 2001) for transmembrane topology prediction. Only the common fragments were used to predict the highly immunogenic T-cell epitopes.

### Prediction of T-cell epitopes, transmembrane topology screening and antigenicity analysis

From the IEDB database (Immune Epitope Database), both MHC-I (http://tools.iedb.org/mhci/) and MHC-II prediction tools (http://tools.iedb.org/mhcii/) were used to predict the MHC-I binding and MHC-II binding peptides respectively (Vita et al., 2014). The TMHMM server (http://www.cbs.dtu.dk/services/TMHMM/) predicted the transmembrane helices in proteins (Krogh et al., 2001). Again, VaxiJen v2.0 server (http://www.ddg-pharmfac.net/vaxijen/) was used to determine the antigenicity of predicted CTL (Cytoxic T-lymphocytes) and HTL (Helper T-lymphocytes) epitopes.

### Population coverage analysis, allergenicity assessment and toxicity analysis

Population coverage analysis is crucial due to variation of HLA distribution among different ethnic groups and geographic regions around the world. In this study, population coverage for each individual epitope was analyzed by IEDB population coverage calculation tool analysis resource (http://tools.iedb.org/population/). The allergenicity pattern of the predicted epitopes were determined through four servers i.e. AllerTOP (Dimitrov et al., 2013), AllergenFP (Dimitrov et al., 2014), Allergen Online (Goodman et al., 2016) and Allermatch (Fiers et al., 2004) were, while the toxicity level was demonstrated using ToxinPred server (Hasan et al., 2019b).

### Epitope conservancy analysis

To determine the extent of desired epitope distributions in the homologous protein set epitope conservancy analysis is a vital step. IEDB’s epitope conservancy analysis tool (http://tools.iedb.org/conservancy/) was selected for the analysis of conservancy level by considering the identities of the selected proteins.

### Designing three-dimensional (3D) epitope structure and molecular docking analysis

The top epitopes were allowed for the docking study after analyzing through different bioinformatics tools. PEP-FOLD server was used to design and retrieve the 3D structure of most potent selected epitopes (Maupetit et al., 2010). The docking was conducted using AutoDOCKVina program at 1.00°A spacing (Morris et al., 2009). The exhaustiveness parameters were kept at 8.00 and the numbers of outputs were set at 10. OpenBabel (version 2.3.1) was used to convert the output PDBQT files in PDB format. The docking interaction was visualized with the PyMOL molecular graphics system, version 1.5.0.4 (https://www.pymol.org/).

### B-Cell epitope prediction and Screening

Three different algorithms from IEDB were used to identify the most potent B cell (BCL) epitopes of the selected antigenic proteins. The algorithms include Bepipred linear epitope prediction, Emini surface accessibility (Emini et al., 1985) and Kolaskar & Tongaonkar antigenicity scale analysis (Kolaskar & Tongaonkar, 1990). The top B cell epitopes were selected based on their allergenicity pattern and VaxiJen score.

### Epitope cluster analysis and vaccine construction

Epitope cluster analysis tool from IEDB (Dhanda et al., 2018) was used to identify the overlapping peptides among the top CTL, HTL and BCL epitopes at minimum sequence identity threshold of 100%. The identified clusters and singletons (unique epitopes) were utilized in a sequential manner to design the final vaccine constructs. Each vaccine proteins started with an adjuvant followed by the top epitopes. Interactions of adjuvants with toll like receptors (TLRs) induce robust immune reactions (Rana & Akhter, 2016). Hence, three different adjuvants were utilized in the study including beta defensin (a 45 mer peptide), L7/L12 ribosomal protein and HABA protein (*M. tuberculosis*, accession number: AGV15514.1). PADRE sequence was also incorporated along with the adjuvant and peptides with a view to overcome the problem caused by highly polymorphic HLA alleles. EAAAK, GGGS, GPGPG and KK linkers were used to conjugate the adjuvant, CTL, HTL and BCL epitopes respectively.

### Allergenicity, antigenicity and solubility prediction of different vaccine constructs

Allergenicity pattern of the designed vaccines were determined by AlgPred v.2.0 (Saha and Raghava, 2000). VaxiJen v2.0 server (Doytchinova and Flower, 2007) was further used to evaluate the probable antigenicity of the constructs in order to suggest the superior vaccine candidate. Protein-sol software (Hebditch et al., 2017) analyzed the solubility score of the proposed vaccine candidates by calculating the surface distribution charge, hydrophocity and the stability at 91 different combinations of pH and ionic strength.

### Physicochemical characterization and secondary structure analysis

ProtParam, a tool provided by ExPASy server (Hasan et al., 2015a) was used to functionally characterize the vaccine proteins. Molecular weight, aliphatic index, isoelectric pH, hydropathicity, instability index, GRAVY values, estimat half-life and other physicochemical properties were analyzed. The Prabi server (https://npsa-prabi.ibcp.fr/) predicted the alpha helix, beta sheet and coil structure of the vaccine constructs through GOR4 secondary structure prediction method.

### Vaccine tertiary structure prediction, refinement, validation and disulfide engineering

I-TASSER server (Zhang, 2010) performed 3D modeling of the designed vaccines depending on the level of similarity between target protein and available template structure in PDB (Hasan et al., 2015b). Refinement was conducted using ModRefiner (Xu and Zhang, 2011) to improve the accuracy of the predicted 3D modeled structure. The refined protein structure was further validated by Ramachandran plot assessment through MolProbity software (Davis et al., 2004). DbD2, an online tool was used to design disulfide bonds for the designed construct (Craig & Dombkowski, 2013).

### Conformational B-cell and IFN-***γ*** inducing epitopes prediction

The conformational B-cell epitopes in the vaccine were predicted via ElliPro server (http://tools.iedb.org/ellipro/) with minimum score 0.5 and maximum distance 7 angstrom (Ponomarenko et al., 2004). Moreover, IFN-γ inducing epitopes within the vaccine were predicted using IFNepitope server (Hajighahramani et al., 2017) with motif and SVM hybrid prediction approach (http://crdd.osdd.net/raghava/ifnepitope/scan.php).

### Protein-protein docking

Different Pattern Recognition Receptors (PRRs) including both membrane associated Toll-like receptors (TLR-3, TLR-7) and cytoplasmic RIG-I-like receptors (RIG-I, MDA5) can recognize infections caused by the members of Arenaviridae (Borrow et al., 2010). Subsequent studies also revealed that α-dystroglycan (αDG) expressed at high levels in skeletal muscle (Ibraghimov-Beskrovnaya et al., 1993) acts as a primary receptor for Old World arenaviruses. The 3D structure of different MHC molecules and human receptors (TLR-3, RIG-I, MDA5, αDG) were retrieved from RCSB protein data bank. Protein-protein docking was conducted to determine the binding affinity of designed vaccines with different HLA alleles and human immune receptors via PatchDock (Hasan et al., 2019c). Docked complexes from PatchDock were subjected to the FireDock server to refine the complexes.

### Molecular dynamics simulation

Molecular dynamics study was performed to strengthen the in silico prediction via iMODS server (Lopez-Blanco et al., 2017). The structural dynamics of protein complex (V1-Toll like receptor 3) was investigated by using this server due to its much faster and effective assessments than other molecular dynamics (MD) simulations tools (Awan et al., 2017). The iMODS server explained the collective motion of proteins by analyzing the normal modes (NMA) in internal coordinates (Tama & Brooks, 2006). Stability was determined by comparing the essential dynamics of proteins to their normal modes (Aalten et al., 1997). The direction of the complex and extent of the motions was predicted in terms of deformability, eigenvalues, B-factors and covariance.

### Codon adaptation and in silico cloning

Codon adaptation was performed to accelerate the expression of construct V1 in *E. coli* strain K12. JCAT server was used for this purpose, while Rho independent transcription termination, prokaryote ribosome-binding site and cleavage sites of several restriction enzymes (i.e. BglII and BglI) were avoided during the operation (Grote et al., 2005). The optimized sequence of vaccine protein V1 was reversed and then conjugated with BglII and BglI restriction site at the N-terminal and C-terminal sites respectively. To insert the adapted sequence into pET28a(+) vector between the BglII (401) and BglI (2187), SnapGene (Solanki & Tiwari, 2018) restriction cloning module was utilized.

## Results

### Retrieval of viral proteomes and antigenic protein selection

The entire viral proteomes of LASV, LCMV, Lujo virus and Guanarito virus were retrieved from NCBI protein database in FASTA format. Only the structural proteins proteins were prioritized for vaccine candidacy. Among the rescued proteins, Nucleoproteins from Lassa virus (AAX49342) and LCMV (ADY11071), and Nucleocapsid proteins from Guanarito virus (AAS55657) and Lujo virus (AFP21515) were selected for vaccine development on the basis of highest antigenicity score (Supplementary file 1). Different physiochemical parameters of the proteins analyzed via ProtParam tool are shown in Table 1.

**Table 1:** ProtParam analysis of selected antigenic proteins

| ***Virus*** | ***Selected proteins*** | ***Accession ID*** | ***Molecular weight*** | ***Instability index*** | ***Half life*** | ***pI*** | ***Amino acids*** | ***Extinction coefficient*** |
| --- | --- | --- | --- | --- | --- | --- | --- | --- |
| Lassa virus | Nucleoprotein | ADY11071 | 62998.24 | 36.64 | >10h | 8.6 | 569 | 53860 |
| LCMV | Nucloprotein | AAX49342 | 62177.47 | 40.20 | >10h | 8.8 | 558 | 56380 |
| Lujo virus | Nucleocapsid protein | AFP21515 | 63104.55 | 39.35 | 30h | 8.7 | 558 | 51630 |
| Guanarito virus | Nucleocapsid protein | AAS55657 | 62345.44 | 34.93 | >10h | 8.2 | 560 | 46410 |

### Retrieval of homologous protein sets and identification of conserved regions

Different homologous protein sets for each protein were generated after BLASTp search using NCBI BLAST tools. A total of 5, 12, 3, and 9 conserved fragments were found among the LASV, LCMV, Lujo and Guanarito viral proteins respectively (Table 2).

**Table 2:**
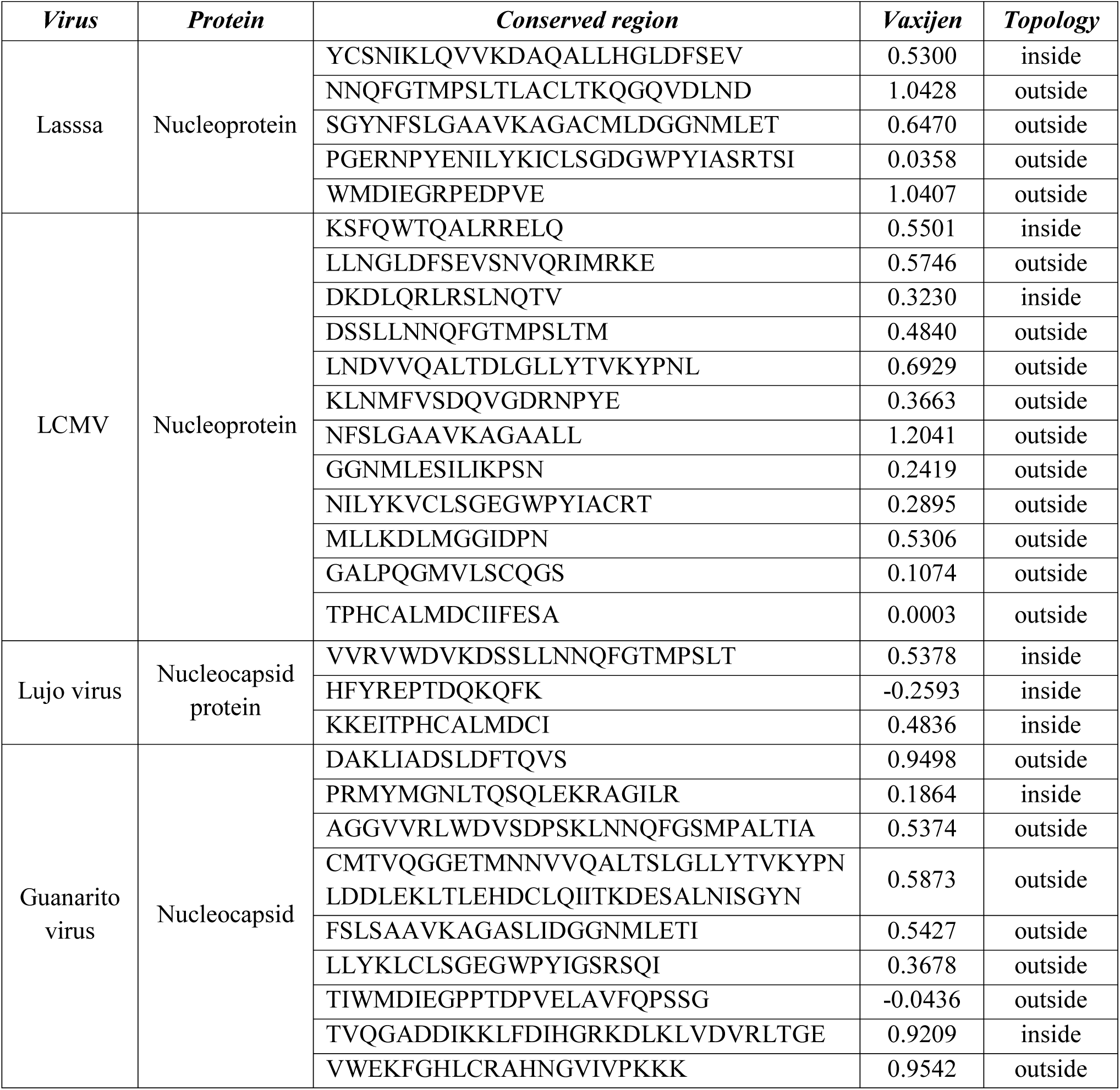
Identified conserved regions among different homologous protein sets of LASV, LCMV, Lujo virus and Guanarito virus

### Antigenicity prediction and transmembrane topology analysis of the conserved fragments

Results showed that 4, 6, 2, 6 and conserved sequences from identified Lassa, LCMV, Lujo, and Guanarito viral proteins respectively met the criteria of default threshold level in VaxiJen (Table 2). Moreover, transmembrane topology screening revealed, among the immunogenic conserved sequences 3, 2, 5 and 5 sequences from the corresponding proteins fulfilled the criteria of exomembrane characteristics (Table 2).

### Prediction of T-cell epitopes, transmembrane topology screening and antigenicity analysis

Numerous immunogenic epitopes from the conserved sequences were generated that could bind maximum number of HLA cells with high binding affinity (Supplementary file 2 and Supplementary file 3). Top epitopes from each of the protein were selected as putative T cell epitope candidates based on their transmembrane topology screening and antigenicity score (Table 3). Epitopes with a positive score of immunogenicity exhibited potential to elicit effective T-cell response.

**Table 3:**
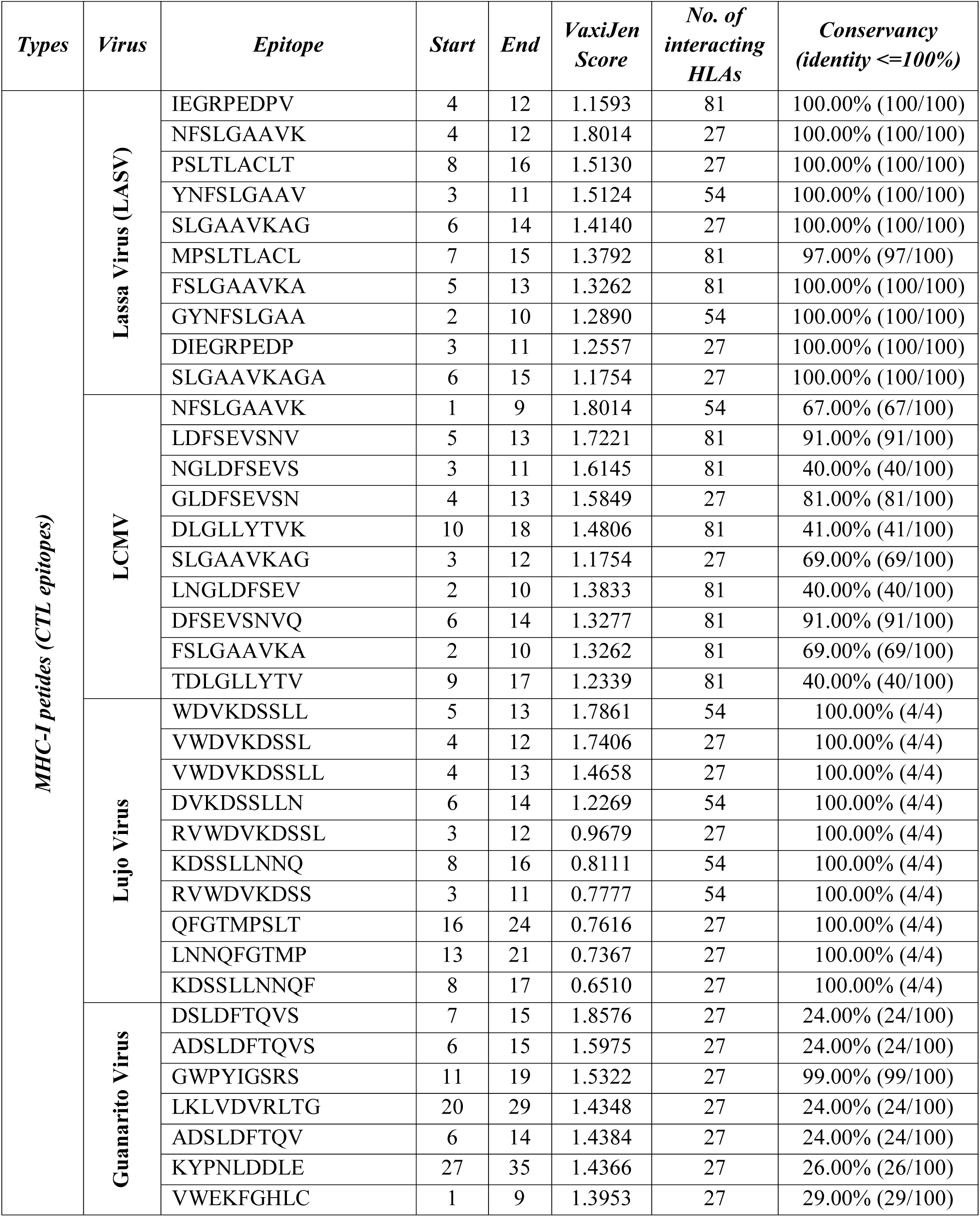
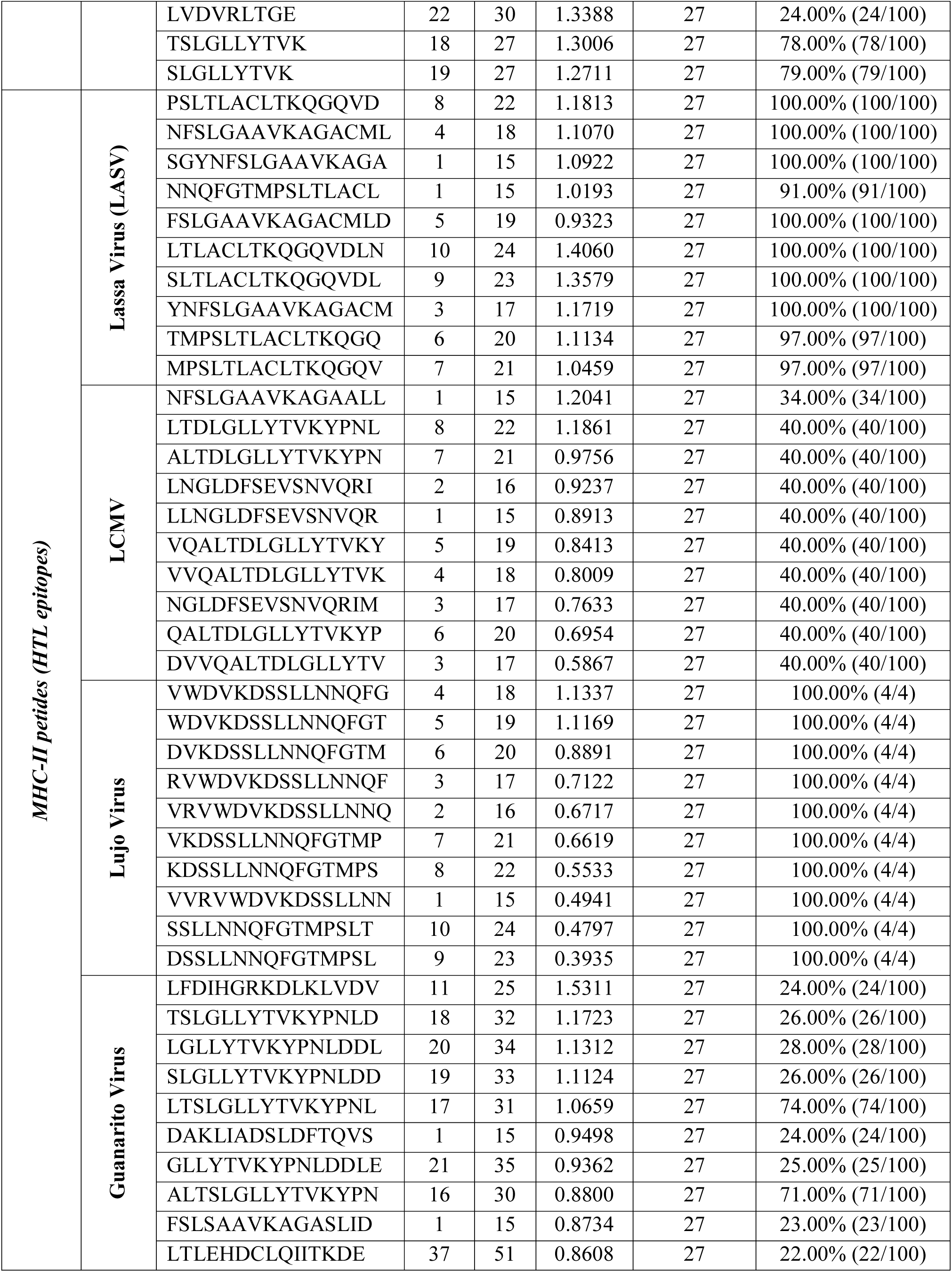
Predicted T-cell (CTL and HTL) epitopes of Nucleoproteins (Lassa virus and LCMV) and Nucleocapsid proteins (Lujo virus and Guanarito virus)

### Population coverage, allergenicity assessment and toxicity analysis of T-cell epitopes

Results showed that population from the most geographic areas can be covered by the predicted T-cell epitopes. Population coverage results for the epitopes of four different viral proteins are shown in Fig. 2. Through the allergenicity assessment by four servers (i.e. AllerTOP, AllergenFP, Allergen online, Allermatch), epitopes that were found to be non-allergen for human were identified (Supplementary file 4 and Supplementary file 5). Epitopes those were indicated as allergen for human and classified as toxic or undefined were removed from the predicted list of epitopes (Table 3).

**Fig. 2:**
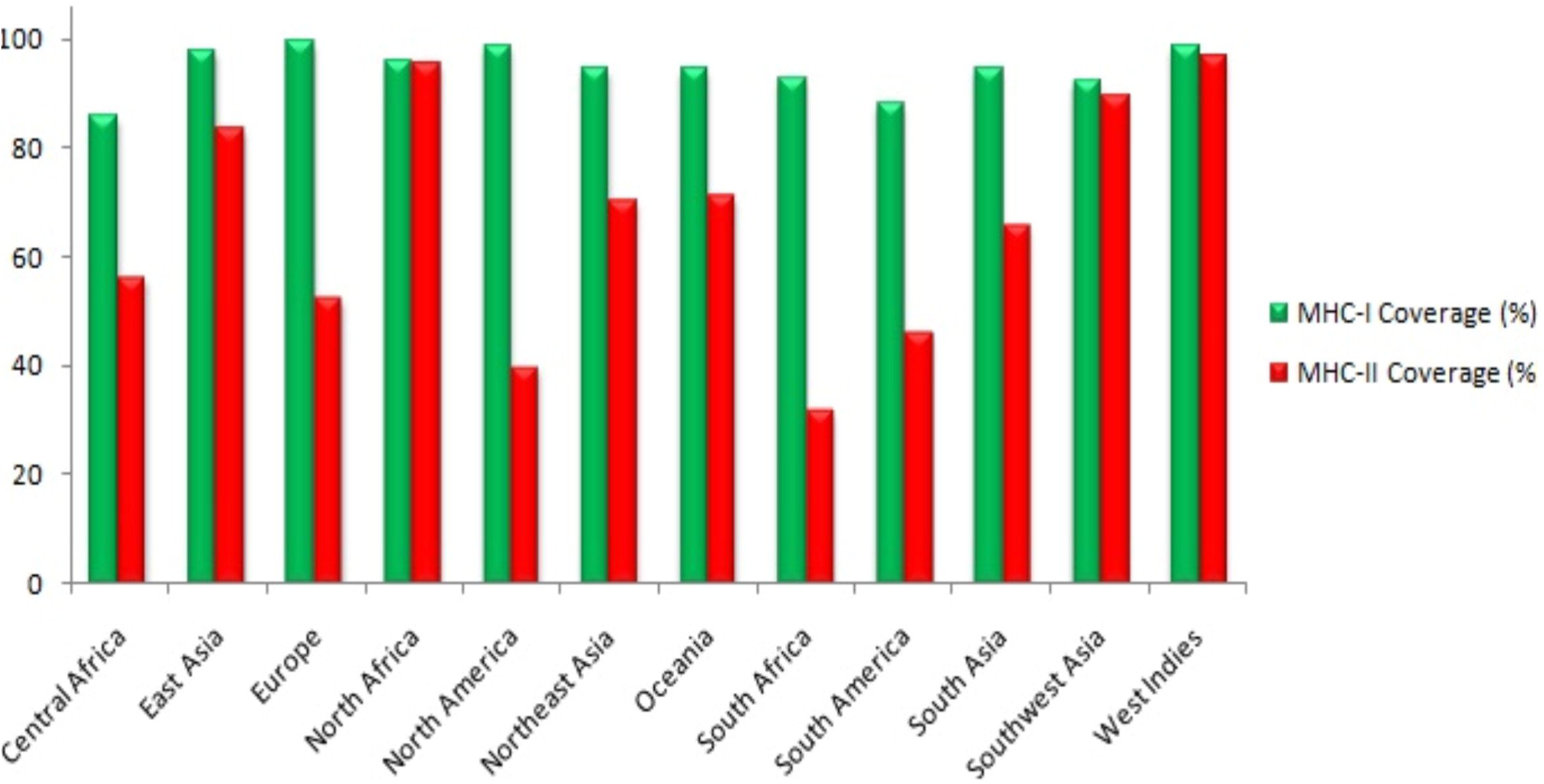
Population coverage analysis of predicted T-cell epitopes (MHC-I and MHC-II peptides).

### Epitope conservancy analysis

Putative epitopes generated from 4 diverse viral proteins were found to be highly conserved within different strains (Table 3). The top epitopes showing conservancy at a superior level (ranging from 40% to 100%) were allowed for further docking study and used to design the final vaccine constructs to ensure a broad spectrum vaccine efficacy (Table 4).

**Table 4:**
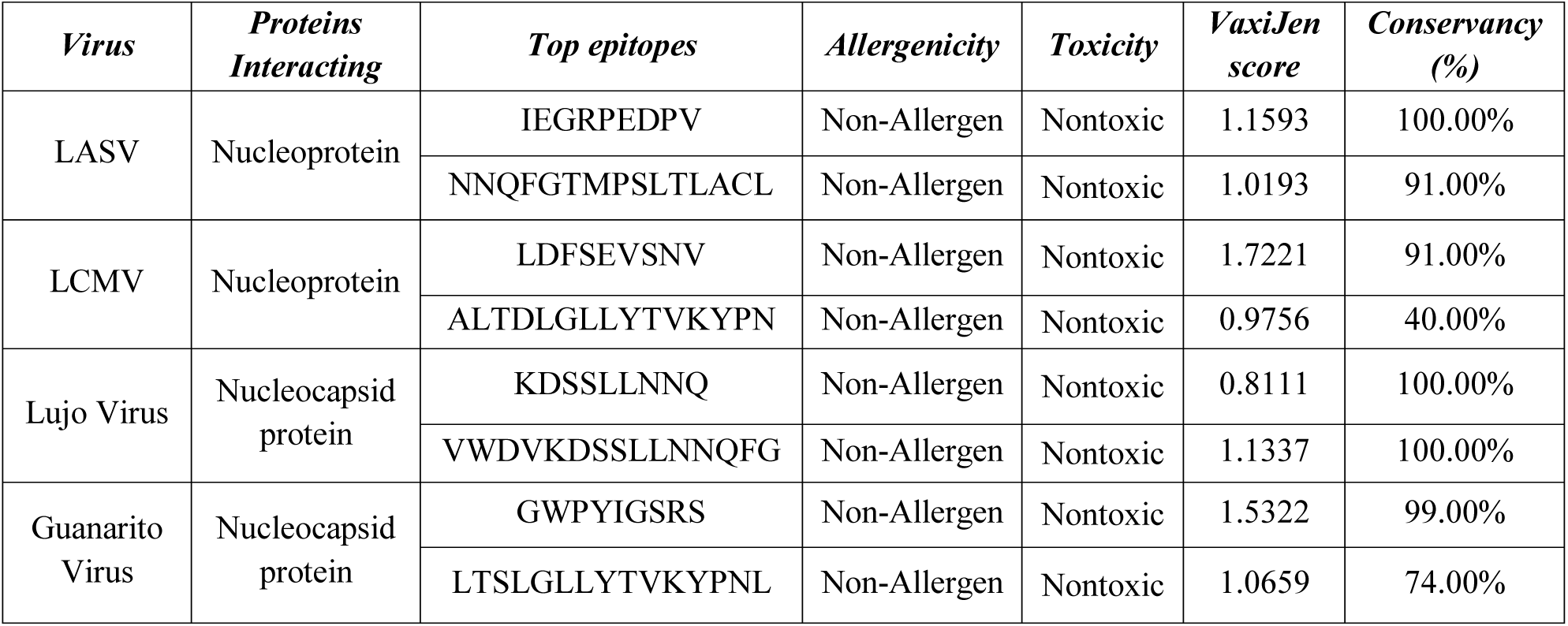
Proposed CTL and HTL epitopes for vaccine construction

### Designing three-dimensional (3D) epitope structure and molecular docking analysis

A total of 8 T-cell (4 CTL and 4 HTL) epitopes were subjected to PEP-FOLD server for 3D structure conversion and their interactions with HLA molecules were anlyzed. HLA-A*11:01 (Class-I) and HLA-DRB1*01:01 (Class-II) were selected for docking analysis based on available PDB structures deposited in the database. Results confirmed that all the predicted epitopes bound in the groove of MHC molecules with a negative binding energy which were biologically significant (Table 5).

**Table 5:**
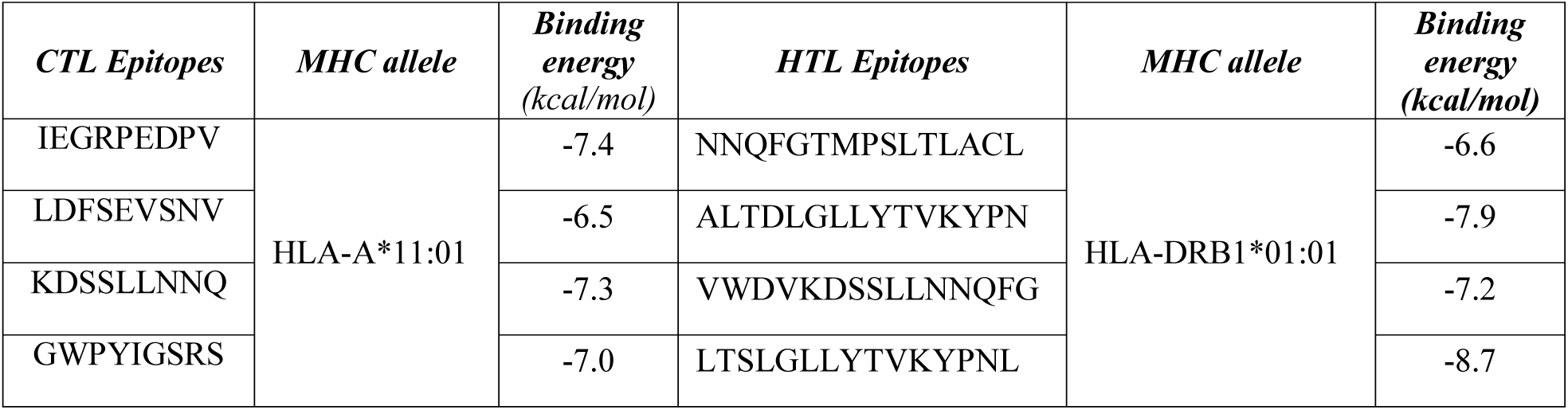
Binding energy of predicted epitopes and MHC alleles generated from molecular docking by AutoDock.

### B-cell epitope prediction and screening

B-cell epitopes were predicted according to the analysis via three different algorithms (i.e. Bepipred Linear Epitope prediction, Emini Surface Accessibility, Kolaskar & Tongaonkar Antigenicity prediction) from IEDB. Top BCL epitopes for selected four proteins were further screened based on their antigenicity scoring and allergenicity pattern which were used to design the final vaccine construct (Table 6).

**Table 6:**
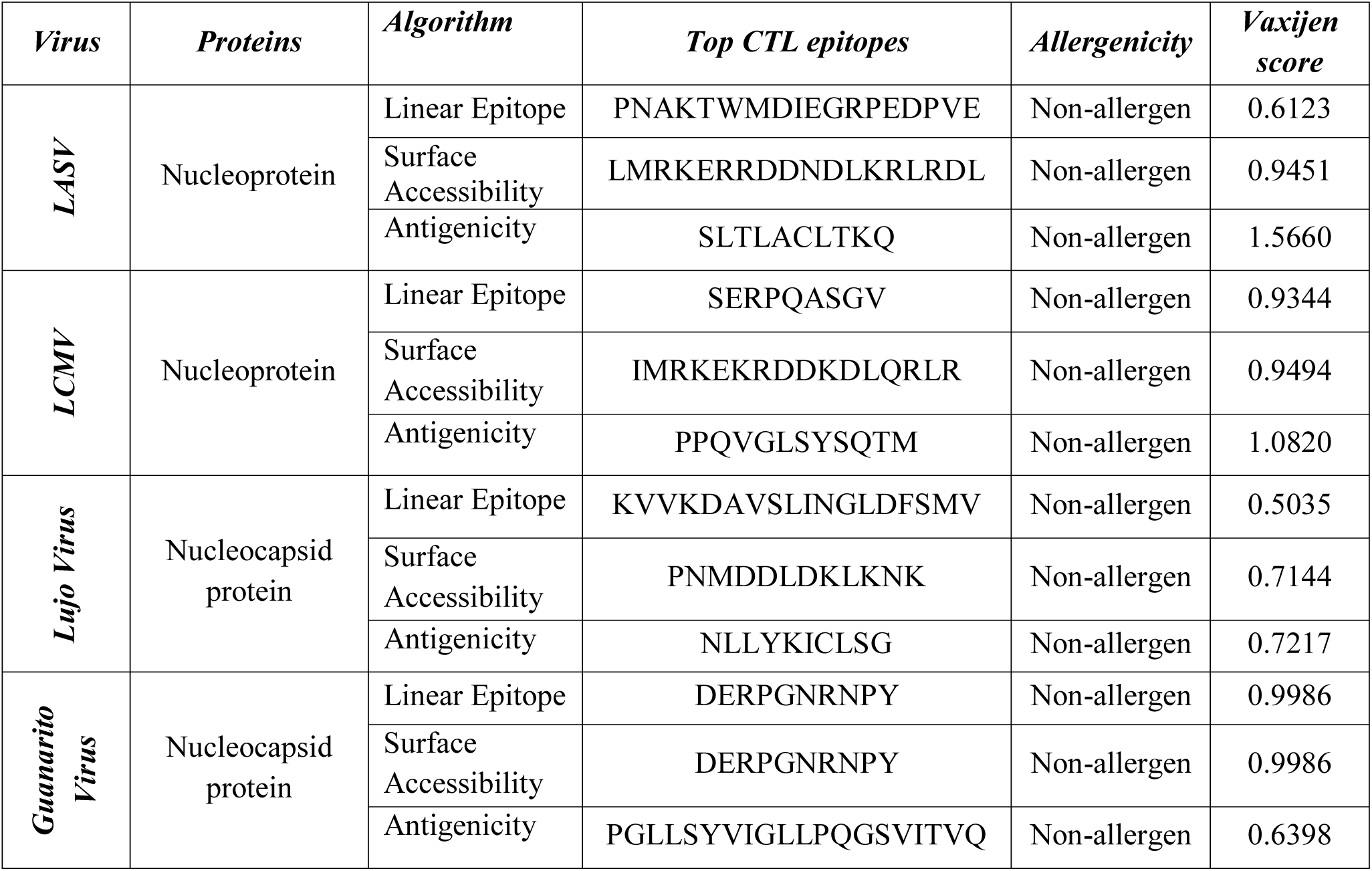
Allergenicity assessment antigenicity analysis of the predicted B-cell epitopes

### Epitope cluster analysis and vaccine construction

Epitope cluster analysis tool from IEDB predicted the clusters among the top epitopes (4 CTL, 4 HTL and 12 BCL epitopes) proposed in Table 4 and Table 6. A total 19 clusters were identified which were utilized to design vaccine constructs in the study. Each construct designed were composed of a protein adjuvant followed by T-cell and B-cell epitopes with their respective linkers. PADRE sequence was incorporated to maximize the efficacy and potency of the peptide vaccine. A total 3 vaccine molecules (i.e. V1, V2, and V3) of 394, 479 and 508 amino acid residues were constructed (Table 7) and further investigated to evaluate their immunogenic potential.

**Table 7:**
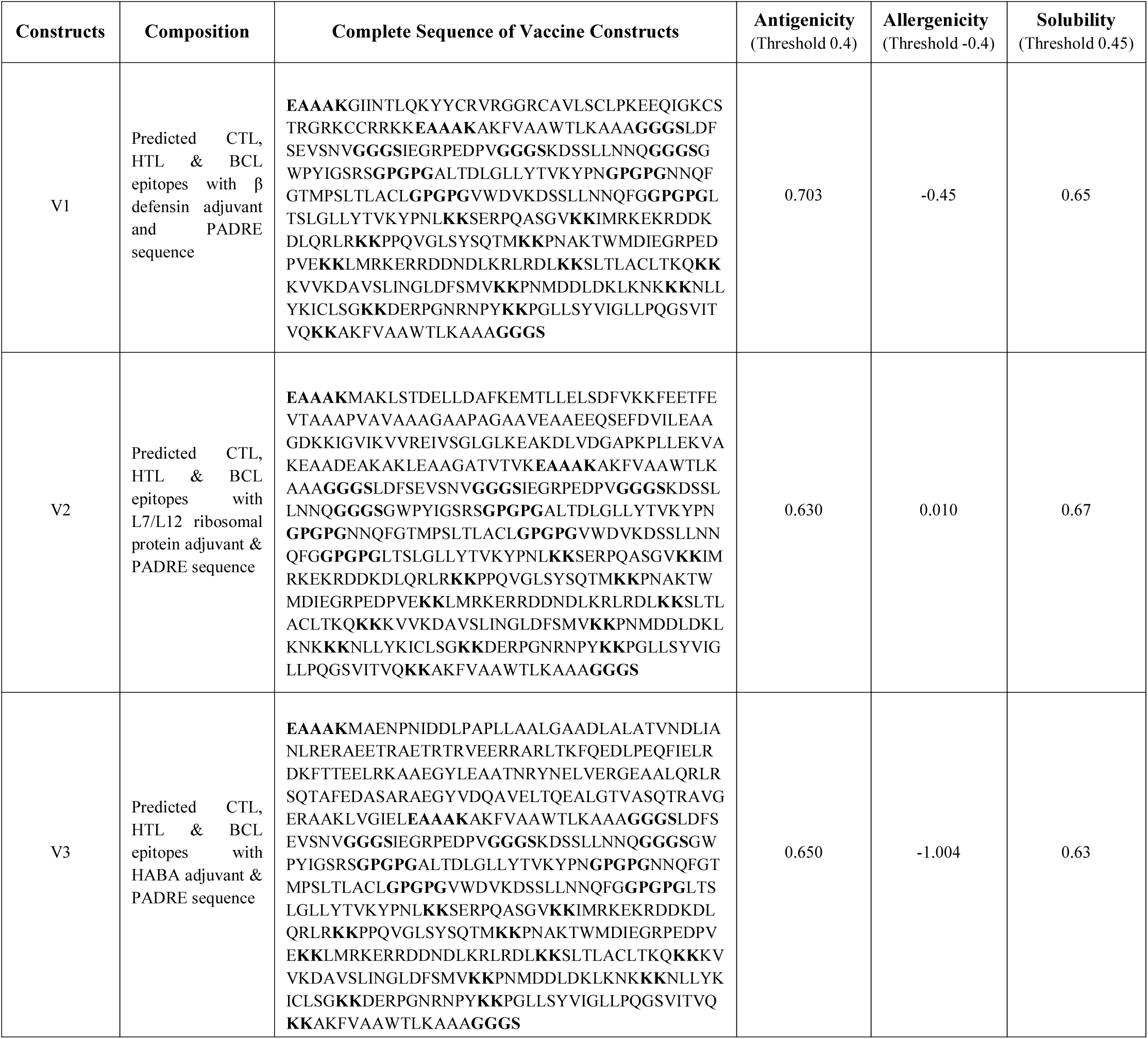
Allergenicity, antigenicity and solubility prediction of the designed vaccine constructs

### Allergenicity, antigenicity and solubility prediction of different vaccine constructs

Results showed that construct V1 and V2 were non-allergic in behavior, while V3 exhibited allergenic pattern (Table 7). However, V1 was best in terms of safety and found superior as potential vaccine candidate with better antigenicity (0.703) and solubility score (0.65) (Fig. 3).

**Fig. 3:**
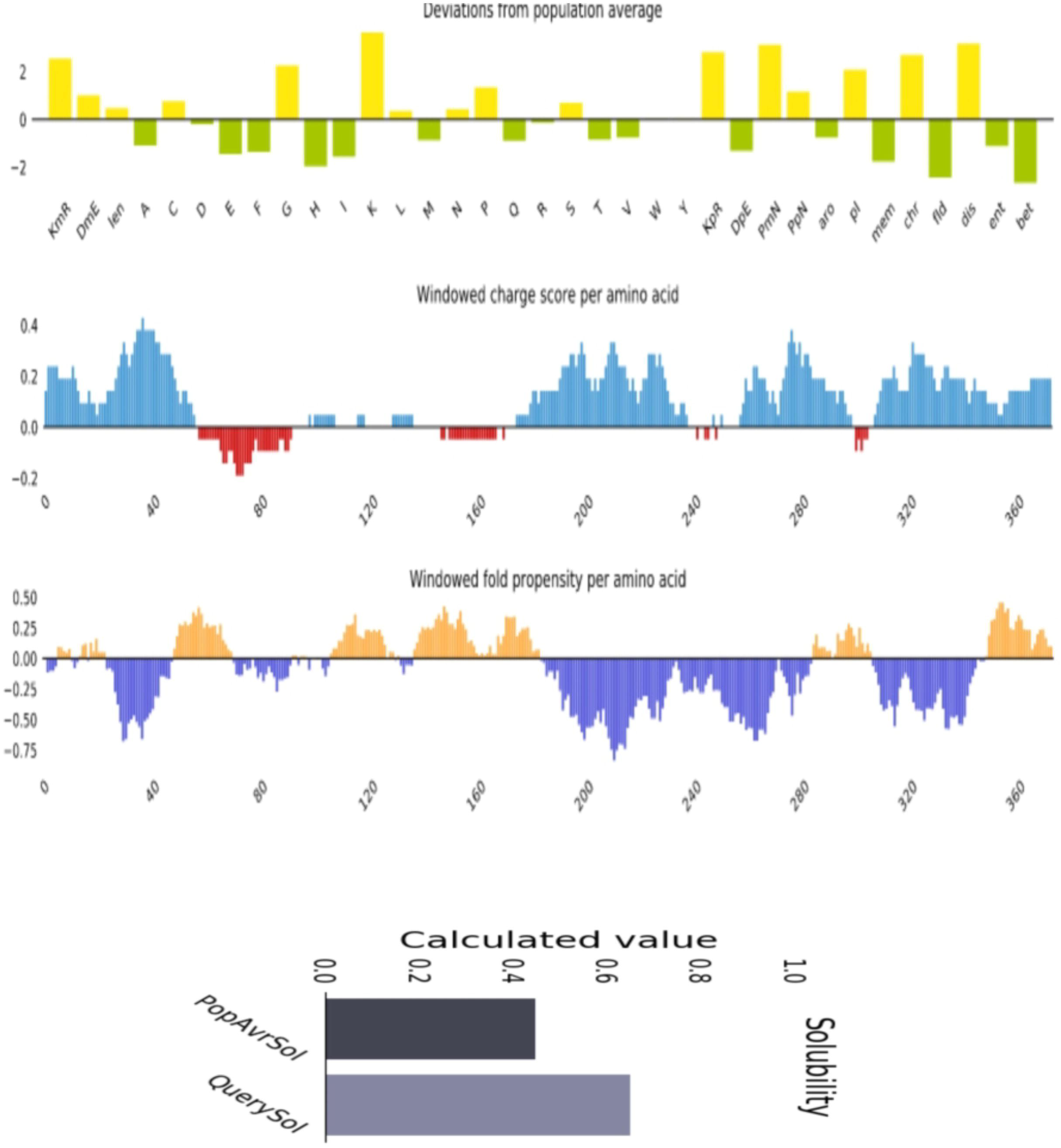
Solubility prediction of designed vaccine protein V1 using via Protein-sol server.

### Physicochemical characterization and secondary structure analysis of vaccine protein

Vaccine construct V1 was characterized on the basis of physical and chemical properties. The computed instability index of the protein was 35.87 which classified it as a stable one. The theoretical pI 9.96 indicated that the protein will have net negative charge above the pI. Assuming all cysteine residues are reduced at 0.1% absorption, the extinction coefficient evaluated was 43890. The estimated half-life of the vaccine was expected to be 1 h in mammalian reticulocytes in vitro, while >10 h in *E. coli* and 30 minutes in yeast in vivo. Thermostability and hydrophilic nature of the vaccine protein was represented by aliphatic index (75.25) and GRAVY value (-0.625). Secondary structure of the construct V1 confirmed to have 26.14% alpha helix, 20.56% sheet and 53.30% coil structure (Fig. 4).

**Fig. 4:**
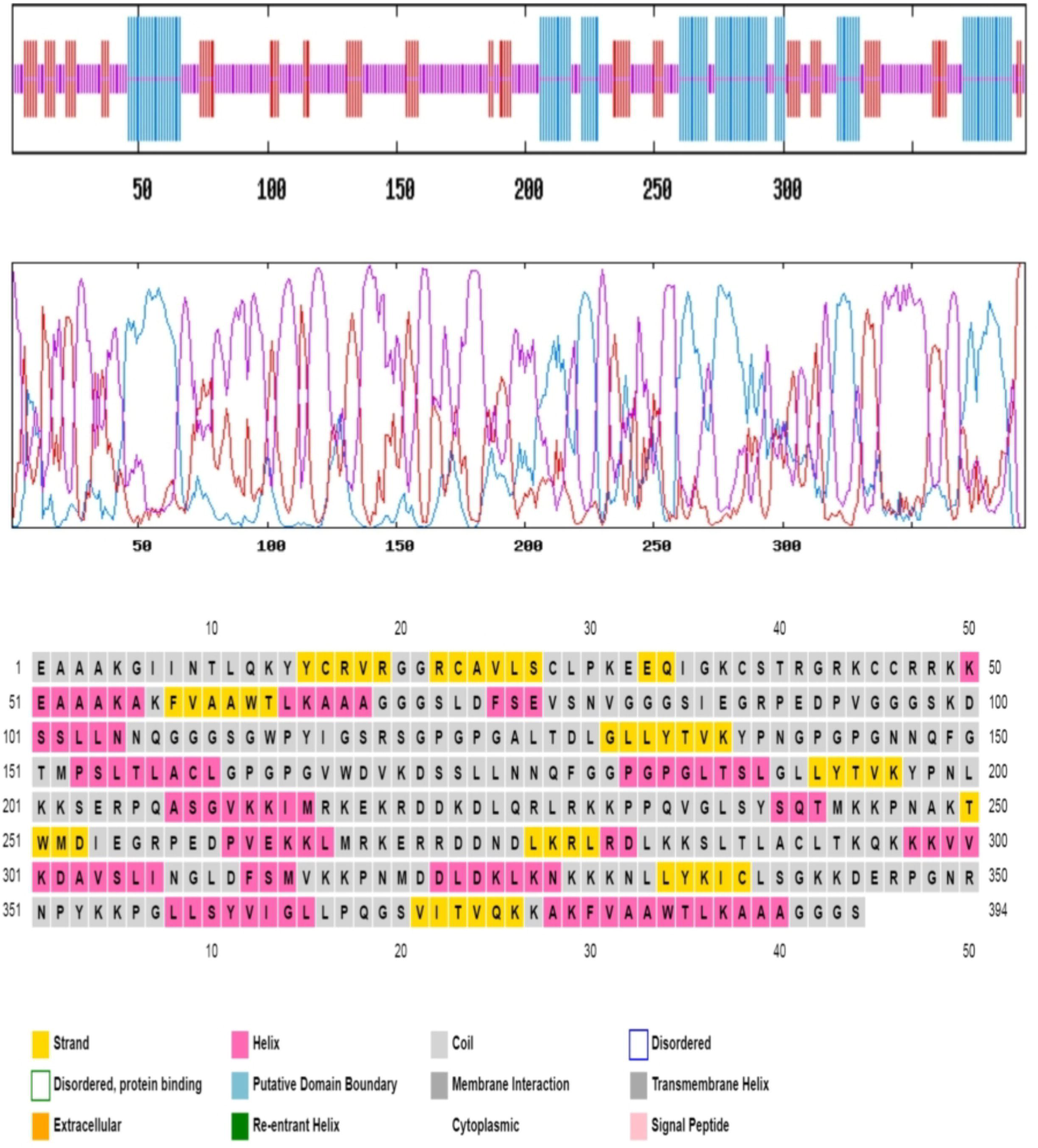
Secondary structure analysis of the designed construct V1.

### Vaccine tertiary structure prediction, refinement, validation and disulfide engineering

I-TASSER generated 5 tertiary structures of the designed construct V1 using top 10 threading templates by LOMETS. TM score and RMSD were estimated based on C score which was minimum for Model 1 (-0.77), thus ensuring its better quality (Fig. 5A). The refined structure was validated through Ramachandran plot analysis which revealed that 94.6% residues were in the allowed and 5.36% residues in the outlier region (Fig. 5B). Modeled tertiary structure of vaccine construct V2 and V3 have been shown in Fig. 6. A total 29 pairs of amino acid residue were identified having the capability to form disulfide bond by DbD2 server. However, after evaluation of the residue pairs in terms of energy, chi3 and B-factor parameter, only 2 pairs (ALA 24-CYS28 and GLU77-GLU87) satisfied the criteria for disulfide bond formation which were replaced with cysteine. The value of chi3 considered for the residue screening was between –87 to +97 while the energy value was less than 2.5.

**Fig. 5:**
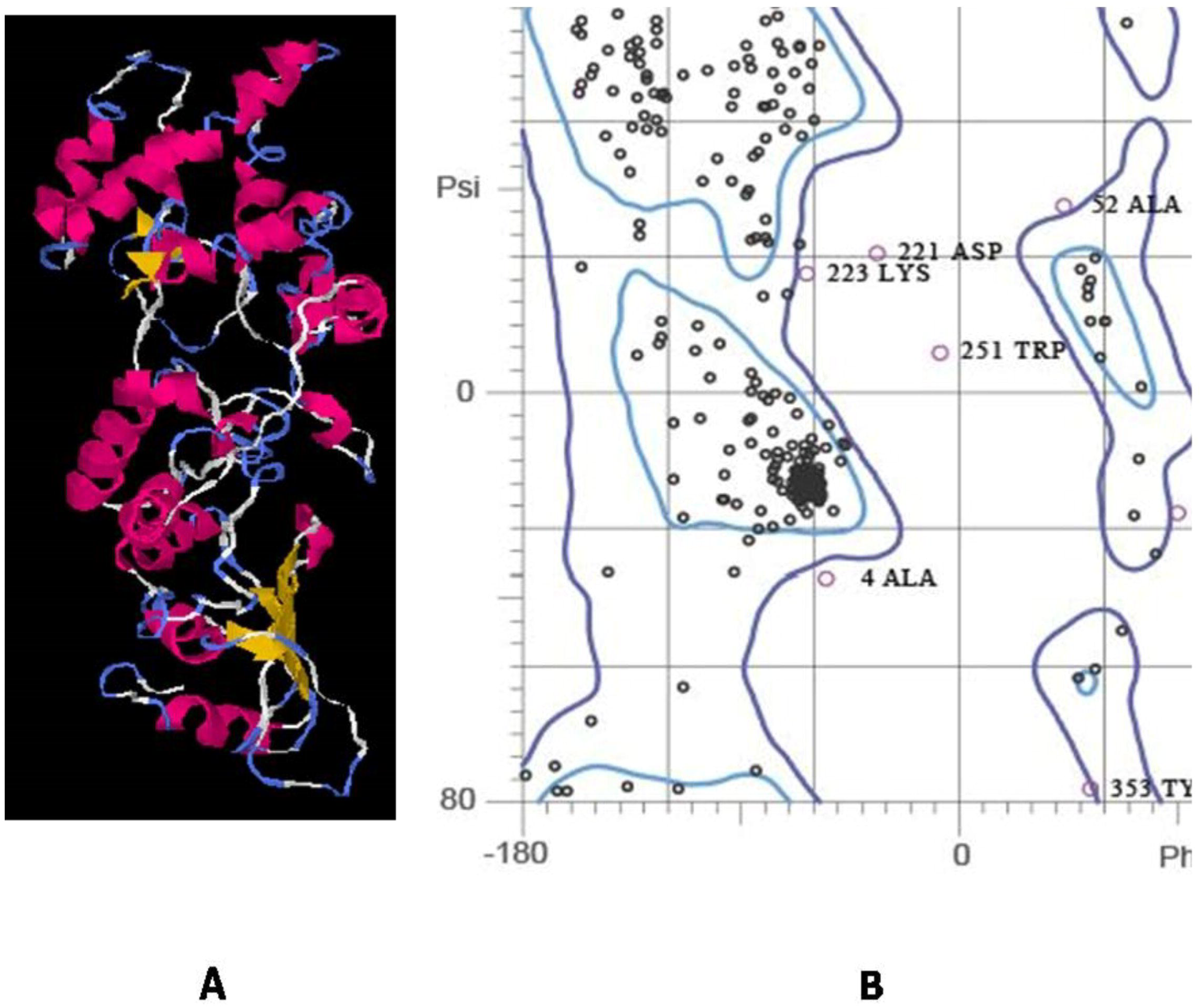
Homology modeling of vaccine protein V1 via I-TASSER (A) and validation of the 3D smodel via by Ramachandran plot analysis (B).

**Fig. 6:**
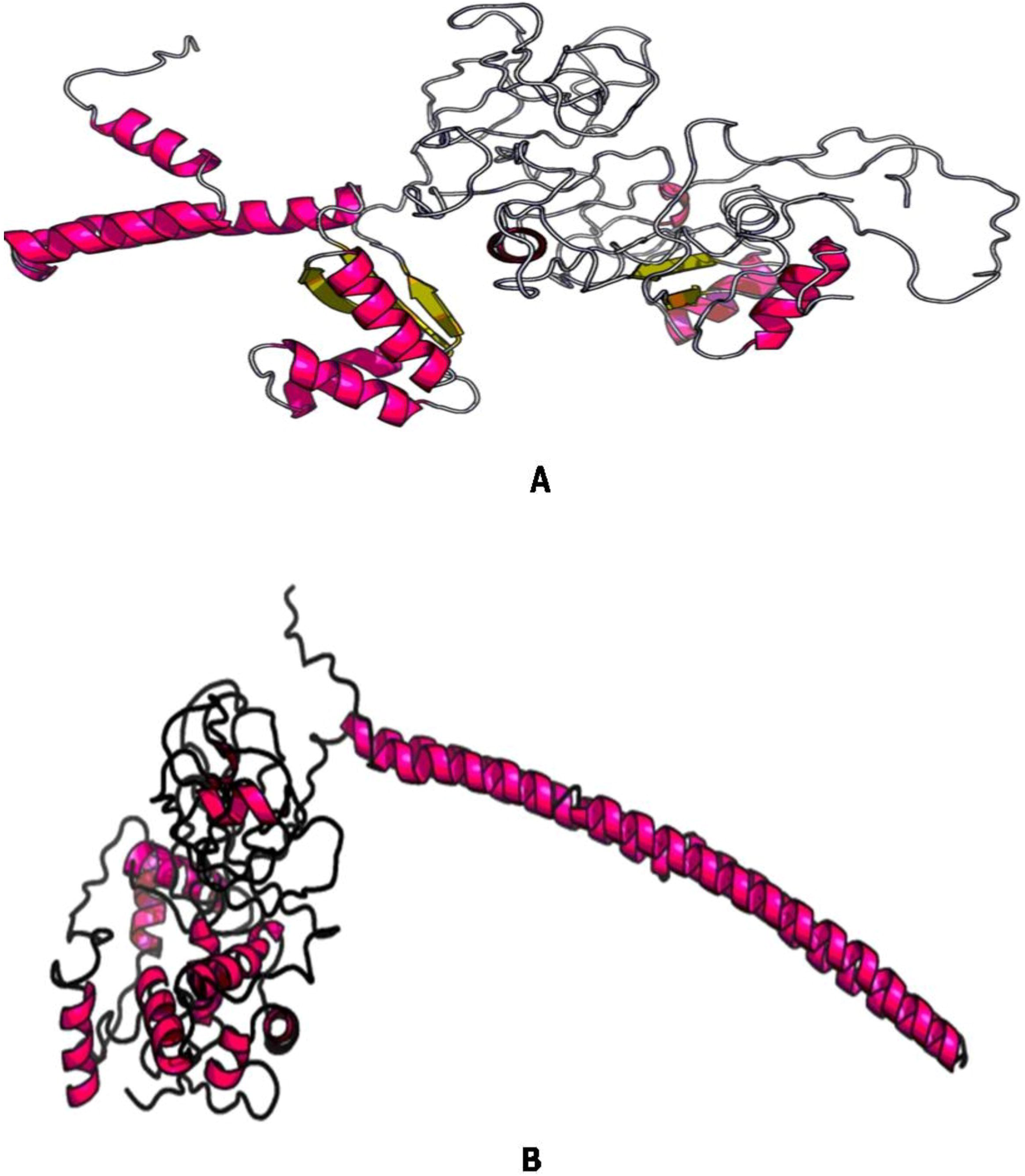
3D modelled structure of vaccine protein V2 (A) and V3 (B).

### Conformational B-cell and IFN-***γ*** inducing epitopes prediction

A total of 5 conformational B-cell epitopes were predicted by using 3D structure of the proposed vaccine as an input. Epitopes No. 2 and 4 were considered as the broadest and smallest conformational B-cell epitopes with 85 and 13 amino acid residues (Table 8). Results also revealed that most of the residues, which were located in the multi-epitope region in our designed vaccine were included in the predicted conformational B-cell epitopes Moreover, the sequence of the final vaccine was applied for prediction of 15-mer IFN-γ inducing epitopes. Results showed that there were 100 positive IFN-γ inducing epitopes from which 8 had a score ≥1 (Supplementary file 6). Residues of 375-390 regions in the vaccine showed highest score of 1.209 (Table 9).

**Table 8:** Conformational B-cell epitopes from 3D model of the vaccine construct V1

| No. | Residues | No. of residues | Score |
| --- | --- | --- | --- |
| 1 | A:A66, A:A67, A:A68, A:G69, A:G70, A:G71, A:S72, A:L73, A:D74, A:F75, A:S76, A:E77, A:V78, A:S79, A:N80, A:V81, A:G82, A:I86, A:D276, A:L277, A:K278, A:R279, A:L280, A:R281 | 24 | 0.758 |
| 2 | A:S98, A:K99, A:D100, A:S101, A:S102, A:L103, A:L104, A:N105, A:N106, A:Q107, A:G108, A:G109, A:G110, A:L127, A:T128, A:D129, A:L130, A:I307, A:N308, A:G309, A:L310, A:D311, A:F312, A:S313, A:M314, A:V315, A:K316, A:K317, A:P318, A:N319, A:M320, A:D321, A:D322, A:L323, A:D324, A:N328, A:K329, A:K330, A:K331, A:N332, A:L333, A:L334, A:S340, A:G341, A:K342, A:K343, A:D344, A:E345, A:R346, A:P347, A:G348, A:N349, A:R350, A:N351, A:P352, A:Y353, A:K354, A:K355, A:L366, A:P367, A:Q368, A:G369, A:S370, A:V371, A:I372, A:T373, A:V374, A:K377, A:A378, A:K379, A:F380, A:V381, A:A382, A:A383, A:W384, A:T385, A:L386, A:K387, A:A388, A:A389, A:A390, A:G391, A:G392, A:G393, A:S394 | 85 | 0.716 |
| 3 | A:E1, A:A2, A:A3, A:A4, A:K5, A:G6, A:I8, A:N9, A:T10, A:K13, A:G163, A:P164, A:G165, A:V166, A:W167, A:D168, A:V169, A:K170, A:D171, A:S172, A:S173, A:L174, A:L175, A:N176, A:N177, A:Q178, A:F179, A:G180, A:G181, A:P184, A:T187, A:S188, A:G190, A:L191, A:L192, A:T194, A:V195, A:K196, A:Y197, A:P198, A:N199, A:L200, A:K201, A:K202, A:S203, A:E204, A:P206, A:Q207, A:A208, A:S209, A:G210, A:V211, A:K212, A:K213, A:I214, A:M215, A:R216, A:K217, A:E218, A:K219, A:R220, A:D221, A:D222, A:K223, A:Q226, A:K230, A:T250, A:W251, A:D253, A:I254, A:E255, A:G256, A:R257, A:P258, A:E259 | 75 | 0.671 |
| 4 | A:D282, A:L283, A:K284, A:K285, A:S286, A:L287, A:T288, A:L289, A:A290, A:C291, A:L292, A:T293, A:G357 | 13 | 0.591 |
| 5 | A:C28, A:K31, A:E32, A:E33, A:Q34, A:I35, A:G36, A:K37, A:C38, A:T40, A:R41, A:K44, A:C45, A:R47 | 14 | 0.56 |

**Table 9:**
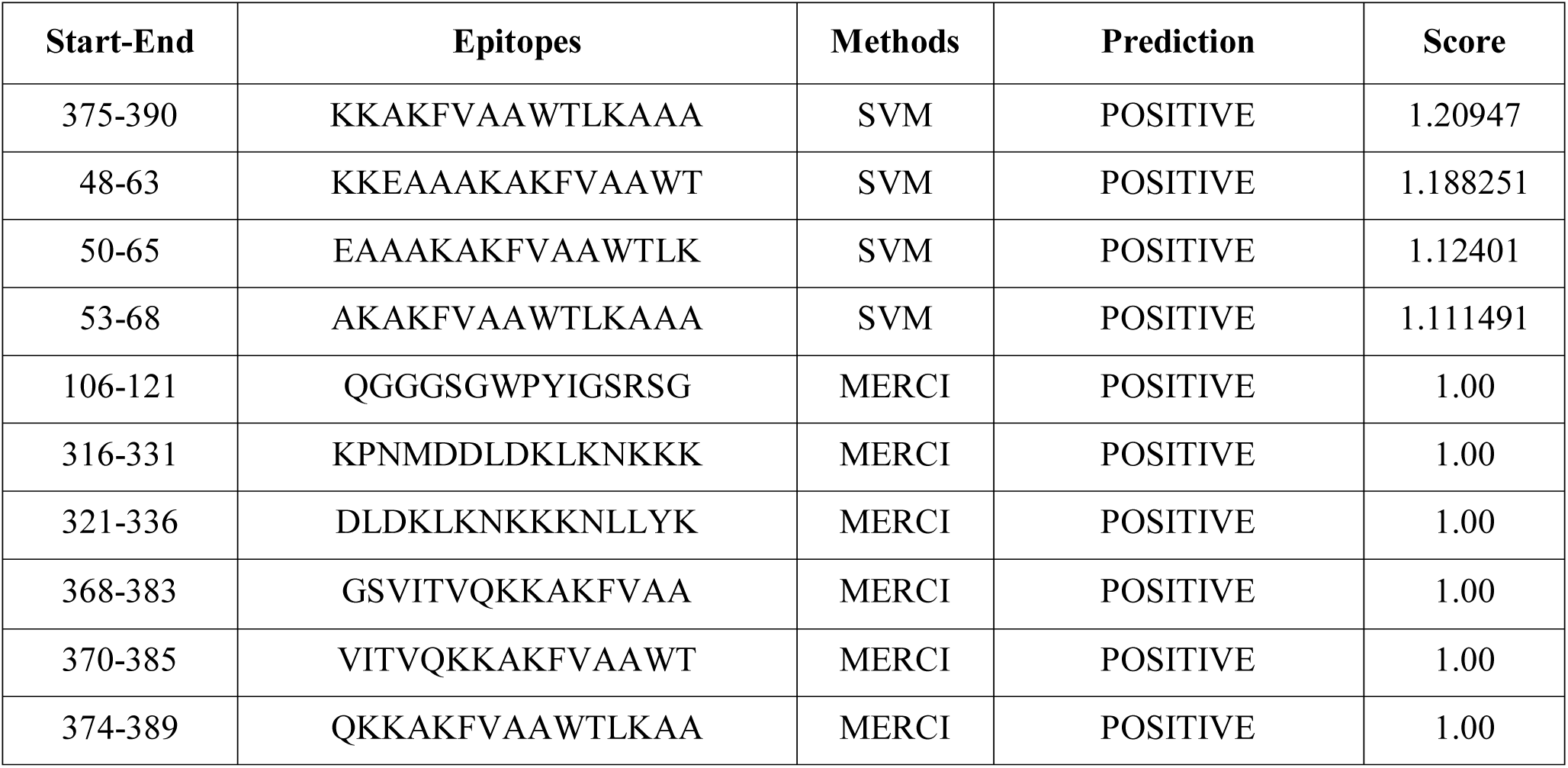
Predicted IFN-γ inducing epitopes from the proposed vaccine

### Protein-protein docking

The affinity between the constructed vaccines and HLA alleles were evaluated using molecular docking via Patchdock server. The server ranked the docked complexes based on Atomic Contact Energy (ACE), complementarity score and approximate interface area of the complex. Results revealed that construct V1 was superior in terms of free binding Energy (Table 10). Moreover, binding affinity of construct V1 with different human immune receptors was also demonstrated (Table 10). The lowest binding energy of the complexes indicated the highest binding affinity between receptor and vaccine construct.

**Table 10:** Docking scores of vaccine constructs with different HLA alleles i.e. HLA-DRB1*03:01 (1a6a), HLA-DRB5*01:01 (1h15), HLA-DRB1*01:01 (2fse), HLA-DRB3*01:01 (2q6w), HLA-DRB1*04:01 (2seb) and HLA-DRB3*02:02 (3c5j) and receptors i.e. TLR 3 (2a0z), MDA 5 (3b6e), α dystroglycan (5llk), RIG-1 (2qfd)

| *Vaccine construct* | *PDB IDs of HLA/receptors* | *Global Energy* | *Hydrogen bond energy* | *ACE* | *Score* | *Area* |
| --- | --- | --- | --- | --- | --- | --- |
| V1 | 1a6a | -2.65 | -3.20 | 5.18 | 16280 | 2111.50 |
|  | 1h15 | -6.40 | -2.75 | 7.08 | 17246 | 2552.90 |
|  | 2fse | -26.12 | -2.94 | 0.28 | 16062 | 2181.20 |
|  | 2q6w | -2.59 | -4.48 | 9.33 | 15968 | 2084.90 |
|  | 2seb | -28.54 | -3.71 | 5.51 | 17904 | 2308.00 |
|  | 3c5j | -28.29 | -5.12 | 9.40 | 16956 | 2397.50 |
| V2 | 1a6a | -0.55 | -2.61 | 12.31 | 16394 | 2349.80 |
|  | 1h15 | -0.55 | -4.21 | 10.79 | 17406 | 2402.80 |
|  | 2fse | 12.04 | -5.21 | 12.26 | 18306 | 2646.60 |
|  | 2q6w | -0.33 | -3.06 | 6.47 | 18262 | 2940.70 |
|  | 2seb | -5.52 | -0.82 | -1.77 | 21390 | 3942.60 |
|  | 3c5j | -22.27 | -3.87 | 9.04 | 16846 | 2299.50 |
| V3 | 1a6a | -0.23 | -4.64 | 10.85 | 17830 | 2324.60 |
|  | 1h15 | -16.48 | -3.48 | 15.41 | 19260 | 2658.10 |
|  | 2fse | -9.38 | -1.68 | 12.37 | 19720 | 2651.90 |
|  | 2q6w | -23.12 | -4.09 | 10.44 | 19724 | 2701.10 |
|  | 2seb | -19.67 | -0.52 | -3.87 | 18596 | 3097.60 |
|  | 3c5j | 1.78 | -5.71 | 12.71 | 18192 | 2430.00 |
| V1 | 2a0z | -10.82 | -3.34 | 2.33 | 14764 | 2172.70 |
|  | 3b6e | -3.91 | -3.33 | 16.69 | 15762 | 1889.60 |
|  | 5llk | -17.17 | -6.88 | 4.72 | 15410 | 1906.70 |
|  | 2qfd | -0.89 | -3.03 | 9.80 | 14636 | 1953.30 |

### Molecular dynamics simulation

The high ranked complex between vaccine molecules and TLR-3 were selected for analysis through Normal mode analysis (NMA). NMA was performed to describe the stability of proteins and large scale mobility. Results showed that the mobility of vaccine protein V1 and α-dystroglycan were oriented towards each other. Probable deformabilty of the complex was indicated by hinges in the chain due to distortion of the individual residues (Fig. 7A). The B-factor values which was equivalent to RMS inferred via NMA (Fig. 7B). Eigenvalue for the complex demonstrated was 2.2023e−06 (Fig. 7C). Colored bars showed the individual (red) and cummulative (green) variances which were inversely related to eigenvalue (Fig. 7D). The indicated coupling between Different interactions between residues were revealed by covariance matrix i.e. correlated, uncorrelated and anti-correlated motions indicated by red, white and blue colors respectively (Fig. 7E). The elastic network model (Fig. 7F). It identified the pairs of atoms those were connected via springs.

**Fig. 7:**
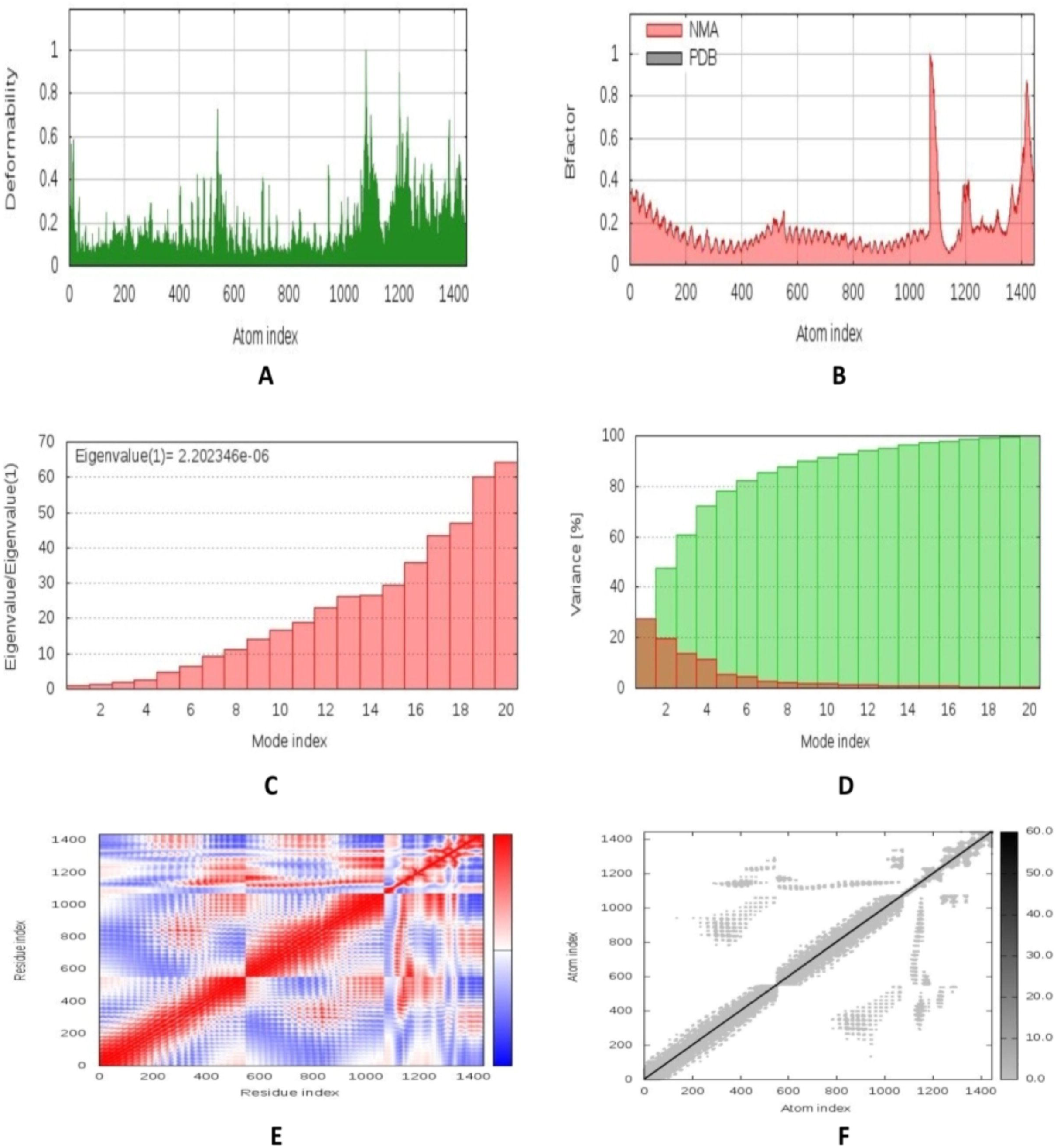
Molecular dynamics simulation of vaccine protein V1 and TLR-3 complex. Stability of the protein-protein complex was investigated through deformability (A), B-factor (B), eigenvalue (C), variance (D), covariance (E) and elastic network (F) analysis.

### Codon adaptation and in silico cloning

The Codon Adaptation Index for the optimized codons of construct V1 was 0.969 determined via JCAT server. The GC content of the adapted codons was also significant (50.93%). An insert of 1194 bp was obtained which lacked restriction sites for BglII and BglI ensuring, thus ensuring safety for cloning purpose. The codons were inserted into pET28a(+) vector along with BglII and BglI restriction sites and a clone of 4777 base pair was produced (Fig. 8).

**Fig. 8:**
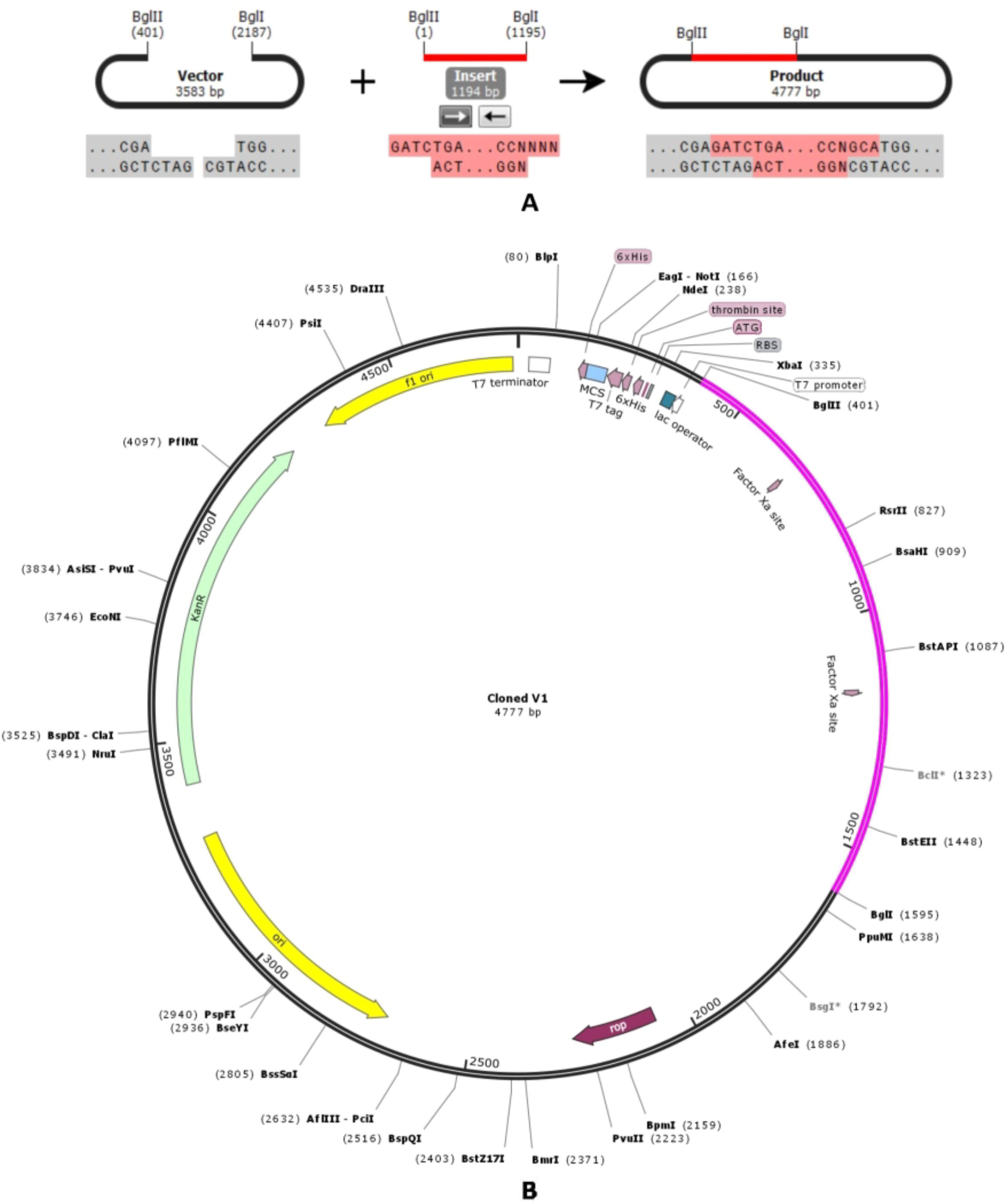
Restriction digestion (A) and *in silico* cloning (B) of the gene sequence of final construct V1 into pET28a(+) expression vector. Target sequence was inserted between BglII (401) and BGlI (2187) indicated in violate color.

## Discussion

Arenaviruses exhibit a catastrophic potential to destroy the public health scenario in several regions of the world. Few members of Arenavidae family pose a credible bioterrorism threat (Hauer et al., 2002), while six of them, including LASV and LCMV are classified as Category A agents by the National Institute of Allergy and Infectious Diseases (Borio et al., 2002; Charrel and de Lamballerie, 2003). Despite the significance of arenaviruses in public health and biodefense readiness, to date there are no vaccines approved by the Food and Drug Administration (FDA) (Brisse and Ly, 2019; Buchmeier et al 2007; Cheng et al., 2015). Current anti-arenavirus therapy is partially effective and requires an early and intravenous administration of nucleoside analog ribavirin that may cause significant side effects (Bausch et al., 2010; Moreno et al., 2011). Therefore, there is an unmet need to develop both safe and protective vaccines to combat pathogenic arenavirus infections in humans.

The live attenuated Candid#1 strain of Junin virus has been shown to be an effective vaccine against Argentine Hemorrhagic Fever (Ambrosio et al., 2011; Maiztegui et a., 1998). Researchers also found that Junin Virus Vaccine Antibodies are effective against Machupo Virus as well (Clark et al., 2011). However, the preventive measures to combat infections caused by LASV, LCMV, Lujo and Machupo virus has not yet gained considerable success. Although, ML29, a live attenuated vaccine has been shown to provide effective protection *in vivo* against Lassa virus (Carrion et al., 2007; Lukashevich et al., 2008), the mechanism for ML29 attenuation remains unknown (Olschlager and Flatz, 2013). Therefore, the incorporation of a limited number of additional mutations into the ML29 genome may result in viruses with enhanced virulence (Greenbaum et al., 2012). In the present study attempts were taken to develop a vaccine candidate using the most antigenic viral proteins of Arenaviridae family that could elicit broad spectrum immunity in the host. To the best of our knowledge, similar genome based screening and reverse vaccinology approach have not yet been explored to design an arenavirus vaccine.

Nucleoprotein (NP) specific CD8+ T cells play a major role in virus control and immune stimulation in the host (Schildknecht et al., 2008). Meulen and coworkers (2000) found that Lassa fever survivors had strong CD4+ T cell responses against LASV Nucleoprotein (ter Meulen et al., 2000). In another study, a single inoculation of a plasmid encoding full-length Lassa nucleoprotein induced CD8(+) T cell responses in mice model and protected against LCMV (Rodriguez-Carreno et al., 2005). All these findings suggest that anti-NP response at an early stage effectively controls and contributes to cross-protective immunity against arenavirus infections. In the present study, the superiority of Nucleoproteins of LASV, LCMV, Lujo and Guanarito virus in terms of antigenicity score were also revealed (Supplementary File 1). Hence, a multivalent vaccine strategy was implemented to protect against nucleoprotein antigens from different arenavirus species. To ensure protective response against a longer range of virus strains for a longer period, the candidate epitopes must remain in the highly conserved region. Thus, only the conserved sequences from each of the viral proteins (Table 2) were used for epitope prediction, which we believe, would generate more acceptable and universally effective vaccine constructs. Determination of potent T cell epitopes is a pivotal step during vaccine design, since T-cells play the key role in antibody production through antigen presentation (Amorim et al., 2016). Both MHC-I (CTL) and MHC-II restricted (HTL) epitopes were predicted using IEDB T cell epitope prediction tools and top candidates were screened based on transmembrane topology screening, antigenicity scoring, allergenicity pattern, toxicity profile and conservancy analysis (Table 3). The proposed epitopes showed a high cumulative population coverage in most of the geographic regions including 88% and 86% MHC-I coverage in South America and Africa respectively (Fig. 2). The top epitopes were highly conserved among different viral strains ranging from 40% to 100% (Table 4). In the next step, the predicted CTL and HTL epitopes were checked for their ability to bind with MHC-I and MHC-II alleles. Results showed that all the predicted epitopes were efficient binders of corresponding MHC allele (Table 5). B-cell epitopes boost neutralizing-antibody responses in the host. To achieve protective immunity against arenavirus infection, B-cell (BCL) epitopes were identified using three different algorithms (i.e. Linear Epitope, Surface Accessibility, Antigenicity) from IEDB (Table 6). Moreover, our results revealed that among the top epitopes, T cell epitope GWPYIGSRS were conserved in Argentine mammarenavirus (Junin virus) and Brazilian mammarenavirus (Sabia virus), while B cell epitope NLLYKICLSG were conserved in Bolivian mammarenavirus (Machupo virus) and Brazilian mammarenavirus (Sabia virus), indicating the possibility of final vaccine constructs to confer broad range immunity in the host against the arenaviruses.

The screened CTL, HTL and BCL epitopes were combined using respective linker to design the final vaccine constructs. Three different constructs (i.e. V1, V2, V3) were generated using 3 distinct linkers and investigated in terms of safety and efficacy (Table 7). Vaccine protein V1 was identified superior considering the immunogenicity score, allergenicity, hydrophobicity and other physicochemical properties. To strengthen our prediction, all three constructs were subjected to 3D modeling and interactions between designed vaccine constructs with different HLA molecules (i.e. HLA-DRB1*03:01, HLA-DRB5*01:01, HLA-DRB1*01:01, HLA-DRB3*01:01, HLA-DRB1*04:01 and HLA-DRB3*02:02) were determined (Table 7). Again construct V1 was found to be best in terms of free binding energy. IFN-γ is the signature cytokine of both the innate and adaptive immune systems with ability to provok antiviral immune responses and protection against reinfection. The release of IFN-γ enhances the magnitude of antiviral cytotoxic T lymphocytes (CTLs) responses and aids in production of neutralizing IgG (Nosrati et al., 2019). Construct V1 was screened to identify such IFN-γ producing epitopes which showed positive results (Table 9). Moreover, docking analysis was also performed to explore the binding affinity of construct V1 and different human immune-receptors (TLR 3, MDA 5, α dystroglycan, RIG-1) to evaluate the efficacy of used adjuvants (Table 10). α-dystroglycan (α-DG), a peripheral membrane protein, acts as an anchor between the submembranous cytoskeleton and the extracellular matrix which is widely expressed in most cells (Durbeej et al., 1998). It has been identified as a cellular receptor for LASV, certain strains of LCMV and the New World arenaviruses (Kunz et al., 2005, Spiropoulou et al., 2002). Molecular dynamics study by iMOD server ensured the stability of V1-TLR3 complex at molecular level. Finally, the designed construct V1 was reverse transcribed and inserted within pET28a(+) vector for heterologous expression in *E. coli* srain K12.

The predicted results were based on different sequence analysis and various immune databases. Due to the encouraging findings of the study, we suggest further wet lab based analysis using model animals for the experimental validation of our predicted vaccine candidates. Moreover, novel antigen delivery systems such as Nano-delivery platforms could enhance the efficacy of the proposed vaccine (Hojo 2014; Trovato and Berardinis 2015).

## Conclusion

*In-silico* bioinformatics study could be considered as a promising strategy to accelerate vaccine development against highly pathogenic organisms. In the present study, such approach was employed to design a novel heterosubtypic peptide vaccine against the most deadly viruses of Arenaviridae family requiring urgent need for effective medications and preventive measures. The study suggests, the proposed vaccine could stimulate both humoral and cellular mediated immune responses and serve as a potential vaccine against arenaviruses. However, in vitro and in vivo immunological experiments are highly recommended to validate the efficacy of designed vaccine constructs.

## Funding information

This research did not receive any specific grant from funding agencies in the public, commercial, or not-for-profit sectors.

## Conflict of interest

The Authors declare that they have no conflicts of interest.

## Acknowledgments

Authors would like to acknowledge the Department of Microbial Biotechnology and Department of Pharmaceuticals and Industrial Biotechnology at Sylhet Agricultural University for the technical support of the project.

