## Supplementary file 1 for "Conglomeration of highly antigenic nucleoproteins to inaugurate a heterosubtypic next generation vaccine candidate against Arenaviridae family"

| ***Proteins*** | ***VaxiJen Score*** |
| --- | --- |
| Nucleocapsid protein | 0.4657 |
| Nucleoprotein | 0.4663* |
| Glycoprotein precursor | 0.5042 |
| L polymerase | 0.4317 |

1. **Lassa virus (LCMV)**

**2. LCMV**

| ***Proteins*** | ***VaxiJen Score Score Score*** |
| --- | --- |
| L protein | 0.4617 |
| hypothetical protein | 0.4181 |
| envelope glycoprotein | 0.4382 |
| glycoprotein precursor | 0.4300 |
| Nucleoprotein | 0.5274* |
| zinc binding protein | 0.3209 |
| RNA-dependent RNA polymerase | 0.4652 |

**3. Lujo virus**

| ***Proteins*** | ***VaxiJen Score*** |
| --- | --- |
| Nucleocapsid protein | 0.5378* |
| Glycoprotein precursor | 0.2945 |
| Z protein | 0.8235 |
| Nucleoprotein | 0.5378 |

**4. Guanarito virus**

| ***Proteins*** | ***VaxiJen Score*** |
| --- | --- |
| Nucleocapsid protein | 0.5979 |
| Nucleocapsid protein | 0.5182* |
| RNA dependent RNA polymerase | 0.3744 |
| Glycoprotein precursor | 0.4174 |
