## Supplementary file 2 for "Conglomeration of highly antigenic nucleoproteins to inaugurate a heterosubtypic next generation vaccine candidate against Arenaviridae family"

| **Epitope**   1. **Lassa Virus (LASV)** | **Start** | **End** | **length** | **Antigenic value** | **Topology** | **No. of HLA allele** |
| --- | --- | --- | --- | --- | --- | --- |
| NQFGTMPSL | 2 | 10 | 9 | 0.2169 | out | 81 |
| MPSLTLACL | 7 | 15 | 9 | 1.3792 | out | 81 |
| SLTLACLTK | 9 | 17 | 9 | 1.4398 | in | 81 |
| YNFSLGAAVK | 3 | 12 | 10 | 1.6750 | in | 27 |
| GTMPSLTLA | 5 | 13 | 9 | 0.6293 | out | 81 |
| SLGAAVKAGA | 6 | 15 | 10 | 1.1754 | out | 27 |
| IEGRPEDPV | 4 | 12 | 9 | 1.1593 | out | 81 |
| AAVKAGACM | 9 | 17 | 9 | 0.4505 | in | 81 |
| YNFSLGAAV | 3 | 11 | 9 | 1.5124 | out | 54 |
| CMLDGGNML | 16 | 24 | 9 | 0.2412 | out | 81 |
| QFGTMPSLTL | 3 | 12 | 10 | 0.7735 | out | 27 |
| FSLGAAVKA | 5 | 13 | 9 | 1.3262 | out | 81 |
| MLDGGNMLET | 17 | 26 | 10 | 0.2183 | out | 27 |
| GYNFSLGAA | 2 | 10 | 9 | 1.2890 | out | 54 |
| NFSLGAAVK | 4 | 12 | 9 | 1.8014 | out | 27 |
| AVKAGACML | 10 | 18 | 9 | 0.4308 | in | 54 |
| FGTMPSLTL | 4 | 12 | 9 | 0.5793 | out | 27 |
| LTLACLTKQG | 10 | 19 | 10 | 1.4239 | in | 27 |
| MLDGGNMLE | 17 | 25 | 9 | 0.2202 | out | 27 |
| TMPSLTLAC | 6 | 14 | 9 | 1.1353 | out | 27 |
| ACMLDGGNM | 15 | 23 | 9 | 0.4207 | in | 54 |
| LACLTKQGQV | 12 | 21 | 10 | 1.0326 | in | 27 |
| LGAAVKAGA | 7 | 15 | 9 | 1.1852 | in | 54 |
| SGYNFSLGA | 1 | 9 | 9 | 1.1560 | out | 27 |
| LTKQGQVDL | 15 | 23 | 9 | 1.1525 | in | 81 |
| PSLTLACLT | 8 | 16 | 9 | 1.5130 | out | 27 |
| MDIEGRPED | 2 | 10 | 9 | 1.6690 | in | 81 |
| KAGACMLDGG | 12 | 21 | 10 | 0.3462 | in | 27 |
| LTLACLTKQ | 10 | 18 | 9 | 1.6679 | in | 27 |
| KAGACMLDG | 12 | 20 | 9 | 0.3798 | in | 54 |
| TLACLTKQG | 11 | 19 | 9 | 1.1541 | in | 54 |
| SLGAAVKAG | 6 | 14 | 9 | 1.4140 | out | 27 |
| AGACMLDGGN | 13 | 22 | 10 | 0.3635 | out | 27 |
| GAAVKAGAC | 8 | 16 | 9 | 0.4245 | in | 27 |
| NNQFGTMPS | 1 | 9 | 9 | 0.5609 | out | 27 |
| QFGTMPSLT | 3 | 11 | 9 | 0.7616 | out | 27 |
| LACLTKQGQ | 12 | 20 | 9 | 1.1557 | in | 27 |
| KQGQVDLND | 17 | 25 | 9 | 1.5505 | in | 54 |
| WMDIEGRPE | 1 | 9 | 9 | 1.5164 | in | 27 |
| ACLTKQGQV | 13 | 21 | 9 | 0.6964 | in | 54 |
| VKAGACMLD | 11 | 19 | 9 | 0.5651 | in | 27 |
| AGACMLDGG | 13 | 21 | 9 | 0.3359 | out | 27 |
| CLTKQGQVD | 14 | 22 | 9 | 0.7456 | in | 27 |
| TKQGQVDLN | 16 | 24 | 9 | 1.5089 | in | 27 |
| GACMLDGGN | 14 | 22 | 9 | 0.3847 | out | 27 |
| EGRPEDPVE | 5 | 13 | 9 | 0.7206 | out | 27 |
| LDGGNMLET | 18 | 26 | 9 | 0.2706 | out | 27 |
| DIEGRPEDP | 3 | 11 | 9 | 1.2557 | out | 27 |

2. **LCMV**

| **Epitope** | **Start** | **End** | **length** | **Antigenic value** | **Topology** | **No. of HLA allele** |
| --- | --- | --- | --- | --- | --- | --- |
| EVSNVQRIMR | 9 | 18 | 10 | 0.0712 | in | 27 |
| SEVSNVQRI | 8 | 16 | 9 | 0.4379 | in | 81 |
| DFSEVSNVQR | 6 | 15 | 10 | 1.0062 | in | 27 |
| LLNGLDFSEV | 1 | 10 | 10 | 1.0539 | out | 27 |
| LTDLGLLYTV | 8 | 17 | 10 | 1.1600 | out | 27 |
| LLKDLMGGI | 2 | 10 | 9 | -0.2958 | out | 81 |
| AAVKAGAAL | 6 | 14 | 9 | 0.7473 | in | 81 |
| NQFGTMPSL | 7 | 15 | 9 | 0.2169 | out | 81 |
| EVSNVQRIM | 9 | 17 | 9 | 0.3017 | in | 81 |
| VSNVQRIMR | 10 | 18 | 9 | -0.1279 | in | 54 |
| LLNNQFGTM | 4 | 12 | 9 | 0.3449 | out | 81 |
| FSEVSNVQR | 7 | 15 | 9 | 0.6900 | in | 81 |
| ALTDLGLLY | 7 | 15 | 9 | 1.2301 | out | 81 |
| LYTVKYPNL | 14 | 22 | 9 | 0.7742 | Out | 54 |
| FGTMPSLTM | 9 | 17 | 9 | 0.5672 | out | 54 |
| GLDFSEVSNV | 4 | 13 | 10 | 1.5849 | out | 27 |
| DVVQALTDL | 3 | 11 | 9 | 0.1074 | out | 81 |
| SLGAAVKAGA | 3 | 12 | 10 | 1.1754 | out | 27 |
| SLLNNQFGT | 3 | 11 | 9 | 0.3276 | out | 81 |
| LDFSEVSNV | 5 | 13 | 9 | 1.7221 | out | 81 |
| LGLLYTVKY | 11 | 19 | 9 | 1.0988 | out | 81 |
| VQALTDLGL | 5 | 13 | 9 | 0.9793 | out | 81 |
| FSLGAAVKA | 2 | 10 | 9 | 1.3262 | out | 81 |
| LLYTVKYPN | 13 | 21 | 9 | 0.6658 | out | 81 |
| AVKAGAALL | 7 | 15 | 9 | 0.6775 | out | 54 |
| SNVQRIMRK | 11 | 19 | 9 | -0.5731 | in | 81 |
| NFSLGAAVK | 1 | 9 | 9 | 1.8014 | out | 54 |
| MLLKDLMGG | 1 | 9 | 9 | 0.0150 | out | 54 |
| DSSLLNNQF | 1 | 9 | 9 | 0.2953 | out | 54 |
| TDLGLLYTV | 9 | 17 | 9 | 1.2339 | out | 81 |
| DLGLLYTVK | 10 | 18 | 9 | 1.4806 | out | 81 |
| LGAAVKAGA | 4 | 12 | 9 | 1.1852 | in | 81 |
| SSLLNNQFG | 2 | 10 | 9 | 0.4632 | out | 81 |
| QALTDLGLL | 6 | 14 | 9 | 0.9791 | out | 81 |
| GAAVKAGAA | 5 | 13 | 9 | 0.8243 | in | 81 |
| LLNGLDFSE | 1 | 9 | 9 | 0.8326 | out | 54 |
| LNDVVQALT | 1 | 9 | 9 | 0.0427 | out | 54 |
| SLGAAVKAG | 3 | 11 | 9 | 1.4140 | out | 81 |
| LNGLDFSEV | 2 | 10 | 9 | 1.3833 | out | 81 |
| GLLYTVKYP | 12 | 20 | 9 | 0.2158 | out | 81 |
| DLMGGIDPN | 5 | 13 | 9 | 1.1459 | out | 54 |
| DFSEVSNVQ | 6 | 14 | 9 | 1.3277 | out | 81 |
| GLDFSEVSN | 4 | 12 | 9 | 1.6058 | out | 81 |
| NNQFGTMPS | 6 | 14 | 9 | 0.5609 | out | 81 |
| NVQRIMRKE | 12 | 20 | 9 | -0.2380 | in | 54 |
| QFGTMPSLT | 8 | 16 | 9 | 0.7616 | out | 81 |
| VVQALTDLG | 4 | 12 | 9 | 0.5147 | out | 81 |
| NGLDFSEVS | 3 | 11 | 9 | 1.6145 | out | 81 |
| NDVVQALTD | 2 | 10 | 9 | -0.2640 | out | 81 |
| KDLMGGIDP | 4 | 12 | 9 | 0.9073 | out | 81 |
| LNNQFGTMP | 5 | 13 | 9 | 0.7367 | out | 81 |
| LKDLMGGID | 3 | 11 | 9 | 0.5750 | out | 81 |

| **Epitope** | **Start** | **End** | **length** | **Antigenic value** | **Topology** | **No. of HLA allele** |
| --- | --- | --- | --- | --- | --- | --- |
| NQFGTMPSL | 15 | 23 | 9 | 0.2169 | out | 81 |
| LLNNQFGTM | 12 | 20 | 9 | 0.3449 | 0ut | 81 |
| VWDVKDSSLL | 4 | 13 | 10 | 1.4658 | out | 27 |
| SLLNNQFGT | 11 | 19 | 9 | 0.3276 | out | 54 |
| RVWDVKDSSL | 3 | 12 | 10 | 0.9679 | in | 27 |
| KDSSLLNNQF | 8 | 17 | 10 | 0.6510 | out | 27 |
| DSSLLNNQF | 9 | 17 | 9 | 0.2953 | out | 54 |
| SSLLNNQFG | 10 | 18 | 9 | 0.4632 | out | 27 |
| WDVKDSSLL | 5 | 13 | 9 | 1.7861 | out | 54 |
| RVWDVKDSS | 3 | 11 | 9 | 0.7777 | in | 54 |
| DVKDSSLLN | 6 | 14 | 9 | 1.2269 | out | 54 |
| VWDVKDSSL | 4 | 12 | 9 | 1.7406 | out | 27 |
| NNQFGTMPS | 14 | 22 | 9 | 0.5609 | out | 54 |
| QFGTMPSLT | 16 | 24 | 9 | 0.7616 | out | 27 |
| KDSSLLNNQ | 8 | 16 | 9 | 0.8111 | in | 54 |
| VVRVWDVKD | 1 | 9 | 9 | 0.5383 | in | 54 |
| VKDSSLLNN | 7 | 15 | 9 | 0.5330 | out | 27 |
| LNNQFGTMP | 13 | 21 | 9 | 0.7367 | out | 27 |
| VRVWDVKDS | 2 | 10 | 9 | 0.6103 | in | 27 |

**Table 03: Lujo virus**

**Table 04: Guanarito virus**

| **Epitope** | **Start** | **End** | **length** | **Antigenic value** | **Topology** | **No. of HLA allele** |
| --- | --- | --- | --- | --- | --- | --- |
| WPYIGSRSQI | 12 | 21 | 10 | 1.4280 | i | 27 |
| ESALNISGY | 21 | 59 | 9 | -0.0392 | o | 27 |
| KLFDIHGRK | 10 | 18 | 9 | 0.8775 | i | 27 |
| PTDPVELAV | 10 | 18 | 9 | 0.6297 | o | 27 |
| LTSLGLLYTV | 17 | 26 | 10 | 0.9627 | o | 27 |
| AAVKAGASL | 5 | 13 | 9 | 0.7024 | i | 27 |
| YPNLDDLEKL | 28 | 37 | 10 | 0.4322 | o | 27 |
| TSLGLLYTVK | 18 | 27 | 10 | 1.3006 | o | 27 |
| RMYMGNLTQ | 2 | 10 | 9 | 0.2116 | i | 27 |
| NQFGSMPAL | 18 | 26 | 9 | 0.2749 | o | 27 |
| ETMNNVVQA | 8 | 16 | 9 | -0.0011 | i | 27 |
| ETMNNVVQAL | 8 | 17 | 10 | -0.0584 | i | 27 |
| KLIADSLDF | 3 | 11 | 9 | 0.1593 | o | 27 |
| LEHDCLQII | 39 | 47 | 9 | 0.6442 | o | 27 |
| EGWPYIGSR | 10 | 18 | 9 | 1.1255 | o | 27 |
| RLWDVSDPSK | 6 | 15 | 10 | 0.2329 | o | 27 |
| YPNLDDLEK | 28 | 36 | 9 | 0.5575 | o | 27 |
| RAHNGVIVPK | 10 | 19 | 10 | 0.8194 | i | 27 |
| TMNNVVQAL | 9 | 17 | 9 | 0.0273 | i | 27 |
| KLNNQFGSM | 15 | 23 | 9 | 1.0028 | i | 27 |
| ALTSLGLLY | 16 | 24 | 9 | 1.0529 | o | 27 |
| MTVQGGETM | 2 | 10 | 9 | 0.7086 | i | 27 |
| LYTVKYPNL | 23 | 31 | 9 | 0.7742 | o | 27 |
| TSLGLLYTV | 18 | 26 | 9 | 1.0632 | o | 27 |
| SQLEKRAGIL | 11 | 20 | 10 | 1.1223 | i | 27 |
| PYIGSRSQI | 13 | 21 | 9 | 1.1749 | i | 27 |
| CLSGEGWPY | 6 | 14 | 9 | -0.1194 | o | 27 |
| SLIDGGNML | 12 | 20 | 9 | 0.1760 | o | 27 |
| YMGNLTQSQL | 4 | 13 | 10 | 0.6416 | i | 27 |
| SLSAAVKAGA | 2 | 11 | 10 | 0.9408 | i | 27 |
| LQIITKDESA | 44 | 53 | 10 | 0.5369 | i | 27 |
| SLGLLYTVK | 19 | 27 | 9 | 1.2711 | o | 27 |
| LTQSQLEKR | 8 | 16 | 9 | 0.7481 | i | 27 |
| IADSLDFTQ | 5 | 13 | 9 | 0.9931 | o | 27 |
| WPYIGSRSQ | 12 | 20 | 9 | 2.0118 | i | 27 |
| KLCLSGEGW | 4 | 12 | 9 | -0.3483 | o | 27 |
| GETMNNVVQ | 7 | 15 | 9 | -0.0346 | I | 27 |
| QLEKRAGIL | 12 | 20 | 9 | 1.1648 | i | 27 |
| CLSGEGWPYI | 6 | 15 | 10 | 0.0662 | o | 27 |
| FGSMPALTI | 20 | 28 | 9 | 0.5644 | o | 27 |
| SQLEKRAGI | 11 | 19 | 9 | 1.0958 | i | 27 |
| SAAVKAGASL | 4 | 13 | 10 | 0.6709 | i | 27 |
| DPSKLNNQF | 12 | 20 | 9 | 0.7301 | o | 27 |
| TVQGADDIKK | 1 | 10 | 10 | 0.5294 | i | 27 |
| QALTSLGLLY | 15 | 24 | 10 | 0.7498 | o | 27 |
| NVVQALTSL | 12 | 20 | 9 | 0.1214 | o | 27 |
| YKLCLSGEGW | 3 | 12 | 10 | -0.3456 | o | 27 |
| DESALNISGY | 50 | 59 | 10 | 0.0083 | o | 27 |
| IEGPPTDPV | 6 | 14 | 9 | -0.1253 | o | 27 |
| VWEKFGHLCR | 1 | 10 | 10 | 0.9698 | i | 27 |
| KKLFDIHGRK | 9 | 18 | 10 | 0.5948 | i | 27 |
| QFGSMPALTI | 19 | 28 | 10 | 0.8518 | o | 27 |
| LCLSGEGWPY | 5 | 14 | 10 | 0.0943 | o | 27 |
| LTSLGLLYT | 17 | 25 | 9 | 1.1555 | o | 27 |
| LLYTVKYPNL | 22 | 31 | 10 | 0.9144 | o | 27 |
| VQALTSLGL | 14 | 22 | 9 | 0.8826 | o | 27 |
| VQGGETMNNV | 4 | 13 | 10 | 0.8873 | i | 27 |
| ALTSLGLLYT | 16 | 25 | 10 | 1.0419 | o | 27 |
| TVQGADDIK | 1 | 9 | 9 | 1.0901 | i | 27 |
| MYMGNLTQS | 3 | 11 | 9 | 0.8213 | i | 27 |
| GASLIDGGNM | 10 | 19 | 10 | 0.4308 | o | 27 |
| LTLEHDCLQI | 37 | 46 | 10 | 1.1842 | o | 27 |
| VELAVFQPS | 14 | 22 | 9 | 0.0368 | o | 27 |
| NLTQSQLEKR | 7 | 16 | 10 | 0.7791 | i | 27 |
| QIITKDESAL | 45 | 54 | 10 | 0.2192 | i | 27 |
| LEHDCLQIIT | 39 | 48 | 10 | 0.4509 | i | 27 |
| AHNGVIVPK | 11 | 19 | 9 | 0.8676 | i | 27 |
| HLCRAHNGV | 7 | 15 | 9 | -0.3709 | I | 27 |
| PTDPVELAVF | 10 | 19 | 10 | 0.3349 | o | 27 |
| NLTQSQLEK | 7 | 15 | 9 | 0.0613 | i | 27 |
| RMYMGNLTQS | 2 | 11 | 10 | 0.2975 | i | 27 |
| RLWDVSDPS | 6 | 14 | 9 | 0.3925 | o | 27 |
| LGLLYTVKY | 20 | 28 | 9 | 1.0988 | o | 27 |
| MYMGNLTQSQ | 3 | 12 | 10 | 0.8186 | i | 27 |
| KLIADSLDFT | 3 | 12 | 10 | 0.5760 | o | 27 |
| LSAAVKAGA | 3 | 11 | 9 | 0.9406 | i | 27 |
| LSGEGWPYI | 7 | 15 | 9 | 0.0332 | o | 27 |
| SLGLLYTVKY | 19 | 28 | 10 | 1.1456 | o | 27 |
| AAVKAGASLI | 5 | 14 | 10 | 0.5233 | i | 27 |
| KKLFDIHGR | 9 | 17 | 9 | 0.7113 | i | 27 |
| DIHGRKDLK | 13 | 21 | 9 | 0.9886 | i | 27 |
| AVKAGASLI | 6 | 14 | 9 | 0.5622 | i | 27 |
| SKLNNQFGSM | 14 | 23 | 10 | 0.9144 | i | 27 |
| WEKFGHLCR | 2 | 10 | 9 | 1.3383 | i | 27 |
| AKLIADSLDF | 2 | 11 | 10 | 0.2510 | o | 27 |
| LYTVKYPNLD | 23 | 32 | 10 | 0.8842 | o | 27 |
| LIADSLDFT | 4 | 12 | 9 | 0.9245 | o | 27 |
| LLYKLCLSG | 1 | 9 | 9 | -0.0598 | o | 27 |
| GEGWPYIGS | 9 | 17 | 9 | 0.3011 | o | 27 |
| LLYTVKYPN | 22 | 30 | 9 | 0.6658 | o | 27 |
| GETMNNVVQA | 7 | 16 | 10 | 0.1341 | i | 27 |
| SDPSKLNNQF | 11 | 20 | 10 | 0.5514 | o | 27 |
| DIHGRKDLKL | 13 | 22 | 10 | 1.2226 | i | 27 |
| DESALNISG | 50 | 58 | 9 | 0.0854 | o | 27 |
| GNLTQSQLEK | 6 | 15 | 10 | 0.1514 | i | 27 |
| DAKLIADSL | 1 | 9 | 9 | -0.0872 | o | 27 |
| AGASLIDGGN | 9 | 18 | 10 | 0.3803 | o | 27 |
| DPVELAVFQ | 12 | 20 | 9 | 0.4073 | o | 27 |
| FSLSAAVKA | 1 | 9 | 9 | 0.9878 | o | 27 |
| NLDDLEKLTL | 30 | 39 | 10 | 0.4825 | o | 27 |
| MTVQGGETMN | 2 | 11 | 10 | 0.9092 | i | 27 |
| IADSLDFTQV | 5 | 14 | 10 | 1.2088 | o | 27 |
| VQGADDIKK | 2 | 10 | 9 | 0.3789 | i | 27 |
| TDPVELAVF | 11 | 19 | 9 | 0.4049 | o | 27 |
| RAHNGVIVP | 10 | 18 | 9 | 0.5120 | i | 27 |
| AHNGVIVPKK | 11 | 20 | 10 | 1.1635 | i | 27 |
| QLEKRAGILR | 12 | 21 | 10 | 0.0092 | i | 27 |
| HLCRAHNGVI | 7 | 16 | 10 | -0.3134 | i | 27 |
| HGRKDLKLV | 15 | 23 | 9 | 1.6492 | i | 27 |
| ESALNISGYN | 51 | 60 | 10 | 0.1921 | o | 27 |
| TMNNVVQALT | 9 | 18 | 10 | 0.0281 | i | 27 |
| LWDVSDPSKL | 7 | 16 | 10 | 0.1638 | o | 27 |
| LLYKLCLSGE | 1 | 10 | 10 | 0.1430 | o | 27 |
| KDLKLVDVR | 18 | 26 | 9 | 2.1171 | i | 27 |
| YMGNLTQSQ | 4 | 12 | 9 | 0.6991 | i | 27 |
| IKKLFDIHGR | 8 | 17 | 10 | 0.3401 | i | 27 |
| MDIEGPPTD | 4 | 12 | 9 | 0.0857 | o | 27 |
| LEKRAGILR | 13 | 21 | 9 | -0.1505 | i | 27 |
| GSMPALTIA | 21 | 29 | 9 | 0.4807 | o | 27 |
| CMTVQGGETM | 1 | 10 | 10 | 0.6659 | i | 27 |
| LSAAVKAGAS | 3 | 12 | 10 | 0.7742 | i | 27 |
| QALTSLGLL | 15 | 23 | 9 | 0.8575 | o | 27 |
| ASLIDGGNM | 11 | 19 | 9 | 0.4038 | o | 27 |
| NVVQALTSLG | 12 | 21 | 10 | 0.4067 | o | 27 |
| PPTDPVELAV | 9 | 18 | 10 | 0.5214 | o | 27 |
| HNGVIVPKK | 12 | 20 | 9 | 1.3491 | i | 27 |
| LWDVSDPSK | 7 | 15 | 9 | 0.1430 | o | 27 |
| SLIDGGNMLE | 12 | 21 | 10 | 0.2879 | o | 27 |
| SALNISGYN | 52 | 60 | 9 | 0.3415 | i | 27 |
| VQALTSLGLL | 14 | 23 | 10 | 0.7725 | o | 27 |
| GEGWPYIGSR | 9 | 18 | 10 | 0.8026 | o | 27 |
| DLKLVDVRL | 19 | 27 | 9 | 2.0917 | i | 27 |
| FSLSAAVKAG | 1 | 10 | 10 | 1.2441 | o | 27 |
| HDCLQIITK | 41 | 49 | 9 | 0.3641 | i | 27 |
| TLEHDCLQII | 38 | 47 | 10 | 0.6578 | i | 27 |
| LAVFQPSSG | 16 | 24 | 9 | -0.3385 | o | 27 |
| EHDCLQIITK | 40 | 49 | 10 | 0.6430 | i | 27 |
| GLLYTVKYPN | 21 | 30 | 10 | 0.4543 | o | 27 |
| VSDPSKLNN | 10 | 18 | 9 | 0.3275 | o | 27 |
| NQFGSMPALT | 18 | 27 | 10 | 0.3607 | o | 27 |
| GGVVRLWDV | 2 | 10 | 9 | 0.0055 | o | 27 |
| MNNVVQALT | 10 | 18 | 9 | -0.0275 | i | 27 |
| MGNLTQSQL | 5 | 13 | 9 | 0.7469 | i | 27 |
| LSGEGWPYIG | 7 | 16 | 10 | 0.3404 | o | 27 |
| GPPTDPVEL | 8 | 16 | 9 | 0.3338 | o | 27 |
| KLVDVRLTG | 21 | 29 | 9 | 1.0107 | i | 27 |
| LCRAHNGVI | 8 | 16 | 9 | -0.2472 | i | 27 |
| LIDGGNMLET | 13 | 22 | 10 | 0.2195 | o | 27 |
| WMDIEGPPT | 3 | 11 | 9 | 0.2938 | o | 27 |
| SLSAAVKAG | 2 | 10 | 9 | 1.1477 | O | 27 |
| IITKDESAL | 46 | 54 | 9 | 0.3026 | i | 27 |
| TKDESALNI | 48 | 56 | 9 | 0.7431 | i | 27 |
| KLVDVRLTGE | 21 | 30 | 10 | 0.9072 | i | 27 |
| LYKLCLSGE | 2 | 10 | 9 | 0.1107 | o | 27 |
| VWEKFGHLC | 1 | 9 | 9 | 1.3953 | o | 27 |
| KYPNLDDLE | 27 | 35 | 9 | 1.4366 | o | 27 |
| ASLIDGGNML | 11 | 20 | 10 | 0.1916 | o | 27 |
| LIDGGNMLE | 13 | 21 | 9 | 0.2213 | o | 27 |
| LFDIHGRKDL | 11 | 20 | 10 | 0.6773 | o | 27 |
| GADDIKKLF | 4 | 12 | 9 | -0.3684 | i | 27 |
| ADDIKKLFDI | 5 | 14 | 10 | -0.0463 | i | 27 |
| FDIHGRKDLK | 12 | 21 | 10 | 1.2782 | i | 27 |
| KFGHLCRAH | 4 | 12 | 9 | 1.9055 | i | 27 |
| KAGASLIDG | 8 | 16 | 9 | 0.2350 | o | 27 |
| KFGHLCRAHN | 4 | 13 | 10 | 1.7709 | i | 27 |
| PSKLNNQFG | 13 | 21 | 9 | 0.9649 | o | 27 |
| KLFDIHGRKD | 10 | 19 | 10 | 0.8576 | i | 27 |
| NNQFGSMPA | 17 | 25 | 9 | 0.7739 | o | 27 |
| NNQFGSMPAL | 17 | 26 | 10 | 0.6923 | o | 27 |
| ADSLDFTQV | 6 | 14 | 9 | 1.4384 | o | 27 |
| KYPNLDDLEK | 27 | 36 | 10 | 0.8144 | o | 27 |
| DIEGPPTDPV | 5 | 14 | 10 | -0.0128 | o | 27 |
| IWMDIEGPPT | 2 | 11 | 10 | -0.1434 | o | 27 |
| EKLTLEHDCL | 35 | 44 | 10 | 0.6102 | o | 27 |
| ITKDESALNI | 47 | 56 | 10 | 0.3392 | i | 27 |
| KDLKLVDVRL | 18 | 27 | 10 | 1.9240 | i | 27 |
| VVQALTSLGL | 13 | 22 | 10 | 0.7699 | o | 27 |
| QIITKDESA | 45 | 53 | 9 | 0.1986 | i | 27 |
| LIADSLDFTQ | 4 | 13 | 10 | 0.6903 | o | 27 |
| IWMDIEGPP | 2 | 10 | 9 | -0.2953 | o | 27 |
| FGSMPALTIA | 20 | 29 | 10 | 0.6323 | o | 27 |
| WDVSDPSKL | 8 | 16 | 9 | 0.4045 | o | 27 |
| KAGASLIDGG | 8 | 17 | 10 | 0.3902 | o | 27 |
| IKKLFDIHG | 8 | 16 | 9 | -0.3743 | i | 27 |
| SAAVKAGAS | 4 | 12 | 9 | 0.6168 | i | 27 |
| LEKLTLEHDC | 34 | 43 | 10 | 0.2223 | i | 27 |
| VELAVFQPSS | 14 | 23 | 10 | 0.0924 | o | 27 |
| LCRAHNGVIV | 8 | 17 | 10 | 0.0352 | i | 27 |
| GGETMNNVVQ | 6 | 15 | 10 | 0.1993 | i | 27 |
| CMTVQGGET | 1 | 9 | 9 | 0.7926 | i | 27 |
| TVKYPNLDDL | 25 | 34 | 10 | 1.1379 | o | 27 |
| ELAVFQPSS | 15 | 23 | 9 | -0.0793 | o | 27 |
| WEKFGHLCRA | 2 | 11 | 10 | 1.1609 | i | 27 |
| ELAVFQPSSG | 15 | 24 | 10 | -0.2459 | o | 27 |
| DIKKLFDIH | 7 | 15 | 9 | 0.3829 | i | 27 |
| PPTDPVELA | 9 | 17 | 9 | 0.6855 | o | 27 |
| RKDLKLVDVR | 17 | 26 | 10 | 1.9287 | i | 27 |
| TLEHDCLQI | 38 | 46 | 10 | 0.9472 | i | 27 |
| KLNNQFGSMP | 15 | 24 | 10 | 0.9731 | o | 27 |
| EKFGHLCRA | 3 | 11 | 9 | 1.3901 | i | 27 |
| EGWPYIGSRS | 10 | 19 | 10 | 1.1750 | o | 27 |
| FDIHGRKDL | 12 | 20 | 9 | 0.8652 | i | 27 |
| YTVKYPNLD | 24 | 32 | 9 | 1.0699 | o | 27 |
| NNVVQALTSL | 11 | 20 | 10 | 0.0100 | i | 27 |
| QGADDIKKLF | 3 | 12 | 10 | -0.2740 | i | 27 |
| DDLEKLTLE | 32 | 40 | 9 | -0.1408 | o | 27 |
| ADSLDFTQVS | 6 | 15 | 10 | 1.5975 | o | 27 |
| HNGVIVPKKK | 12 | 21 | 10 | 1.6874 | i | 27 |
| VKYPNLDDL | 26 | 34 | 9 | 1.2431 | o | 27 |
| KLTLEHDCLQ | 36 | 45 | 10 | 0.6417 | i | 27 |
| DPSKLNNQFG | 12 | 21 | 10 | 0.8367 | o | 27 |
| QFGSMPALT | 16 | 27 | 9 | 0.8106 | o | 27 |
| LYKLCLSGEG | 2 | 11 | 10 | -0.2781 | o | 27 |
| LDDLEKLTL | 31 | 39 | 9 | 0.1609 | o | 27 |
| IEGPPTDPVE | 6 | 15 | 10 | -0.1112 | o | 27 |
| NGVIVPKKK | 13 | 21 | 9 | 1.8982 | i | 27 |
| IDGGNMLETI | 14 | 23 | 10 | 0.1795 | o | 27 |
| LEKLTLEHD | 34 | 42 | 9 | -0.0427 | o | 27 |
| KLTLEHDCL | 36 | 44 | 9 | 0.8165 | o | 27 |
| ITKDESALN | 47 | 55 | 9 | 0.0540 | i | 27 |
| DGGNMLETI | 15 | 23 | 9 | -0.1055 | o | 27 |
| DDIKKLFDI | 6 | 14 | 9 | -0.2574 | i | 27 |
| CRAHNGVIV | 9 | 17 | 9 | 0.1075 | i | 27 |
| MDIEGPPTDP | 4 | 13 | 10 | 0.0943 | o | 27 |
| IHGRKDLKLV | 14 | 23 | 10 | 1.5649 | i | 27 |
| GLLYTVKYP | 21 | 29 | 9 | 0.2158 | o | 27 |
| IITKDESALN | 46 | 55 | 9 | 0.3168 | i | 27 |
| KDESALNISG | 49 | 58 | 10 | 0.3197 | i | 27 |
| LTLEHDCLQ | 37 | 45 | 9 | 0.8932 | o | 27 |
| LKLVDVRLTG | 20 | 29 | 10 | 1.4348 | o | 27 |
| GHLCRAHNGV | 6 | 15 | 10 | -0.0900 | i | 27 |
| DPVELAVFQP | 12 | 21 | 10 | 0.3546 | o | 27 |
| QSQLEKRAGI | 10 | 19 | 10 | 0.7545 | i | 27 |
| NLDDLEKLT | 30 | 38 | 9 | 0.3755 | o | 27 |
| VVQALTSLG | 13 | 21 | 9 | 0.4605 | o | 27 |
| LTQSQLEKRA | 8 | 17 | 10 | 0.7229 | i | 27 |
| GVVRLWDVSD | 3 | 12 | 10 | 0.3262 | o | 27 |
| GPPTDPVELA | 8 | 17 | 10 | 0.5447 | o | 27 |
| IHGRKDLKL | 14 | 22 | 9 | 1.4787 | i | 27 |
| QGGETMNNV | 5 | 13 | 9 | 0.7181 | i | 27 |
| LVDVRLTGE | 22 | 30 | 9 | 1.3388 | o | 27 |
| DLEKLTLEH | 33 | 41 | 9 | 0.3359 | o | 27 |
| LNNQFGSMPA | 16 | 25 | 10 | 0.7284 | o | 27 |
| LQIITKDES | 44 | 52 | 9 | 0.6582 | i | 27 |
| VKAGASLIDG | 7 | 16 | 10 | 0.4654 | o | 27 |
| VVRLWDVSD | 4 | 12 | 9 | 0.4455 | o | 27 |
| TQSQLEKRAG | 9 | 18 | 10 | 0.7707 | i | 27 |
| GGVVRLWDVS | 2 | 11 | 10 | 0.2412 | o | 27 |
| PRMYMGNLTQ | 1 | 10 | 10 | 0.0902 | i | 27 |
| DVSDPSKLNN | 9 | 18 | 10 | 0.4591 | o | 27 |
| GWPYIGSRSQ | 11 | 20 | 10 | 1.5639 | i | 27 |
| AVKAGASLID | 6 | 15 | 10 | 0.7599 | i | 27 |
| LGLLYTVKYP | 20 | 29 | 10 | 0.7348 | o | 27 |
| PNLDDLEKL | 29 | 37 | 9 | 0.4095 | o | 27 |
| YTVKYPNLDD | 24 | 33 | 10 | 1.0207 | o | 27 |
| MGNLTQSQLE | 5 | 14 | 10 | 0.8475 | i | 27 |
| CRAHNGVIVP | 9 | 18 | 10 | 0.1580 | i | 27 |
| TVQGGETMNN | 3 | 12 | 10 | 0.8547 | i | 27 |
| AGGVVRLWD | 1 | 9 | 9 | -0.1228 | o | 27 |
| VKYPNLDDLE | 26 | 35 | 10 | 1.2780 | o | 27 |
| DVSDPSKLN | 9 | 17 | 9 | 0.4953 | o | 27 |
| VQGADDIKKL | 2 | 11 | 10 | 0.2344 | i | 27 |
| AGGVVRLWDV | 1 | 10 | 10 | 0.1957 | o | 27 |
| TIWMDIEGP | 1 | 9 | 9 | -0.8149 | o | 27 |
| TVKYPNLDD | 25 | 33 | 9 | 1.0730 | o | 27 |
| EGPPTDPVEL | 7 | 16 | 10 | 0.3686 | o | 27 |
| MNNVVQALTS | 10 | 19 | 10 | -0.0481 | i | 27 |
| HGRKDLKLVD | 15 | 24 | 10 | 1.8349 | i | 27 |
| DAKLIADSLD | 1 | 10 | 10 | 0.0977 | o | 27 |
| DIKKLFDIHG | 7 | 16 | 10 | -0.0973 | i | 27 |
| FGHLCRAHNG | 5 | 14 | 10 | 0.4479 | i | 27 |
| WMDIEGPPTD | 3 | 12 | 10 | 0.2193 | o | 27 |
| GGETMNNVV | 6 | 14 | 9 | 0.3615 | i | 27 |
| TQSQLEKRA | 9 | 17 | 9 | 0.7365 | i | 27 |
| VRLWDVSDPS | 5 | 14 | 10 | 0.2926 | o | 27 |
| EKFGHLCRAH | 3 | 12 | 10 | 1.2447 | i | 27 |
| RKDLKLVDV | 17 | 25 | 19 | 2.1621 | i | 27 |
| EHDCLQIIT | 40 | 48 | 9 | 0.4831 | i | 27 |
| GWPYIGSRS | 11 | 19 | 9 | 1.5322 | o | 27 |
| DSLDFTQVS | 7 | 15 | 9 | 1.8576 | o | 27 |
| TVQGGETMN | 3 | 11 | 9 | 0.9281 | i | 27 |
| FGHLCRAHN | 5 | 13 | 9 | 0.9764 | i | 27 |
| VQGGETMNN | 4 | 12 | 9 | 0.9598 | i | 27 |
| TDPVELAVFQ | 11 | 20 | 10 | 0.3076 | o | 27 |
| DLKLVDVRLT | 19 | 28 | 10 | 2.4438 | i | 27 |
| KLCLSGEGWP | 4 | 13 | 10 | -0.1189 | o | 27 |
| GASLIDGGN | 10 | 18 | 9 | 0.3930 | o | 27 |
| YKLCLSGEG | 3 | 11 | 9 | -0.0536 | o | 27 |
| PSKLNNQFGS | 13 | 22 | 10 | 0.8133 | o | 27 |
| TIWMDIEGPP | 1 | 10 | 10 | -0.3783 | o | 27 |
| SGEGWPYIGS | 8 | 17 | 10 | 0.3105 | o | 27 |
| DCLQIITKD | 42 | 50 | 9 | 0.6622 | i | 27 |
| VKAGASLID | 7 | 15 | 9 | 0.8510 | i | 27 |
| GVVRLWDVS | 3 | 11 | 9 | 0.2145 | o | 27 |
| WDVSDPSKLN | 8 | 17 | 10 | 0.6364 | o | 27 |
| EKLTLEHDC | 35 | 43 | 9 | 0.4367 | o | 27 |
| KDESALNIS | 49 | 57 | 9 | 0.9535 | i | 27 |
| QGADDIKKL | 3 | 11 | 9 | 0.1669 | i | 27 |
| GADDIKKLFD | 4 | 13 | 10 | -0.2753 | i | 27 |
| VSDPSKLNNQ | 10 | 19 | 10 | 0.5424 | o | 27 |
| LKLVDVRLT | 20 | 28 | 9 | 2.0649 | i | 27 |
| CLQIITKDE | 43 | 51 | 9 | 1.0813 | i | 27 |
| PVELAVFQPS | 13 | 22 | 10 | 0.2450 | o | 27 |
| PRMYMGNLT | 1 | 9 | 9 | 0.2016 | i | 27 |
| LNNQFGSMP | 16 | 24 | 9 | 0.8733 | o | 27 |
| NNVVQALTS | 11 | 19 | 9 | -0.1588 | i | 27 |
| CLQIITKDES | 43 | 52 | 10 | 0.8666 | i | 27 |
| DDIKKLFDIH | 6 | 15 | 10 | -0.1333 | i | 27 |
| LCLSGEGWP | 5 | 13 | 9 | -0.0569 | o | 27 |
| QSQLEKRAG | 10 | 18 | 9 | 0.6431 | i | 27 |
| SKLNNQFGS | 14 | 22 | 9 | 0.8310 | i | 27 |
| GRKDLKLVDV | 16 | 25 | 10 | 2.3188 | i | 27 |
| AGASLIDGG | 9 | 17 | 9 | 0.3452 | o | 27 |
| VRLWDVSDP | 5 | 13 | 9 | 0.2852 | o | 27 |
| QGGETMNNVV | 5 | 14 | 10 | 0.4389 | i | 27 |
| EGPPTDPVE | 7 | 15 | 9 | 0.0701 | o | 27 |
| DDLEKLTLEH | 32 | 41 | 10 | 0.0284 | o | 27 |
| GHLCRAHNG | 6 | 14 | 9 | -0.1944 | i | 27 |
| VVRLWDVSDP | 4 | 13 | 10 | 0.1694 | o | 27 |
| AKLIADSLD | 2 | 10 | 9 | -0.2948 | o | 27 |
| SGEGWPYIG | 8 | 16 | 9 | 0.4220 | o | 27 |
| HDCLQIITKD | 41 | 50 | 10 | 0.4545 | I | 27 |
| GNLTQSQLE | 6 | 14 | 9 | 0.7856 | o | 27 |
| DLEKLTLEHD | 33 | 42 | 10 | 0.0619 | o | 27 |
| PVELAVFQP | 13 | 21 | 9 | 0.2762 | o | 27 |
| IDGGNMLET | 14 | 22 | 9 | 0.3037 | o | 27 |
| LFDIHGRKD | 11 | 19 | 9 | 0.8378 | i | 27 |
| SDPSKLNNQ | 11 | 19 | 9 | 0.5868 | o | 27 |
| TKDESALNIS | 48 | 57 | 10 | 0.9212 | i | 27 |
| LDDLEKLTLE | 31 | 40 | 10 | 0.2265 | o | 27 |
| GRKDLKLVD | 16 | 24 | 9 | 2.3137 | i | 27 |
| ADDIKKLFD | 5 | 13 | 9 | -0.7417 | i | 27 |
| DCLQIITKDE | 42 | 51 | 10 | 0.7493 | i | 27 |
| DIEGPPTDP | 5 | 13 | 9 | -0.0431 | o | 27 |
| PNLDDLEKLT | 39 | 38 | 10 | 0.4135 | o | 27 |
