## Supplementary file 3 for "Conglomeration of highly antigenic nucleoproteins to inaugurate a heterosubtypic next generation vaccine candidate against Arenaviridae family"

1. **Lassa virus**

| **Epitope** | **Start** | **End** | **length** | **Antigenic value** | **Topology** | **No. of HLA allele** |
| --- | --- | --- | --- | --- | --- | --- |
| SGYNFSLGAAVKAGA | 1 | 15 | 15 | 1.0922 | out | 27 |
| GYNFSLGAAVKAGAC | 2 | 16 | 15 | 0.9536 | out | 27 |
| YNFSLGAAVKAGACM | 3 | 17 | 15 | 1.1719 | out | 27 |
| NFSLGAAVKAGACML | 4 | 18 | 15 | 1.1070 | out | 27 |
| FSLGAAVKAGACMLD | 5 | 19 | 15 | 0.9323 | out | 27 |
| NNQFGTMPSLTLACL | 1 | 15 | 15 | 1.0193 | out | 27 |
| NQFGTMPSLTLACLT | 2 | 16 | 15 | 0.8694 | out | 27 |
| QFGTMPSLTLACLTK | 3 | 17 | 15 | 0.9671 | out | 27 |
| FGTMPSLTLACLTKQ | 4 | 18 | 15 | 1.0122 | out | 27 |
| SLGAAVKAGACMLDG | 6 | 20 | 15 | 0.6925 | out | 27 |
| LGAAVKAGACMLDGG | 7 | 21 | 15 | 0.6404 | out | 27 |
| KAGACMLDGGNMLET | 12 | 26 | 15 | 0.3213 | out | 27 |
| VKAGACMLDGGNMLE | 11 | 25 | 15 | 0.4061 | out | 27 |
| AVKAGACMLDGGNML | 10 | 24 | 15 | 0.3412 | in | 27 |
| TMPSLTLACLTKQGQ | 6 | 20 | 15 | 1.1134 | out | 27 |
| MPSLTLACLTKQGQV | 7 | 21 | 15 | 1.0459 | out | 27 |
| PSLTLACLTKQGQVD | 8 | 22 | 15 | 1.1813 | out | 27 |
| SLTLACLTKQGQVDL | 9 | 23 | 15 | 1.3579 | out | 27 |
| LTLACLTKQGQVDLN | 10 | 24 | 15 | 1.4060 | out | 27 |
| TLACLTKQGQVDLND | 11 | 25 | 15 | 1.2790 | in | 27 |
| GTMPSLTLACLTKQG | 5 | 19 | 15 | 0.9712 | out | 27 |
| GAAVKAGACMLDGGN | 8 | 22 | 15 | 0.4672 | in | 27 |
| AAVKAGACMLDGGNM | 9 | 23 | 15 | 0.4531 | in | 27 |

2. **LCMV**

| **Epitope** | **Start** | **End** | **Length** | **Antigenic value** | **Topology** | **No. of HLA allele** |
| --- | --- | --- | --- | --- | --- | --- |
| NFSLGAAVKAGAALL | 1 | 15 | 15 | 1.2041 | out | 27 |
| SLLNNQFGTMPSLTM | 3 | 17 | 15 | 0.5471 | out | 27 |
| LTDLGLLYTVKYPNL | 8 | 22 | 15 | 1.1861 | out | 27 |
| LNDVVQALTDLGLLY | 1 | 15 | 15 | 0.5537 | out | 27 |
| ALTDLGLLYTVKYPN | 7 | 21 | 15 | 0.9756 | out | 27 |
| GLDFSEVSNVQRIMR | 4 | 18 | 15 | 0.7005 | in | 27 |
| LDFSEVSNVQRIMRK | 5 | 19 | 15 | 0.6827 | in | 27 |
| VVQALTDLGLLYTVK | 4 | 18 | 15 | 0.8009 | out | 27 |
| NDVVQALTDLGLLYT | 2 | 16 | 15 | 0.5214 | out | 27 |
| DFSEVSNVQRIMRKE | 6 | 20 | 15 | 0.5219 | in | 27 |
| DVVQALTDLGLLYTV | 3 | 17 | 15 | 0.5867 | out | 27 |
| VQALTDLGLLYTVKY | 5 | 19 | 15 | 0.8413 | out | 27 |
| QALTDLGLLYTVKYP | 6 | 20 | 15 | 0.6954 | out | 27 |
| NGLDFSEVSNVQRIM | 3 | 17 | 15 | 0.7633 | out | 27 |
| LNGLDFSEVSNVQRI | 2 | 16 | 15 | 0.9237 | out | 27 |
| LLNGLDFSEVSNVQR | 1 | 15 | 15 | 0.8913 | out | 27 |
| DSSLLNNQFGTMPSL | 1 | 15 | 15 | 0.3935 | out | 27 |
| SSLLNNQFGTMPSLT | 2 | 16 | 15 | 0.4797 | out | 27 |

3. **Lujo virus**

| **Epitope** | **Start** | **End** | **Length** | **Antigenic value** | **Topology** | **No. of HLA allele** |
| --- | --- | --- | --- | --- | --- | --- |
| VVRVWDVKDSSLLNN | 1 | 15 | 15 | 0.4941 | in | 27 |
| VRVWDVKDSSLLNNQ | 2 | 16 | 15 | 0.6717 | in | 27 |
| DVKDSSLLNNQFGTM | 6 | 20 | 15 | 0.8891 | out | 27 |
| VKDSSLLNNQFGTMP | 7 | 21 | 15 | 0.6619 | out | 27 |
| KDSSLLNNQFGTMPS | 8 | 22 | 15 | 0.5533 | out | 27 |
| DSSLLNNQFGTMPSL | 9 | 23 | 15 | 0.3935 | out | 27 |
| RVWDVKDSSLLNNQF | 3 | 17 | 15 | 0.7122 | in | 27 |
| VWDVKDSSLLNNQFG | 4 | 18 | 15 | 1.1337 | out | 27 |
| WDVKDSSLLNNQFGT | 5 | 19 | 15 | 1.1169 | out | 27 |
| SSLLNNQFGTMPSLT | 10 | 24 | 15 | 0.4797 | out | 27 |

**4. Guanarito virus**

| **Epitope** | **start** | **End** | **Length** | **Antigenic value** | **Topology** | **No. of MHA allele** |
| --- | --- | --- | --- | --- | --- | --- |
| FSLSAAVKAGASLID | 1 | 15 | 15 | 0.8734 | o | 27 |
| NNVVQALTSLGLLYT | 11 | 25 | 15 | 0.4583 | i | 27 |
| HDCLQIITKDESALN | 41 | 55 | 15 | 0.4039 | i | 27 |
| DCLQIITKDESALNI | 42 | 56 | 15 | 0.6442 | i | 27 |
| CLQIITKDESALNIS | 43 | 57 | 15 | 0.9136 | i | 27 |
| LQIITKDESALNISG | 44 | 58 | 15 | 0.4166 | i | 27 |
| MNNVVQALTSLGLLY | 10 | 24 | 15 | 0.4470 | i | 27 |
| TMNNVVQALTSLGLL | 9 | 23 | 15 | 0.4933 | i | 27 |
| SKLNNQFGSMPALTI | 14 | 28 | 15 | 0.7851 | o | 27 |
| KLNNQFGSMPALTIA | 15 | 29 | 15 | 0.8366 | o | 27 |
| ETMNNVVQALTSLGL | 8 | 22 | 15 | 0.4572 | i | 27 |
| NVVQALTSLGLLYTV | 12 | 26 | 15 | 0.5157 | o | 27 |
| LTSLGLLYTVKYPNL | 17 | 31 | 15 | 1.0659 | o | 27 |
| TSLGLLYTVKYPNLD | 18 | 32 | 15 | 1.1723 | o | 27 |
| SLGLLYTVKYPNLDD | 19 | 33 | 15 | 1.1124 | o | 27 |
| LGLLYTVKYPNLDDL | 20 | 34 | 15 | 1.1312 | o | 27 |
| RMYMGNLTQSQLEKR | 2 | 16 | 15 | 0.5057 | i | 27 |
| MYMGNLTQSQLEKRA | 3 | 17 | 15 | 0.7786 | I | 27 |
| DAKLIADSLDFTQVS | 1 | 15 | 15 | 0.9498 | o | 27 |
| VVQALTSLGLLYTVK | 13 | 27 | 15 | 0.7332 | o | 27 |
| KLTLEHDCLQIITKD | 36 | 50 | 15 | 0.6800 | i | 27 |
| EKLTLEHDCLQIITK | 35 | 49 | 15 | 0.5343 | o | 27 |
| LEKLTLEHDCLQIIT | 34 | 48 | 15 | 0.3281 | o | 27 |
| SLSAAVKAGASLIDG | 2 | 16 | 15 | 0.5424 | o | 27 |
| DLEKLTLEHDCLQII | 33 | 47 | 15 | 0.4508 | o | 27 |
| LSAAVKAGASLIDGG | 3 | 17 | 15 | 0.5832 | o | 27 |
| ALTSLGLLYTVKYPN | 16 | 30 | 15 | 0.8800 | o | 27 |
| LTLEHDCLQIITKDE | 37 | 51 | 15 | 0.8608 | o | 27 |
| GETMNNVVQALTSLG | 7 | 21 | 15 | 0.3133 | i | 27 |
| VQALTSLGLLYTVKY | 14 | 28 | 15 | 0.7736 | o | 27 |
| EHDCLQIITKDESAL | 40 | 54 | 15 | 0.5629 | i | 27 |
| LEHDCLQIITKDESA | 39 | 53 | 15 | 0.5491 | o | 27 |
| TLEHDCLQIITKDES | 38 | 52 | 15 | 0.6102 | o | 27 |
| AGGVVRLWDVSDPSK | 1 | 15 | 15 | 0.1889 | o | 27 |
| GGVVRLWDVSDPSKL | 2 | 16 | 15 | 0.1082 | o | 27 |
| GVVRLWDVSDPSKLN | 3 | 17 | 15 | 0.2330 | o | 27 |
| VVRLWDVSDPSKLNN | 4 | 18 | 15 | 0.2586 | o | 27 |
| VRLWDVSDPSKLNNQ | 5 | 19 | 15 | 0.4618 | o | 27 |
| EKFGHLCRAHNGVIV | 3 | 17 | 15 | 0.7474 | i | 27 |
| YMGNLTQSQLEKRAG | 4 | 18 | 15 | 0.7489 | i | 27 |
| PRMYMGNLTQSQLEK | 1 | 15 | 15 | 0.0429 | i | 27 |
| QIITKDESALNISGY | 45 | 59 | 15 | 0.1521 | i | 27 |
| VWEKFGHLCRAHNGV | 1 | 15 | 15 | 0.5447 | o | 27 |
| WEKFGHLCRAHNGVI | 2 | 16 | 15 | 0.6232 | i | 27 |
| KFGHLCRAHNGVIVP | 4 | 18 | 15 | 0.9719 | i | 27 |
| FGHLCRAHNGVIVPK | 5 | 19 | 15 | 0.6489 | o | 27 |
| QALTSLGLLYTVKYP | 15 | 29 | 15 | 0.6268 | o | 27 |
| GHLCRAHNGVIVPKK | 6 | 20 | 15 | 0.5582 | i | 27 |
| CLSGEGWPYIGSRSQ | 6 | 20 | 15 | 0.7924 | o | 27 |
| LSGEGWPYIGSRSQI | 7 | 21 | 15 | 0.5885 | o | 27 |
| IHGRKDLKLVDVRLT | 14 | 28 | 15 | 1.8532 | i | 27 |
| HGRKDLKLVDVRLTG | 15 | 29 | 15 | 1.5423 | i | 27 |
| GRKDLKLVDVRLTGE | 16 | 30 | 15 | 1.6152 | i | 27 |
| SAAVKAGASLIDGGN | 4 | 18 | 15 | 0.4960 | o | 27 |
| GPPTDPVELAVFQPS | 8 | 22 | 15 | 0.1438 | o | 27 |
| PPTDPVELAVFQPSS | 9 | 23 | 15 | 0.1856 | o | 27 |
| PTDPVELAVFQPSSG | 10 | 24 | 15 | 0.1002 | o | 27 |
| GGETMNNVVQALTSL | 6 | 20 | 15 | 0.2434 | i | 27 |
| QGGETMNNVVQALTS | 5 | 19 | 15 | 0.2417 | i | 27 |
| AAVKAGASLIDGGNM | 5 | 19 | 15 | 0.5343 | o | 27 |
| LLYTVKYPNLDDLEK | 22 | 36 | 15 | 0.7160 | o | 27 |
| GLLYTVKYPNLDDLE | 21 | 35 | 15 | 0.9362 | o | 27 |
| LLYKLCLSGEGWPYI | 1 | 15 | 15 | -0.0320 | o | 27 |
| HLCRAHNGVIVPKKK | 7 | 21 | 15 | 0.7629 | i | 27 |
| IITKDESALNISGYN | 46 | 60 | 15 | 0.3004 | i | 27 |
| AVKAGASLIDGGNML | 6 | 20 | 15 | 0.4224 | o | 27 |
| PSKLNNQFGSMPALT | 13 | 27 | 15 | 0.7533 | o | 27 |
| TIWMDIEGPPTDPVE | 1 | 15 | 15 | -0.1387 | o | 27 |
| IWMDIEGPPTDPVEL | 2 | 16 | 15 | 0.1198 | o | 27 |
| GNLTQSQLEKRAGIL | 6 | 20 | 15 | 0.8500 | i | 27 |
| LYKLCLSGEGWPYIG | 2 | 16 | 15 | 0.1136 | o | 27 |
| TVQGADDIKKLFDIH | 1 | 15 | 15 | 0.4550 | i | 27 |
| VQGGETMNNVVQALT | 4 | 18 | 15 | 0.4208 | i | 27 |
| NLTQSQLEKRAGILR | 7 | 21 | 15 | 0.2725 | i | 27 |
| AGASLIDGGNMLETI | 9 | 23 | 15 | 0.2491 | o | 27 |
| TVQGGETMNNVVQAL | 3 | 17 | 15 | 0.4504 | i | 27 |
| TVKYPNLDDLEKLTL | 25 | 39 | 15 | 0.6551 | o | 27 |
| VKYPNLDDLEKLTLE | 26 | 40 | 15 | 0.6730 | o | 27 |
| KYPNLDDLEKLTLEH | 27 | 41 | 15 | 0.7097 | o | 27 |
| YPNLDDLEKLTLEHD | 28 | 42 | 15 | 0.4008 | o | 27 |
| KAGASLIDGGNMLET | 8 | 22 | 15 | 0.3450 | o | 27 |
| MGNLTQSQLEKRAGI | 5 | 19 | 15 | 0.8611 | i | 27 |
| VKAGASLIDGGNMLE | 7 | 21 | 15 | 0.4873 | o | 27 |
| LYTVKYPNLDDLEKL | 23 | 37 | 15 | 0.5427 | o | 27 |
| YTVKYPNLDDLEKLT | 24 | 38 | 15 | 0.6279 | o | 27 |
| VQGADDIKKLFDIHG | 2 | 16 | 15 | 0.0605 | i | 27 |
| YKLCLSGEGWPYIGS | 3 | 17 | 15 | 0.2333 | o | 27 |
| DDIKKLFDIHGRKDL | 6 | 20 | 15 | 0.1193 | i | 27 |
| IKKLFDIHGRKDLKL | 8 | 22 | 15 | 0.7460 | i | 27 |
| ADDIKKLFDIHGRKD | 5 | 19 | 15 | 0.2316 | i | 27 |
| DDLEKLTLEHDCLQI | 32 | 46 | 15 | 0.3782 | o | 27 |
| PNLDDLEKLTLEHDC | 29 | 43 | 15 | 0.4729 | o | 27 |
| EGPPTDPVELAVFQP | 7 | 21 | 15 | 0.1675 | o | 27 |
| DIKKLFDIHGRKDLK | 7 | 21 | 15 | 0.6942 | i | 27 |
| KKLFDIHGRKDLKLV | 9 | 23 | 15 | 1.0615 | i | 27 |
| KLFDIHGRKDLKLVD | 10 | 24 | 15 | 1.3894 | i | 27 |
| LFDIHGRKDLKLVDV | 11 | 25 | 15 | 1.5311 | o | 27 |
| FDIHGRKDLKLVDVR | 12 | 26 | 15 | 1.6867 | i | 27 |
| NLDDLEKLTLEHDCL | 30 | 44 | 15 | 0.5626 | o | 27 |
| LDDLEKLTLEHDCLQ | 31 | 45 | 15 | 0.3259 | o | 27 |
| MTVQGGETMNNVVQA | 2 | 16 | 15 | 0.5628 | i | 27 |
| CMTVQGGETMNNVVQ | 1 | 15 | 15 | 0.5230 | i | 27 |
| GADDIKKLFDIHGRK | 4 | 18 | 15 | 0.3385 | i | 27 |
| RLWDVSDPSKLNNQF | 6 | 20 | 15 | 0.4921 | o | 27 |
| QGADDIKKLFDIHGR | 3 | 17 | 15 | 0.3823 | i | 27 |
| DIHGRKDLKLVDVRL | 13 | 27 | 15 | 1.4867 | i | 27 |
| IEGPPTDPVELAVFQ | 6 | 20 | 15 | 0.0555 | o | 27 |
| DIEGPPTDPVELAVF | 5 | 19 | 15 | 0.1581 | o | 27 |
| LWDVSDPSKLNNQFG | 7 | 21 | 15 | 0.5974 | o | 27 |
| SDPSKLNNQFGSMPA | 11 | 25 | 15 | 0.6093 | o | 27 |
| DPSKLNNQFGSMPAL | 12 | 26 | 15 | 0.6868 | o | 27 |
| VSDPSKLNNQFGSMP | 10 | 24 | 15 | 0.6320 | o | 27 |
| WMDIEGPPTDPVELA | 3 | 17 | 15 | 0.5047 | o | 27 |
| MDIEGPPTDPVELAV | 4 | 18 | 15 | 0.3816 | o | 27 |
| KLCLSGEGWPYIGSR | 4 | 18 | 15 | 0.5621 | o | 27 |
| WDVSDPSKLNNQFGS | 8 | 22 | 15 | 0.6995 | o | 27 |
| DVSDPSKLNNQFGSM | 9 | 23 | 15 | 0.6918 | o | 27 |
| LCLSGEGWPYIGSRS | 5 | 19 | 15 | 0.7691 | o | 27 |
