## Supplementary file 4 for "Conglomeration of highly antigenic nucleoproteins to inaugurate a heterosubtypic next generation vaccine candidate against Arenaviridae family"

***Allergenicity pattern, toxicity profile and conservancy analysis of top CTL epitopes (MHC-1 peptides)***

1. **Lassa Virus**

| Epitope | Allertop | AllergenFp | Allermatch | AllergenOnline | Toxinpred | Conservancy (identity <= 100%) |
| --- | --- | --- | --- | --- | --- | --- |
| IEGRPEDPV | Non-allergen | Non-allergen | Non-allergen | Non-allergen | Non-toxin | 100.00% (100/100) |
| NFSLGAAVK | Non-allergen | Non-allergen | Non-allergen | Non-allergen | Non-toxin | 100.00% (100/100) |
| PSLTLACLT | Allergen | Non-allergen | Non-allergen | Non-allergen | Non-toxin | 100.00% (100/100) |
| YNFSLGAAV | Non-allergen | Allergen | Non-allergen | Non-allergen | Non-toxin | 100.00% (100/100) |
| SLGAAVKAG | Non-allergen | Allergen | Allergen | Non-allergen | Non-toxin | 100.00% (100/100) |
| MPSLTLACL | Non-allergen | Non-allergen | Non-allergen | Non-allergen | Non-toxin | 97.00% (97/100) |
| FSLGAAVKA | Allergen | Allergen | Non-allergen | Non-allergen | Non-toxin | 100.00% (100/100) |
| GYNFSLGAA | Non-allergen | Non-allergen | Non-allergen | Non-allergen | Non-toxin | 100.00% (100/100) |
| DIEGRPEDP | Allergen | Non-allergen | Non-allergen | Non-allergen | Non-toxin | 100.00% (100/100) |
| SLGAAVKAGA | Non-allergen | Allergen | Allergen | Non-allergen | Non-toxin | 100.00% (100/100) |
| TMPSLTLAC | Allergen | Allergen | Non-allergen | Non-allergen | Non-toxin | 97.00% (97/100) |
| QFGTMPSLTL | Non-allergen | Allergen | Non-allergen | Non-allergen | Non-toxin | 97.00% (97/100) |
| EGRPEDPVE | Non-allergen | Non-allergen | Non-allergen | Non-allergen | Non-toxin | 100.00% (100/100) |
| GTMPSLTLA | Non-allergen | Allergen | Non-allergen | Non-allergen | Non-toxin | 97.00% (97/100) |
| FGTMPSLTL | Non-allergen | Allergen | Non-allergen | Non-allergen | Non-toxin | 97.00% (97/100) |

2. **LCMV**

| Epitope | Allertop | AllergenFp | Allermatch | AllergenOnline | Toxinpred | Conservancy (identity <= 100%) |
| --- | --- | --- | --- | --- | --- | --- |
| NFSLGAAVK | Non-allergen | Non-allergen | Non-allergen | Non-allergen | Non-toxin | 67.00% (67/100) |
| LDFSEVSNV | Non-allergen | Non-allergen | Non-allergen | Non-allergen | Non-toxin | 91.00% (91/100) |
| NGLDFSEVS | Non-allergen | Non-allergen | Non-allergen | Non-allergen | Non-toxin | 40.00% (40/100) |
| GLDFSEVSN | Non-allergen | Non-allergen | Non-allergen | Non-allergen | Non-toxin | 81.00% (81/100) |
| DLGLLYTVK | Allergen | Non-allergen | Non-allergen | Non-allergen | Non-toxin | 41.00% (41/100) |
| SLGAAVKAG | Non-allergen | Allergen | Allergen | Non-allergen | Non-toxin | 69.00% (69/100) |
| LNGLDFSEV | Non-allergen | Non-allergen | Non-allergen | Non-allergen | Non-toxin | 40.00% (40/100) |
| DFSEVSNVQ | Non-allergen | Allergen | Non-allergen | Non-allergen | Non-toxin | 91.00% (91/100) |
| FSLGAAVKA | Allergen | Allergen | Non-allergen | Non-allergen | Non-toxin | 69.00% (69/100) |
| TDLGLLYTV | Allergen | Allergen | Non-allergen | Non-allergen | Non-toxin | 40.00% (40/100) |
| ALTDLGLLY | Non-allergen | Non-allergen | Non-allergen | Non-allergen | Non-toxin | 40.00% (40/100) |
| SLGAAVKAGA | Non-allergen | Allergen | Allergen | Non-allergen | Non-toxin | 69.00% (69/100) |
| LTDLGLLYTV | Allergen | Non-allergen | Non-allergen | Non-allergen | Non-toxin | 40.00% (40/100) |
| DLMGGIDPN | Allergen | Allergen | Non-allergen | Non-allergen | Non-toxin | 33.00% (33/100) |
| LGLLYTVKY | Allergen | Non-allergen | Non-allergen | Non-allergen | Non-toxin | 41.00% (41/100) |
| LLNGLDFSEV | Non-allergen | Non-allergen | Non-allergen | Non-allergen | Non-toxin | 40.00% (40/100) |

3. **Lujo Virus**

| Epitope | Allertop v.2 | AllergenFp v.1 | Allermatch | AllergenOnline | Toxinpred | Conservancy (identity <= 100%) |
| --- | --- | --- | --- | --- | --- | --- |
| WDVKDSSLL | Allergen | Allergen | Non-allergen | Non-allergen | Non-toxin | 100.00% (4/4) |
| VWDVKDSSL | Allergen | Non-allergen | Non-allergen | Non-allergen | Non-toxin | 100.00% (4/4) |
| VWDVKDSSLL | Allergen | Non-allergen | Non-allergen | Non-allergen | Non-toxin | 100.00% (4/4) |
| DVKDSSLLN | Allergen | Allergen | Non-allergen | Non-allergen | Non-toxin | 100.00% (4/4) |
| RVWDVKDSSL | Allergen | Non-allergen | Non-allergen | Non-allergen | Non-toxin | 100.00% (4/4) |
| KDSSLLNNQ | Non-allergen | Allergen | Non-allergen | Non-allergen | Non-toxin | 100.00% (4/4) |
| RVWDVKDSS | Allergen | Non-allergen | Non-allergen | Non-allergen | Non-toxin | 100.00% (4/4) |
| QFGTMPSLT | Allergen | Allergen | Non-allergen | Non-allergen | Non-toxin | 100.00% (4/4) |
| LNNQFGTMP | Allergen | Allergen | Non-allergen | Non-allergen | Non-toxin | 100.00% (4/4) |
| KDSSLLNNQF | Non-allergen | Allergen | Non-allergen | Non-allergen | Non-toxin | 100.00% (4/4) |
| VRVWDVKDS | Allergen | Allergen | Non-allergen | Non-allergen | Non-toxin | 100.00% (4/4) |
| NNQFGTMPS | Allergen | Non-allergen | Non-allergen | Non-allergen | Non-toxin | 100.00% (4/4) |
| VVRVWDVKD | Allergen | Non-allergen | Non-allergen | Non-allergen | Non-toxin | 100.00% (4/4) |
| VKDSSLLNN | Allergen | Allergen | Non-allergen | Non-allergen | Non-toxin | 100.00% (4/4) |
| SSLLNNQFG | Non-allergen | Allergen | Non-allergen | Non-allergen | Non-toxin | 100.00% (4/4) |

4. **Guanarito Virus**

| Epitope | Alertop | AllergenFp | Allermatch | AllergenOnline | Toxinpred | Conservancy (identity <= 100%) |
| --- | --- | --- | --- | --- | --- | --- |
| DSLDFTQVS | Non-allergen | Non-allergen | Non-allergen | Non-allergen | Non-allergen toxin | 24.00% (24/100) |
| ADSLDFTQVS | Non-allergen | Non-allergen | Non-allergen | Non-allergen | Non-allergen toxin | 24.00% (24/100) |
| GWPYIGSRS | allergen | Non-allergen | Non-allergen | Non-allergen | Non-allergen toxin | 99.00% (99/100) |
| LKLVDVRLTG | allergen | Non-allergen | Non-allergen | Non-allergen | Non-allergen toxin | 24.00% (24/100) |
| ADSLDFTQV | Non-allergen | allergen | Non-allergen | Non-allergen | Non-allergen toxin | 24.00% (24/100) |
| KYPNLDDLE | Non-allergen | allergen | Non-allergen | Non-allergen | Non-allergen toxin | 26.00% (26/100) |
| VWEKFGHLC | allergen | Non-allergen | Non-allergen | Non-allergen | Non-allergen toxin | 29.00% (29/100) |
| LVDVRLTGE | Non-allergen | Non-allergen | Non-allergen | Non-allergen | Non-allergen toxin | 24.00% (24/100) |
| TSLGLLYTVK | allergen | allergen | Non-allergen | Non-allergen | Non-allergen toxin | 78.00% (78/100) |
| SLGLLYTVK | allergen | Non-allergen | Non-allergen | Non-allergen | Non-allergen toxin | 79.00% (79/100) |
| FSLSAAVKAG | allergen | allergen | Non-allergen | Non-allergen | Non-allergen toxin | 70.00% (70/100) |
| VKYPNLDDL | allergen | allergen | Non-allergen | Non-allergen | Non-allergen toxin | 28.00% (28/100) |
| IADSLDFTQV | Non-allergen | allergen | Non-allergen | Non-allergen | Non-allergen toxin | 24.00% (24/100) |
| LTLEHDCLQI | allergen | Non-allergen | Non-allergen | Non-allergen | Non-allergen toxin | 22.00% (22/100) |
