## Supplementary file 5 for "Conglomeration of highly antigenic nucleoproteins to inaugurate a heterosubtypic next generation vaccine candidate against Arenaviridae family"

***Allergenicity pattern, toxicity profile and conservancy analysis of top HTL epitopes (MHC-II peptides)***

1. **Lassa Virus**

| Epitope | Allertop | AllergenFp | Allermatch | AllergenOnline | Toxinpred | Conservancy (identity <= 100%) |
| --- | --- | --- | --- | --- | --- | --- |
| PSLTLACLTKQGQVD | Allergen | Allergen | Non-allergen | Non-allergen | Non-toxin | 100.00% (100/100) |
| NFSLGAAVKAGACML | Allergen | Non-allergen | Allergen | Non-allergen | Non-toxin | 100.00% (100/100) |
| SGYNFSLGAAVKAGA | Allergen | Non-allergen | Allergen | Non-allergen | Non-toxin | 100.00% (100/100) |
| NNQFGTMPSLTLACL | Non-allergen | Non-allergen | Non-allergen | Non-allergen | Non-toxin | 91.00% (91/100) |
| FSLGAAVKAGACMLD | Allergen | Non-allergen | Allergeb | Non-allergen | Non-toxin | 100.00% (100/100) |
| LTLACLTKQGQVDLN | Allergen | Non-allergen | Non-allergen | Non-allergen | Non-toxin | 100.00% (100/100) |
| SLTLACLTKQGQVDL | Allergen | Non-allergen | Non-allergen | Non-allergen | Non-toxin | 100.00% (100/100) |
| YNFSLGAAVKAGACM | Non-allergen | Non-allergen | Allergen | Non-allergen | Non-toxin | 100.00% (100/100) |
| TMPSLTLACLTKQGQ | Non-allergen | Non-allergen | Non-allergen | Non-allergen | Non-toxin | 97.00% (97/100) |
| MPSLTLACLTKQGQV | Non-allergen | Non-allergen | Non-allergen | Non-allergen | Non-toxin | 97.00% (97/100) |
| FGTMPSLTLACLTKQ | Non-allergen | Non-allergen | Non-allergen | Non-allergen | Non-toxin | 97.00% (97/100) |
| GTMPSLTLACLTKQG | Non-allergen | Non-allergen | Non-allergen | Non-allergen | Non-toxin | 97.00% (97/100) |
| QFGTMPSLTLACLTK | Non-allergen | Non-allergen | Non-allergen | Non-allergen | Non-toxin | 97.00% (97/100) |
| GYNFSLGAAVKAGAC | Allergen | Non-allergen | Allergen | Non-allergen | Non-toxin | 100.00% (100/100) |
| NQFGTMPSLTLACLT | Non-allergen | Non-allergen | Non-allergen | Non-allergen | Non-toxin | 97.00% (97/100) |

2. **LCMV**

| Epitope | Allertop | AllergenFp | Allermatch | AllergenOnline | Toxinpred | Conservancy (identity <= 100%) |
| --- | --- | --- | --- | --- | --- | --- |
| NFSLGAAVKAGAALL | Non-allergen | Non-allergen | Allergen | Non-allergen | Non-toxin | 34.00% (34/100) |
| LTDLGLLYTVKYPNL | Allergen | Non-allergen | Non-allergen | Non-allergen | Non-toxin | 40.00% (40/100) |
| ALTDLGLLYTVKYPN | Non-allergen | Non-allergen | Non-allergen | Non-allergen | Non-toxin | 40.00% (40/100) |
| LNGLDFSEVSNVQRI | Non-allergen | Non-allergen | Non-allergen | Non-allergen | Non-toxin | 40.00% (40/100) |
| LLNGLDFSEVSNVQR | Non-allergen | Non-allergen | Non-allergen | Non-allergen | Non-toxin | 40.00% (40/100) |
| VQALTDLGLLYTVKY | Non-allergen | Non-allergen | Non-allergen | Non-allergen | Non-toxin | 40.00% (40/100) |
| VVQALTDLGLLYTVK | Allergen | Allergen | Non-allergen | Non-allergen | Non-toxin | 40.00% (40/100) |
| NGLDFSEVSNVQRIM | Non-allergen | Non-allergen | Non-allergen | Non-allergen | Non-toxin | 40.00% (40/100) |
| QALTDLGLLYTVKYP | Non-allergen | Non-allergen | Non-allergen | Non-allergen | Non-toxin | 40.00% (40/100) |
| DVVQALTDLGLLYTV | Non-allergen | Non-allergen | Non-allergen | Non-allergen | Non-toxin | 40.00% (40/100) |
| LNDVVQALTDLGLLY | Non-allergen | Non-allergen | Non-allergen | Non-allergen | Non-toxin | 40.00% (40/100) |
| SLLNNQFGTMPSLTM | Allergen | Non-allergen | Non-allergen | Non-allergen | Non-toxin | 43.00% (43/100) |
| NDVVQALTDLGLLYT | Non-allergen | Allergen | Non-allergen | Non-allergen | Non-toxin | 40.00% (40/100) |
| SSLLNNQFGTMPSLT | Allergen | Non-allergen | Non-allergen | Non-allergen | Non-toxin | 45.00% (45/100) |

3. **Lujo Virus**

| Epitope | Allertop | AllergenFp | Allermatch | AllergenOnline | Toxinpred | Conservancy (identity <= 100%) |
| --- | --- | --- | --- | --- | --- | --- |
| VWDVKDSSLLNNQFG | Non-allergen | Non-allergen | Non-allergen | Non-allergen | Non-toxin | 4.00% (4/100) |
| WDVKDSSLLNNQFGT | Non-allergen | Non-allergen | Non-allergen | Non-allergen | Non-toxin | 4.00% (4/100) |
| DVKDSSLLNNQFGTM | Non-allergen | Non-allergen | Non-allergen | Non-allergen | Non-toxin | 4.00% (4/100) |
| RVWDVKDSSLLNNQF | Non-allergen | Allergen | Non-allergen | Non-allergen | Non-toxin | 4.00% (4/100) |
| VRVWDVKDSSLLNNQ | Non-allergen | Allergen | Non-allergen | Non-allergen | Non-toxin | 4.00% (4/100) |
| VKDSSLLNNQFGTMP | Non-allergen | Allergen | Non-allergen | Non-allergen | Non-toxin | 4.00% (4/100) |
| KDSSLLNNQFGTMPS | Non-allergen | Non-allergen | Non-allergen | Non-allergen | Non-toxin | 4.00% (4/100) |
| VVRVWDVKDSSLLNN | Non-allergen | Non-allergen | Non-allergen | Non-allergen | Non-toxin | 4.00% (4/100) |
| SSLLNNQFGTMPSLT | Allergen | Non-allergen | Non-allergen | Non-allergen | Non-toxin | 4.00% (4/100) |
| DSSLLNNQFGTMPSL | Allergen | Non-allergen | Non-allergen | Non-allergen | Non-toxin | 4.00% (4/100) |

4. **Guanarito Virus**

| Epitope | Allertop | AllergenFp | Allermatch | Allergen Online | Toxipred | Conservancy (identity <= 100%) |
| --- | --- | --- | --- | --- | --- | --- |
| LFDIHGRKDLKLVDV | allergen | Non-allergen | Non-allergen | Non-allergen | Non-toxin | 24.00% (24/100) |
| TSLGLLYTVKYPNLD | alllergen | Non-allergen | Non-allergen | Non-allergen | Non-toxin | 26.00% (26/100) |
| LGLLYTVKYPNLDDL | Non-allergen | Non-allergen | Non-allergen | Non-allergen | Non-toxin | 28.00% (28/100) |
| SLGLLYTVKYPNLDD | Non-allergen | Non-allergen | Non-allergen | Non-allergen | Non-toxin | 26.00% (26/100) |
| LTSLGLLYTVKYPNL | Non-allergen | Non-allergen | Non-allergen | Non-allergen | Non-toxin | 74.00% (74/100) |
| DAKLIADSLDFTQVS | Non-allergen | Non-allergen | Non-allergen | Non-allergen | Non-toxin | 24.00% (24/100) |
| GLLYTVKYPNLDDLE | allergen | Non-allergen | Non-allergen | Non-allergen | Non-toxin | 25.00% (25/100) |
| ALTSLGLLYTVKYPN | allergen | Non-allergen | Non-allergen | Non-allergen | Non-toxin | 71.00% (71/100) |
| FSLSAAVKAGASLID | allergen | Non-allergen | allergen | Non-allergen | Non-toxin | 23.00% (23/100) |
| LTLEHDCLQIITKDE | Non-allergen | Non-allergen | allergen | Non-allergen | Non-toxin | 22.00% (22/100) |
| KLNNQFGSMPALTIA | allergen | Non-allergen | Non-allergen | Non-allergen | Non-toxin | 48.00% (48/100) |
| CLSGEGWPYIGSRSQ | allergen | Non-allergen | Non-allergen | Non-allergen | Non-toxin | 36.00% (36/100) |
| SKLNNQFGSMPALTI | allergen | allergen | Non-allergen | Non-allergen | Non-toxin | 47.00% (47/100) |
| MYMGNLTQSQLEKRA | Non-allergen | Non-allergen | Non-allergen | Non-allergen | Non-toxin | 22.00% (22/100) |
| VQALTSLGLLYTVKY | allergen | allergen | Non-allergen | Non-allergen | Non-toxin | 48.00% (48/100) |
