## Supplementary file 6 for "Conglomeration of highly antigenic nucleoproteins to inaugurate a heterosubtypic next generation vaccine candidate against Arenaviridae family"

| **Number** | **Start-End** | **Epitopes** | **Methods** | **Prediction** | **Score** |
| --- | --- | --- | --- | --- | --- |
| 1 | 106-121 | [QGGGSGWPYIGSRSG](http://crdd.osdd.net/raghava/ifnepitope/pep_design.php?sequence=QGGGSGWPYIGSRSG&method=hybrid&model=main) | MERCI | POSITIVE | 1 |
| 2 | 316-331 | [KPNMDDLDKLKNKKK](http://crdd.osdd.net/raghava/ifnepitope/pep_design.php?sequence=KPNMDDLDKLKNKKK&method=hybrid&model=main) | MERCI | POSITIVE | 1 |
| 3 | 317-332 | [PNMDDLDKLKNKKKN](http://crdd.osdd.net/raghava/ifnepitope/pep_design.php?sequence=PNMDDLDKLKNKKKN&method=hybrid&model=main) | MERCI | POSITIVE | 1 |
| 4 | 318-333 | [NMDDLDKLKNKKKNL](http://crdd.osdd.net/raghava/ifnepitope/pep_design.php?sequence=NMDDLDKLKNKKKNL&method=hybrid&model=main) | MERCI | POSITIVE | 1 |
| 5 | 319-334 | [MDDLDKLKNKKKNLL](http://crdd.osdd.net/raghava/ifnepitope/pep_design.php?sequence=MDDLDKLKNKKKNLL&method=hybrid&model=main) | MERCI | POSITIVE | 1 |
| 6 | 320-335 | [DDLDKLKNKKKNLLY](http://crdd.osdd.net/raghava/ifnepitope/pep_design.php?sequence=DDLDKLKNKKKNLLY&method=hybrid&model=main) | MERCI | POSITIVE | 1 |
| 7 | 321-336 | [DLDKLKNKKKNLLYK](http://crdd.osdd.net/raghava/ifnepitope/pep_design.php?sequence=DLDKLKNKKKNLLYK&method=hybrid&model=main) | MERCI | POSITIVE | 1 |
| 8 | 322-337 | [LDKLKNKKKNLLYKI](http://crdd.osdd.net/raghava/ifnepitope/pep_design.php?sequence=LDKLKNKKKNLLYKI&method=hybrid&model=main) | MERCI | POSITIVE | 1 |
| 9 | 368-383 | [GSVITVQKKAKFVAA](http://crdd.osdd.net/raghava/ifnepitope/pep_design.php?sequence=GSVITVQKKAKFVAA&method=hybrid&model=main) | MERCI | POSITIVE | 1 |
| 10 | 369-384 | [SVITVQKKAKFVAAW](http://crdd.osdd.net/raghava/ifnepitope/pep_design.php?sequence=SVITVQKKAKFVAAW&method=hybrid&model=main) | MERCI | POSITIVE | 1 |
| 11 | 370-385 | [VITVQKKAKFVAAWT](http://crdd.osdd.net/raghava/ifnepitope/pep_design.php?sequence=VITVQKKAKFVAAWT&method=hybrid&model=main) | MERCI | POSITIVE | 1 |
| 12 | 371-386 | [ITVQKKAKFVAAWTL](http://crdd.osdd.net/raghava/ifnepitope/pep_design.php?sequence=ITVQKKAKFVAAWTL&method=hybrid&model=main) | MERCI | POSITIVE | 1 |
| 13 | 372-387 | [TVQKKAKFVAAWTLK](http://crdd.osdd.net/raghava/ifnepitope/pep_design.php?sequence=TVQKKAKFVAAWTLK&method=hybrid&model=main) | MERCI | POSITIVE | 1 |
| 14 | 373-388 | [VQKKAKFVAAWTLKA](http://crdd.osdd.net/raghava/ifnepitope/pep_design.php?sequence=VQKKAKFVAAWTLKA&method=hybrid&model=main) | MERCI | POSITIVE | 1 |
| 15 | 374-389 | [QKKAKFVAAWTLKAA](http://crdd.osdd.net/raghava/ifnepitope/pep_design.php?sequence=QKKAKFVAAWTLKAA&method=hybrid&model=main) | MERCI | POSITIVE | 1 |
| 16 | 0-15 | [EAAAKGIINTLQKYY](http://crdd.osdd.net/raghava/ifnepitope/pep_design.php?sequence=EAAAKGIINTLQKYY&method=hybrid&model=main) | MERCI | NEGATIVE | 2 |
| 17 | 1-16 | [AAAKGIINTLQKYYC](http://crdd.osdd.net/raghava/ifnepitope/pep_design.php?sequence=AAAKGIINTLQKYYC&method=hybrid&model=main) | MERCI | NEGATIVE | 2 |
| 18 | 2-17 | [AAKGIINTLQKYYCR](http://crdd.osdd.net/raghava/ifnepitope/pep_design.php?sequence=AAKGIINTLQKYYCR&method=hybrid&model=main) | MERCI | NEGATIVE | 1 |
| 19 | 13-28 | [YYCRVRGGRCAVLSC](http://crdd.osdd.net/raghava/ifnepitope/pep_design.php?sequence=YYCRVRGGRCAVLSC&method=hybrid&model=main) | MERCI | NEGATIVE | 1 |
| 20 | 14-29 | [YCRVRGGRCAVLSCL](http://crdd.osdd.net/raghava/ifnepitope/pep_design.php?sequence=YCRVRGGRCAVLSCL&method=hybrid&model=main) | MERCI | NEGATIVE | 2 |
| 21 | 15-30 | [CRVRGGRCAVLSCLP](http://crdd.osdd.net/raghava/ifnepitope/pep_design.php?sequence=CRVRGGRCAVLSCLP&method=hybrid&model=main) | MERCI | NEGATIVE | 2 |
| 22 | 16-31 | [RVRGGRCAVLSCLPK](http://crdd.osdd.net/raghava/ifnepitope/pep_design.php?sequence=RVRGGRCAVLSCLPK&method=hybrid&model=main) | MERCI | NEGATIVE | 2 |
| 23 | 17-32 | [VRGGRCAVLSCLPKE](http://crdd.osdd.net/raghava/ifnepitope/pep_design.php?sequence=VRGGRCAVLSCLPKE&method=hybrid&model=main) | MERCI | NEGATIVE | 2 |
| 24 | 18-33 | [RGGRCAVLSCLPKEE](http://crdd.osdd.net/raghava/ifnepitope/pep_design.php?sequence=RGGRCAVLSCLPKEE&method=hybrid&model=main) | MERCI | NEGATIVE | 2 |
| 25 | 19-34 | [GGRCAVLSCLPKEEQ](http://crdd.osdd.net/raghava/ifnepitope/pep_design.php?sequence=GGRCAVLSCLPKEEQ&method=hybrid&model=main) | MERCI | NEGATIVE | 2 |
| 26 | 20-35 | [GRCAVLSCLPKEEQI](http://crdd.osdd.net/raghava/ifnepitope/pep_design.php?sequence=GRCAVLSCLPKEEQI&method=hybrid&model=main) | MERCI | NEGATIVE | 2 |
| 27 | 59-74 | [AAWTLKAAAGGGSLD](http://crdd.osdd.net/raghava/ifnepitope/pep_design.php?sequence=AAWTLKAAAGGGSLD&method=hybrid&model=main) | MERCI | NEGATIVE | 1 |
| 28 | 60-75 | [AWTLKAAAGGGSLDF](http://crdd.osdd.net/raghava/ifnepitope/pep_design.php?sequence=AWTLKAAAGGGSLDF&method=hybrid&model=main) | MERCI | NEGATIVE | 1 |
| 29 | 61-76 | [WTLKAAAGGGSLDFS](http://crdd.osdd.net/raghava/ifnepitope/pep_design.php?sequence=WTLKAAAGGGSLDFS&method=hybrid&model=main) | MERCI | NEGATIVE | 1 |
| 30 | 62-77 | [TLKAAAGGGSLDFSE](http://crdd.osdd.net/raghava/ifnepitope/pep_design.php?sequence=TLKAAAGGGSLDFSE&method=hybrid&model=main) | MERCI | NEGATIVE | 1 |
| 31 | 63-78 | [LKAAAGGGSLDFSEV](http://crdd.osdd.net/raghava/ifnepitope/pep_design.php?sequence=LKAAAGGGSLDFSEV&method=hybrid&model=main) | MERCI | NEGATIVE | 1 |
| 32 | 64-79 | [KAAAGGGSLDFSEVS](http://crdd.osdd.net/raghava/ifnepitope/pep_design.php?sequence=KAAAGGGSLDFSEVS&method=hybrid&model=main) | MERCI | NEGATIVE | 1 |
| 33 | 68-83 | [GGGSLDFSEVSNVGG](http://crdd.osdd.net/raghava/ifnepitope/pep_design.php?sequence=GGGSLDFSEVSNVGG&method=hybrid&model=main) | MERCI | NEGATIVE | 1 |
| 34 | 69-84 | [GGSLDFSEVSNVGGG](http://crdd.osdd.net/raghava/ifnepitope/pep_design.php?sequence=GGSLDFSEVSNVGGG&method=hybrid&model=main) | MERCI | NEGATIVE | 1 |
| 35 | 70-85 | [GSLDFSEVSNVGGGS](http://crdd.osdd.net/raghava/ifnepitope/pep_design.php?sequence=GSLDFSEVSNVGGGS&method=hybrid&model=main) | MERCI | NEGATIVE | 1 |
| 36 | 71-86 | [SLDFSEVSNVGGGSI](http://crdd.osdd.net/raghava/ifnepitope/pep_design.php?sequence=SLDFSEVSNVGGGSI&method=hybrid&model=main) | MERCI | NEGATIVE | 1 |
| 37 | 72-87 | [LDFSEVSNVGGGSIE](http://crdd.osdd.net/raghava/ifnepitope/pep_design.php?sequence=LDFSEVSNVGGGSIE&method=hybrid&model=main) | MERCI | NEGATIVE | 1 |
| 38 | 73-88 | [DFSEVSNVGGGSIEG](http://crdd.osdd.net/raghava/ifnepitope/pep_design.php?sequence=DFSEVSNVGGGSIEG&method=hybrid&model=main) | MERCI | NEGATIVE | 1 |
| 39 | 74-89 | [FSEVSNVGGGSIEGR](http://crdd.osdd.net/raghava/ifnepitope/pep_design.php?sequence=FSEVSNVGGGSIEGR&method=hybrid&model=main) | MERCI | NEGATIVE | 1 |
| 40 | 75-90 | [SEVSNVGGGSIEGRP](http://crdd.osdd.net/raghava/ifnepitope/pep_design.php?sequence=SEVSNVGGGSIEGRP&method=hybrid&model=main) | MERCI | NEGATIVE | 1 |
| 41 | 76-91 | [EVSNVGGGSIEGRPE](http://crdd.osdd.net/raghava/ifnepitope/pep_design.php?sequence=EVSNVGGGSIEGRPE&method=hybrid&model=main) | MERCI | NEGATIVE | 1 |
| 42 | 85-100 | [IEGRPEDPVGGGSKD](http://crdd.osdd.net/raghava/ifnepitope/pep_design.php?sequence=IEGRPEDPVGGGSKD&method=hybrid&model=main) | MERCI | NEGATIVE | 1 |
| 43 | 86-101 | [EGRPEDPVGGGSKDS](http://crdd.osdd.net/raghava/ifnepitope/pep_design.php?sequence=EGRPEDPVGGGSKDS&method=hybrid&model=main) | MERCI | NEGATIVE | 1 |
| 44 | 87-102 | [GRPEDPVGGGSKDSS](http://crdd.osdd.net/raghava/ifnepitope/pep_design.php?sequence=GRPEDPVGGGSKDSS&method=hybrid&model=main) | MERCI | NEGATIVE | 1 |
| 45 | 88-103 | [RPEDPVGGGSKDSSL](http://crdd.osdd.net/raghava/ifnepitope/pep_design.php?sequence=RPEDPVGGGSKDSSL&method=hybrid&model=main) | MERCI | NEGATIVE | 1 |
| 46 | 89-104 | [PEDPVGGGSKDSSLL](http://crdd.osdd.net/raghava/ifnepitope/pep_design.php?sequence=PEDPVGGGSKDSSLL&method=hybrid&model=main) | MERCI | NEGATIVE | 1 |
| 47 | 90-105 | [EDPVGGGSKDSSLLN](http://crdd.osdd.net/raghava/ifnepitope/pep_design.php?sequence=EDPVGGGSKDSSLLN&method=hybrid&model=main) | MERCI | NEGATIVE | 1 |
| 48 | 91-106 | [DPVGGGSKDSSLLNN](http://crdd.osdd.net/raghava/ifnepitope/pep_design.php?sequence=DPVGGGSKDSSLLNN&method=hybrid&model=main) | MERCI | NEGATIVE | 1 |
| 49 | 100-115 | [SSLLNNQGGGSGWPY](http://crdd.osdd.net/raghava/ifnepitope/pep_design.php?sequence=SSLLNNQGGGSGWPY&method=hybrid&model=main) | MERCI | NEGATIVE | 1 |
| 50 | 101-116 | [SLLNNQGGGSGWPYI](http://crdd.osdd.net/raghava/ifnepitope/pep_design.php?sequence=SLLNNQGGGSGWPYI&method=hybrid&model=main) | MERCI | NEGATIVE | 1 |
| 51 | 102-117 | [LLNNQGGGSGWPYIG](http://crdd.osdd.net/raghava/ifnepitope/pep_design.php?sequence=LLNNQGGGSGWPYIG&method=hybrid&model=main) | MERCI | NEGATIVE | 1 |
| 52 | 103-118 | [LNNQGGGSGWPYIGS](http://crdd.osdd.net/raghava/ifnepitope/pep_design.php?sequence=LNNQGGGSGWPYIGS&method=hybrid&model=main) | MERCI | NEGATIVE | 1 |
| 53 | 104-119 | [NNQGGGSGWPYIGSR](http://crdd.osdd.net/raghava/ifnepitope/pep_design.php?sequence=NNQGGGSGWPYIGSR&method=hybrid&model=main) | MERCI | NEGATIVE | 1 |
| 54 | 121-136 | [PGPGALTDLGLLYTV](http://crdd.osdd.net/raghava/ifnepitope/pep_design.php?sequence=PGPGALTDLGLLYTV&method=hybrid&model=main) | MERCI | NEGATIVE | 1 |
| 55 | 122-137 | [GPGALTDLGLLYTVK](http://crdd.osdd.net/raghava/ifnepitope/pep_design.php?sequence=GPGALTDLGLLYTVK&method=hybrid&model=main) | MERCI | NEGATIVE | 3 |
| 56 | 123-138 | [PGALTDLGLLYTVKY](http://crdd.osdd.net/raghava/ifnepitope/pep_design.php?sequence=PGALTDLGLLYTVKY&method=hybrid&model=main) | MERCI | NEGATIVE | 3 |
| 57 | 124-139 | [GALTDLGLLYTVKYP](http://crdd.osdd.net/raghava/ifnepitope/pep_design.php?sequence=GALTDLGLLYTVKYP&method=hybrid&model=main) | MERCI | NEGATIVE | 3 |
| 58 | 125-140 | [ALTDLGLLYTVKYPN](http://crdd.osdd.net/raghava/ifnepitope/pep_design.php?sequence=ALTDLGLLYTVKYPN&method=hybrid&model=main) | MERCI | NEGATIVE | 3 |
| 59 | 126-141 | [LTDLGLLYTVKYPNG](http://crdd.osdd.net/raghava/ifnepitope/pep_design.php?sequence=LTDLGLLYTVKYPNG&method=hybrid&model=main) | MERCI | NEGATIVE | 3 |
| 60 | 127-142 | [TDLGLLYTVKYPNGP](http://crdd.osdd.net/raghava/ifnepitope/pep_design.php?sequence=TDLGLLYTVKYPNGP&method=hybrid&model=main) | MERCI | NEGATIVE | 3 |
| 61 | 128-143 | [DLGLLYTVKYPNGPG](http://crdd.osdd.net/raghava/ifnepitope/pep_design.php?sequence=DLGLLYTVKYPNGPG&method=hybrid&model=main) | MERCI | NEGATIVE | 2 |
| 62 | 180-195 | [GPGPGLTSLGLLYTV](http://crdd.osdd.net/raghava/ifnepitope/pep_design.php?sequence=GPGPGLTSLGLLYTV&method=hybrid&model=main) | MERCI | NEGATIVE | 1 |
| 63 | 181-196 | [PGPGLTSLGLLYTVK](http://crdd.osdd.net/raghava/ifnepitope/pep_design.php?sequence=PGPGLTSLGLLYTVK&method=hybrid&model=main) | MERCI | NEGATIVE | 3 |
| 64 | 182-197 | [GPGLTSLGLLYTVKY](http://crdd.osdd.net/raghava/ifnepitope/pep_design.php?sequence=GPGLTSLGLLYTVKY&method=hybrid&model=main) | MERCI | NEGATIVE | 3 |
| 65 | 183-198 | [PGLTSLGLLYTVKYP](http://crdd.osdd.net/raghava/ifnepitope/pep_design.php?sequence=PGLTSLGLLYTVKYP&method=hybrid&model=main) | MERCI | NEGATIVE | 3 |
| 66 | 184-199 | [GLTSLGLLYTVKYPN](http://crdd.osdd.net/raghava/ifnepitope/pep_design.php?sequence=GLTSLGLLYTVKYPN&method=hybrid&model=main) | MERCI | NEGATIVE | 3 |
| 67 | 185-200 | [LTSLGLLYTVKYPNL](http://crdd.osdd.net/raghava/ifnepitope/pep_design.php?sequence=LTSLGLLYTVKYPNL&method=hybrid&model=main) | MERCI | NEGATIVE | 3 |
| 68 | 186-201 | [TSLGLLYTVKYPNLK](http://crdd.osdd.net/raghava/ifnepitope/pep_design.php?sequence=TSLGLLYTVKYPNLK&method=hybrid&model=main) | MERCI | NEGATIVE | 3 |
| 69 | 187-202 | [SLGLLYTVKYPNLKK](http://crdd.osdd.net/raghava/ifnepitope/pep_design.php?sequence=SLGLLYTVKYPNLKK&method=hybrid&model=main) | MERCI | NEGATIVE | 2 |
| 70 | 289-304 | [ACLTKQKKKVVKDAV](http://crdd.osdd.net/raghava/ifnepitope/pep_design.php?sequence=ACLTKQKKKVVKDAV&method=hybrid&model=main) | MERCI | NEGATIVE | 1 |
| 71 | 290-305 | [CLTKQKKKVVKDAVS](http://crdd.osdd.net/raghava/ifnepitope/pep_design.php?sequence=CLTKQKKKVVKDAVS&method=hybrid&model=main) | MERCI | NEGATIVE | 1 |
| 72 | 291-306 | [LTKQKKKVVKDAVSL](http://crdd.osdd.net/raghava/ifnepitope/pep_design.php?sequence=LTKQKKKVVKDAVSL&method=hybrid&model=main) | MERCI | NEGATIVE | 1 |
| 73 | 292-307 | [TKQKKKVVKDAVSLI](http://crdd.osdd.net/raghava/ifnepitope/pep_design.php?sequence=TKQKKKVVKDAVSLI&method=hybrid&model=main) | MERCI | NEGATIVE | 1 |
| 74 | 293-308 | [KQKKKVVKDAVSLIN](http://crdd.osdd.net/raghava/ifnepitope/pep_design.php?sequence=KQKKKVVKDAVSLIN&method=hybrid&model=main) | MERCI | NEGATIVE | 1 |
| 75 | 294-309 | [QKKKVVKDAVSLING](http://crdd.osdd.net/raghava/ifnepitope/pep_design.php?sequence=QKKKVVKDAVSLING&method=hybrid&model=main) | MERCI | NEGATIVE | 1 |
| 76 | 295-310 | [KKKVVKDAVSLINGL](http://crdd.osdd.net/raghava/ifnepitope/pep_design.php?sequence=KKKVVKDAVSLINGL&method=hybrid&model=main) | MERCI | NEGATIVE | 1 |
| 77 | 296-311 | [KKVVKDAVSLINGLD](http://crdd.osdd.net/raghava/ifnepitope/pep_design.php?sequence=KKVVKDAVSLINGLD&method=hybrid&model=main) | MERCI | NEGATIVE | 1 |
| 78 | 323-338 | [DKLKNKKKNLLYKIC](http://crdd.osdd.net/raghava/ifnepitope/pep_design.php?sequence=DKLKNKKKNLLYKIC&method=hybrid&model=main) | MERCI | NEGATIVE | 1 |
| 79 | 324-339 | [KLKNKKKNLLYKICL](http://crdd.osdd.net/raghava/ifnepitope/pep_design.php?sequence=KLKNKKKNLLYKICL&method=hybrid&model=main) | MERCI | NEGATIVE | 3 |
| 80 | 325-340 | [LKNKKKNLLYKICLS](http://crdd.osdd.net/raghava/ifnepitope/pep_design.php?sequence=LKNKKKNLLYKICLS&method=hybrid&model=main) | MERCI | NEGATIVE | 3 |
| 81 | 326-341 | [KNKKKNLLYKICLSG](http://crdd.osdd.net/raghava/ifnepitope/pep_design.php?sequence=KNKKKNLLYKICLSG&method=hybrid&model=main) | MERCI | NEGATIVE | 3 |
| 82 | 327-342 | [NKKKNLLYKICLSGK](http://crdd.osdd.net/raghava/ifnepitope/pep_design.php?sequence=NKKKNLLYKICLSGK&method=hybrid&model=main) | MERCI | NEGATIVE | 3 |
| 83 | 328-343 | [KKKNLLYKICLSGKK](http://crdd.osdd.net/raghava/ifnepitope/pep_design.php?sequence=KKKNLLYKICLSGKK&method=hybrid&model=main) | MERCI | NEGATIVE | 3 |
| 84 | 329-344 | [KKNLLYKICLSGKKD](http://crdd.osdd.net/raghava/ifnepitope/pep_design.php?sequence=KKNLLYKICLSGKKD&method=hybrid&model=main) | MERCI | NEGATIVE | 2 |
| 85 | 330-345 | [KNLLYKICLSGKKDE](http://crdd.osdd.net/raghava/ifnepitope/pep_design.php?sequence=KNLLYKICLSGKKDE&method=hybrid&model=main) | MERCI | NEGATIVE | 2 |
| 86 | 331-346 | [NLLYKICLSGKKDER](http://crdd.osdd.net/raghava/ifnepitope/pep_design.php?sequence=NLLYKICLSGKKDER&method=hybrid&model=main) | MERCI | NEGATIVE | 2 |
| 87 | 332-347 | [LLYKICLSGKKDERP](http://crdd.osdd.net/raghava/ifnepitope/pep_design.php?sequence=LLYKICLSGKKDERP&method=hybrid&model=main) | MERCI | NEGATIVE | 2 |
| 88 | 333-348 | [LYKICLSGKKDERPG](http://crdd.osdd.net/raghava/ifnepitope/pep_design.php?sequence=LYKICLSGKKDERPG&method=hybrid&model=main) | MERCI | NEGATIVE | 1 |
| 89 | 348-363 | [NRNPYKKPGLLSYVI](http://crdd.osdd.net/raghava/ifnepitope/pep_design.php?sequence=NRNPYKKPGLLSYVI&method=hybrid&model=main) | MERCI | NEGATIVE | 2 |
| 90 | 349-364 | [RNPYKKPGLLSYVIG](http://crdd.osdd.net/raghava/ifnepitope/pep_design.php?sequence=RNPYKKPGLLSYVIG&method=hybrid&model=main) | MERCI | NEGATIVE | 2 |
| 91 | 350-365 | [NPYKKPGLLSYVIGL](http://crdd.osdd.net/raghava/ifnepitope/pep_design.php?sequence=NPYKKPGLLSYVIGL&method=hybrid&model=main) | MERCI | NEGATIVE | 2 |
| 92 | 351-366 | [PYKKPGLLSYVIGLL](http://crdd.osdd.net/raghava/ifnepitope/pep_design.php?sequence=PYKKPGLLSYVIGLL&method=hybrid&model=main) | MERCI | NEGATIVE | 8 |
| 93 | 352-367 | [YKKPGLLSYVIGLLP](http://crdd.osdd.net/raghava/ifnepitope/pep_design.php?sequence=YKKPGLLSYVIGLLP&method=hybrid&model=main) | MERCI | NEGATIVE | 10 |
| 94 | 353-368 | [KKPGLLSYVIGLLPQ](http://crdd.osdd.net/raghava/ifnepitope/pep_design.php?sequence=KKPGLLSYVIGLLPQ&method=hybrid&model=main) | MERCI | NEGATIVE | 10 |
| 95 | 354-369 | [KPGLLSYVIGLLPQG](http://crdd.osdd.net/raghava/ifnepitope/pep_design.php?sequence=KPGLLSYVIGLLPQG&method=hybrid&model=main) | MERCI | NEGATIVE | 10 |
| 96 | 355-370 | [PGLLSYVIGLLPQGS](http://crdd.osdd.net/raghava/ifnepitope/pep_design.php?sequence=PGLLSYVIGLLPQGS&method=hybrid&model=main) | MERCI | NEGATIVE | 10 |
| 97 | 356-371 | [GLLSYVIGLLPQGSV](http://crdd.osdd.net/raghava/ifnepitope/pep_design.php?sequence=GLLSYVIGLLPQGSV&method=hybrid&model=main) | MERCI | NEGATIVE | 10 |
| 98 | 357-372 | [LLSYVIGLLPQGSVI](http://crdd.osdd.net/raghava/ifnepitope/pep_design.php?sequence=LLSYVIGLLPQGSVI&method=hybrid&model=main) | MERCI | NEGATIVE | 11 |
| 99 | 358-373 | [LSYVIGLLPQGSVIT](http://crdd.osdd.net/raghava/ifnepitope/pep_design.php?sequence=LSYVIGLLPQGSVIT&method=hybrid&model=main) | MERCI | NEGATIVE | 9 |
| 100 | 359-374 | [SYVIGLLPQGSVITV](http://crdd.osdd.net/raghava/ifnepitope/pep_design.php?sequence=SYVIGLLPQGSVITV&method=hybrid&model=main) | MERCI | NEGATIVE | 9 |
| 101 | 360-375 | [YVIGLLPQGSVITVQ](http://crdd.osdd.net/raghava/ifnepitope/pep_design.php?sequence=YVIGLLPQGSVITVQ&method=hybrid&model=main) | MERCI | NEGATIVE | 7 |
| 102 | 361-376 | [VIGLLPQGSVITVQK](http://crdd.osdd.net/raghava/ifnepitope/pep_design.php?sequence=VIGLLPQGSVITVQK&method=hybrid&model=main) | MERCI | NEGATIVE | 4 |
| 103 | 362-377 | [IGLLPQGSVITVQKK](http://crdd.osdd.net/raghava/ifnepitope/pep_design.php?sequence=IGLLPQGSVITVQKK&method=hybrid&model=main) | MERCI | NEGATIVE | 2 |
| 104 | 363-378 | [GLLPQGSVITVQKKA](http://crdd.osdd.net/raghava/ifnepitope/pep_design.php?sequence=GLLPQGSVITVQKKA&method=hybrid&model=main) | MERCI | NEGATIVE | 1 |
| 105 | 364-379 | [LLPQGSVITVQKKAK](http://crdd.osdd.net/raghava/ifnepitope/pep_design.php?sequence=LLPQGSVITVQKKAK&method=hybrid&model=main) | MERCI | NEGATIVE | 1 |
| 106 | 3-18 | [AKGIINTLQKYYCRV](http://crdd.osdd.net/raghava/ifnepitope/pep_design.php?sequence=AKGIINTLQKYYCRV&method=hybrid&model=main) | SVM | NEGATIVE | -0.4811069 |
| 107 | 4-19 | [KGIINTLQKYYCRVR](http://crdd.osdd.net/raghava/ifnepitope/pep_design.php?sequence=KGIINTLQKYYCRVR&method=hybrid&model=main) | SVM | NEGATIVE | -0.40192947 |
| 108 | 5-20 | [GIINTLQKYYCRVRG](http://crdd.osdd.net/raghava/ifnepitope/pep_design.php?sequence=GIINTLQKYYCRVRG&method=hybrid&model=main) | SVM | NEGATIVE | -0.37130542 |
| 109 | 6-21 | [IINTLQKYYCRVRGG](http://crdd.osdd.net/raghava/ifnepitope/pep_design.php?sequence=IINTLQKYYCRVRGG&method=hybrid&model=main) | SVM | NEGATIVE | -0.58743555 |
| 110 | 7-22 | [INTLQKYYCRVRGGR](http://crdd.osdd.net/raghava/ifnepitope/pep_design.php?sequence=INTLQKYYCRVRGGR&method=hybrid&model=main) | SVM | NEGATIVE | -0.65489384 |
| 111 | 8-23 | [NTLQKYYCRVRGGRC](http://crdd.osdd.net/raghava/ifnepitope/pep_design.php?sequence=NTLQKYYCRVRGGRC&method=hybrid&model=main) | SVM | NEGATIVE | -0.57124458 |
| 112 | 9-24 | [TLQKYYCRVRGGRCA](http://crdd.osdd.net/raghava/ifnepitope/pep_design.php?sequence=TLQKYYCRVRGGRCA&method=hybrid&model=main) | SVM | NEGATIVE | -0.38970485 |
| 113 | 10-25 | [LQKYYCRVRGGRCAV](http://crdd.osdd.net/raghava/ifnepitope/pep_design.php?sequence=LQKYYCRVRGGRCAV&method=hybrid&model=main) | SVM | NEGATIVE | -0.50913676 |
| 114 | 11-26 | [QKYYCRVRGGRCAVL](http://crdd.osdd.net/raghava/ifnepitope/pep_design.php?sequence=QKYYCRVRGGRCAVL&method=hybrid&model=main) | SVM | NEGATIVE | -0.38085703 |
| 115 | 12-27 | [KYYCRVRGGRCAVLS](http://crdd.osdd.net/raghava/ifnepitope/pep_design.php?sequence=KYYCRVRGGRCAVLS&method=hybrid&model=main) | SVM | NEGATIVE | -0.2994713 |
| 116 | 21-36 | [RCAVLSCLPKEEQIG](http://crdd.osdd.net/raghava/ifnepitope/pep_design.php?sequence=RCAVLSCLPKEEQIG&method=hybrid&model=main) | SVM | NEGATIVE | -0.86346741 |
| 117 | 22-37 | [CAVLSCLPKEEQIGK](http://crdd.osdd.net/raghava/ifnepitope/pep_design.php?sequence=CAVLSCLPKEEQIGK&method=hybrid&model=main) | SVM | NEGATIVE | -0.74464143 |
| 118 | 23-38 | [AVLSCLPKEEQIGKC](http://crdd.osdd.net/raghava/ifnepitope/pep_design.php?sequence=AVLSCLPKEEQIGKC&method=hybrid&model=main) | SVM | NEGATIVE | -0.6976502 |
| 119 | 24-39 | [VLSCLPKEEQIGKCS](http://crdd.osdd.net/raghava/ifnepitope/pep_design.php?sequence=VLSCLPKEEQIGKCS&method=hybrid&model=main) | SVM | NEGATIVE | -0.81952843 |
| 120 | 25-40 | [LSCLPKEEQIGKCST](http://crdd.osdd.net/raghava/ifnepitope/pep_design.php?sequence=LSCLPKEEQIGKCST&method=hybrid&model=main) | SVM | NEGATIVE | -0.74976887 |
| 121 | 26-41 | [SCLPKEEQIGKCSTR](http://crdd.osdd.net/raghava/ifnepitope/pep_design.php?sequence=SCLPKEEQIGKCSTR&method=hybrid&model=main) | SVM | NEGATIVE | -0.53112116 |
| 122 | 27-42 | [CLPKEEQIGKCSTRG](http://crdd.osdd.net/raghava/ifnepitope/pep_design.php?sequence=CLPKEEQIGKCSTRG&method=hybrid&model=main) | SVM | NEGATIVE | -0.34688682 |
| 123 | 28-43 | [LPKEEQIGKCSTRGR](http://crdd.osdd.net/raghava/ifnepitope/pep_design.php?sequence=LPKEEQIGKCSTRGR&method=hybrid&model=main) | SVM | NEGATIVE | -0.33297862 |
| 124 | 29-44 | [PKEEQIGKCSTRGRK](http://crdd.osdd.net/raghava/ifnepitope/pep_design.php?sequence=PKEEQIGKCSTRGRK&method=hybrid&model=main) | SVM | POSITIVE | 0.24193665 |
| 125 | 30-45 | [KEEQIGKCSTRGRKC](http://crdd.osdd.net/raghava/ifnepitope/pep_design.php?sequence=KEEQIGKCSTRGRKC&method=hybrid&model=main) | SVM | POSITIVE | 0.73719216 |
| 126 | 31-46 | [EEQIGKCSTRGRKCC](http://crdd.osdd.net/raghava/ifnepitope/pep_design.php?sequence=EEQIGKCSTRGRKCC&method=hybrid&model=main) | SVM | POSITIVE | 0.82166713 |
| 127 | 32-47 | [EQIGKCSTRGRKCCR](http://crdd.osdd.net/raghava/ifnepitope/pep_design.php?sequence=EQIGKCSTRGRKCCR&method=hybrid&model=main) | SVM | POSITIVE | 0.55986934 |
| 128 | 33-48 | [QIGKCSTRGRKCCRR](http://crdd.osdd.net/raghava/ifnepitope/pep_design.php?sequence=QIGKCSTRGRKCCRR&method=hybrid&model=main) | SVM | POSITIVE | 0.37535174 |
| 129 | 34-49 | [IGKCSTRGRKCCRRK](http://crdd.osdd.net/raghava/ifnepitope/pep_design.php?sequence=IGKCSTRGRKCCRRK&method=hybrid&model=main) | SVM | POSITIVE | 0.69267842 |
| 130 | 35-50 | [GKCSTRGRKCCRRKK](http://crdd.osdd.net/raghava/ifnepitope/pep_design.php?sequence=GKCSTRGRKCCRRKK&method=hybrid&model=main) | SVM | POSITIVE | 0.63686641 |
| 131 | 36-51 | [KCSTRGRKCCRRKKE](http://crdd.osdd.net/raghava/ifnepitope/pep_design.php?sequence=KCSTRGRKCCRRKKE&method=hybrid&model=main) | SVM | POSITIVE | 0.56732319 |
| 132 | 37-52 | [CSTRGRKCCRRKKEA](http://crdd.osdd.net/raghava/ifnepitope/pep_design.php?sequence=CSTRGRKCCRRKKEA&method=hybrid&model=main) | SVM | POSITIVE | 0.29576505 |
| 133 | 38-53 | [STRGRKCCRRKKEAA](http://crdd.osdd.net/raghava/ifnepitope/pep_design.php?sequence=STRGRKCCRRKKEAA&method=hybrid&model=main) | SVM | POSITIVE | 0.37433555 |
| 134 | 39-54 | [TRGRKCCRRKKEAAA](http://crdd.osdd.net/raghava/ifnepitope/pep_design.php?sequence=TRGRKCCRRKKEAAA&method=hybrid&model=main) | SVM | POSITIVE | 0.49026956 |
| 135 | 40-55 | [RGRKCCRRKKEAAAK](http://crdd.osdd.net/raghava/ifnepitope/pep_design.php?sequence=RGRKCCRRKKEAAAK&method=hybrid&model=main) | SVM | POSITIVE | 0.34262454 |
| 136 | 41-56 | [GRKCCRRKKEAAAKA](http://crdd.osdd.net/raghava/ifnepitope/pep_design.php?sequence=GRKCCRRKKEAAAKA&method=hybrid&model=main) | SVM | POSITIVE | 0.2628675 |
| 137 | 42-57 | [RKCCRRKKEAAAKAK](http://crdd.osdd.net/raghava/ifnepitope/pep_design.php?sequence=RKCCRRKKEAAAKAK&method=hybrid&model=main) | SVM | POSITIVE | 0.23815639 |
| 138 | 43-58 | [KCCRRKKEAAAKAKF](http://crdd.osdd.net/raghava/ifnepitope/pep_design.php?sequence=KCCRRKKEAAAKAKF&method=hybrid&model=main) | SVM | POSITIVE | 0.26126896 |
| 139 | 44-59 | [CCRRKKEAAAKAKFV](http://crdd.osdd.net/raghava/ifnepitope/pep_design.php?sequence=CCRRKKEAAAKAKFV&method=hybrid&model=main) | SVM | POSITIVE | 0.010566599 |
| 140 | 45-60 | [CRRKKEAAAKAKFVA](http://crdd.osdd.net/raghava/ifnepitope/pep_design.php?sequence=CRRKKEAAAKAKFVA&method=hybrid&model=main) | SVM | POSITIVE | 0.21963392 |
| 141 | 46-61 | [RRKKEAAAKAKFVAA](http://crdd.osdd.net/raghava/ifnepitope/pep_design.php?sequence=RRKKEAAAKAKFVAA&method=hybrid&model=main) | SVM | POSITIVE | 0.73119114 |
| 142 | 47-62 | [RKKEAAAKAKFVAAW](http://crdd.osdd.net/raghava/ifnepitope/pep_design.php?sequence=RKKEAAAKAKFVAAW&method=hybrid&model=main) | SVM | POSITIVE | 0.89663429 |
| 143 | 48-63 | [KKEAAAKAKFVAAWT](http://crdd.osdd.net/raghava/ifnepitope/pep_design.php?sequence=KKEAAAKAKFVAAWT&method=hybrid&model=main) | SVM | POSITIVE | 1.1882513 |
| 144 | 49-64 | [KEAAAKAKFVAAWTL](http://crdd.osdd.net/raghava/ifnepitope/pep_design.php?sequence=KEAAAKAKFVAAWTL&method=hybrid&model=main) | SVM | POSITIVE | 1.1490985 |
| 145 | 50-65 | [EAAAKAKFVAAWTLK](http://crdd.osdd.net/raghava/ifnepitope/pep_design.php?sequence=EAAAKAKFVAAWTLK&method=hybrid&model=main) | SVM | POSITIVE | 1.1240097 |
| 146 | 51-66 | [AAAKAKFVAAWTLKA](http://crdd.osdd.net/raghava/ifnepitope/pep_design.php?sequence=AAAKAKFVAAWTLKA&method=hybrid&model=main) | SVM | POSITIVE | 1.111491 |
| 147 | 52-67 | [AAKAKFVAAWTLKAA](http://crdd.osdd.net/raghava/ifnepitope/pep_design.php?sequence=AAKAKFVAAWTLKAA&method=hybrid&model=main) | SVM | POSITIVE | 1.111491 |
| 148 | 53-68 | [AKAKFVAAWTLKAAA](http://crdd.osdd.net/raghava/ifnepitope/pep_design.php?sequence=AKAKFVAAWTLKAAA&method=hybrid&model=main) | SVM | POSITIVE | 1.111491 |
| 149 | 54-69 | [KAKFVAAWTLKAAAG](http://crdd.osdd.net/raghava/ifnepitope/pep_design.php?sequence=KAKFVAAWTLKAAAG&method=hybrid&model=main) | SVM | POSITIVE | 0.79460947 |
| 150 | 55-70 | [AKFVAAWTLKAAAGG](http://crdd.osdd.net/raghava/ifnepitope/pep_design.php?sequence=AKFVAAWTLKAAAGG&method=hybrid&model=main) | SVM | POSITIVE | 0.61623789 |
| 151 | 56-71 | [KFVAAWTLKAAAGGG](http://crdd.osdd.net/raghava/ifnepitope/pep_design.php?sequence=KFVAAWTLKAAAGGG&method=hybrid&model=main) | SVM | POSITIVE | 0.67221404 |
| 152 | 57-72 | [FVAAWTLKAAAGGGS](http://crdd.osdd.net/raghava/ifnepitope/pep_design.php?sequence=FVAAWTLKAAAGGGS&method=hybrid&model=main) | SVM | POSITIVE | 0.52759093 |
| 153 | 58-73 | [VAAWTLKAAAGGGSL](http://crdd.osdd.net/raghava/ifnepitope/pep_design.php?sequence=VAAWTLKAAAGGGSL&method=hybrid&model=main) | SVM | POSITIVE | 0.54670018 |
| 154 | 65-80 | [AAAGGGSLDFSEVSN](http://crdd.osdd.net/raghava/ifnepitope/pep_design.php?sequence=AAAGGGSLDFSEVSN&method=hybrid&model=main) | SVM | POSITIVE | 0.0035323699 |
| 155 | 66-81 | [AAGGGSLDFSEVSNV](http://crdd.osdd.net/raghava/ifnepitope/pep_design.php?sequence=AAGGGSLDFSEVSNV&method=hybrid&model=main) | SVM | NEGATIVE | -0.073686789 |
| 156 | 67-82 | [AGGGSLDFSEVSNVG](http://crdd.osdd.net/raghava/ifnepitope/pep_design.php?sequence=AGGGSLDFSEVSNVG&method=hybrid&model=main) | SVM | POSITIVE | 0.0014408598 |
| 157 | 77-92 | [VSNVGGGSIEGRPED](http://crdd.osdd.net/raghava/ifnepitope/pep_design.php?sequence=VSNVGGGSIEGRPED&method=hybrid&model=main) | SVM | POSITIVE | 0.40858957 |
| 158 | 78-93 | [SNVGGGSIEGRPEDP](http://crdd.osdd.net/raghava/ifnepitope/pep_design.php?sequence=SNVGGGSIEGRPEDP&method=hybrid&model=main) | SVM | POSITIVE | 0.56861828 |
| 159 | 79-94 | [NVGGGSIEGRPEDPV](http://crdd.osdd.net/raghava/ifnepitope/pep_design.php?sequence=NVGGGSIEGRPEDPV&method=hybrid&model=main) | SVM | POSITIVE | 0.5680472 |
| 160 | 80-95 | [VGGGSIEGRPEDPVG](http://crdd.osdd.net/raghava/ifnepitope/pep_design.php?sequence=VGGGSIEGRPEDPVG&method=hybrid&model=main) | SVM | POSITIVE | 0.57546466 |
| 161 | 81-96 | [GGGSIEGRPEDPVGG](http://crdd.osdd.net/raghava/ifnepitope/pep_design.php?sequence=GGGSIEGRPEDPVGG&method=hybrid&model=main) | SVM | POSITIVE | 0.91993946 |
| 162 | 82-97 | [GGSIEGRPEDPVGGG](http://crdd.osdd.net/raghava/ifnepitope/pep_design.php?sequence=GGSIEGRPEDPVGGG&method=hybrid&model=main) | SVM | POSITIVE | 0.91993946 |
| 163 | 83-98 | [GSIEGRPEDPVGGGS](http://crdd.osdd.net/raghava/ifnepitope/pep_design.php?sequence=GSIEGRPEDPVGGGS&method=hybrid&model=main) | SVM | POSITIVE | 0.65581632 |
| 164 | 84-99 | [SIEGRPEDPVGGGSK](http://crdd.osdd.net/raghava/ifnepitope/pep_design.php?sequence=SIEGRPEDPVGGGSK&method=hybrid&model=main) | SVM | POSITIVE | 0.53971395 |
| 165 | 92-107 | [PVGGGSKDSSLLNNQ](http://crdd.osdd.net/raghava/ifnepitope/pep_design.php?sequence=PVGGGSKDSSLLNNQ&method=hybrid&model=main) | SVM | NEGATIVE | -0.18643217 |
| 166 | 93-108 | [VGGGSKDSSLLNNQG](http://crdd.osdd.net/raghava/ifnepitope/pep_design.php?sequence=VGGGSKDSSLLNNQG&method=hybrid&model=main) | SVM | NEGATIVE | -0.15047304 |
| 167 | 94-109 | [GGGSKDSSLLNNQGG](http://crdd.osdd.net/raghava/ifnepitope/pep_design.php?sequence=GGGSKDSSLLNNQGG&method=hybrid&model=main) | SVM | POSITIVE | 0.31503207 |
| 168 | 95-110 | [GGSKDSSLLNNQGGG](http://crdd.osdd.net/raghava/ifnepitope/pep_design.php?sequence=GGSKDSSLLNNQGGG&method=hybrid&model=main) | SVM | POSITIVE | 0.31503207 |
| 169 | 96-111 | [GSKDSSLLNNQGGGS](http://crdd.osdd.net/raghava/ifnepitope/pep_design.php?sequence=GSKDSSLLNNQGGGS&method=hybrid&model=main) | SVM | POSITIVE | 0.17631271 |
| 170 | 97-112 | [SKDSSLLNNQGGGSG](http://crdd.osdd.net/raghava/ifnepitope/pep_design.php?sequence=SKDSSLLNNQGGGSG&method=hybrid&model=main) | SVM | NEGATIVE | -0.14324969 |
| 171 | 98-113 | [KDSSLLNNQGGGSGW](http://crdd.osdd.net/raghava/ifnepitope/pep_design.php?sequence=KDSSLLNNQGGGSGW&method=hybrid&model=main) | SVM | POSITIVE | 0.20692182 |
| 172 | 99-114 | [DSSLLNNQGGGSGWP](http://crdd.osdd.net/raghava/ifnepitope/pep_design.php?sequence=DSSLLNNQGGGSGWP&method=hybrid&model=main) | SVM | POSITIVE | 0.36486518 |
| 173 | 105-120 | [NQGGGSGWPYIGSRS](http://crdd.osdd.net/raghava/ifnepitope/pep_design.php?sequence=NQGGGSGWPYIGSRS&method=hybrid&model=main) | SVM | POSITIVE | 0.13535092 |
| 174 | 107-122 | [GGGSGWPYIGSRSGP](http://crdd.osdd.net/raghava/ifnepitope/pep_design.php?sequence=GGGSGWPYIGSRSGP&method=hybrid&model=main) | SVM | POSITIVE | 0.048681536 |
| 175 | 108-123 | [GGSGWPYIGSRSGPG](http://crdd.osdd.net/raghava/ifnepitope/pep_design.php?sequence=GGSGWPYIGSRSGPG&method=hybrid&model=main) | SVM | NEGATIVE | -0.17141292 |
| 176 | 109-124 | [GSGWPYIGSRSGPGP](http://crdd.osdd.net/raghava/ifnepitope/pep_design.php?sequence=GSGWPYIGSRSGPGP&method=hybrid&model=main) | SVM | NEGATIVE | -0.6017 |
| 177 | 110-125 | [SGWPYIGSRSGPGPG](http://crdd.osdd.net/raghava/ifnepitope/pep_design.php?sequence=SGWPYIGSRSGPGPG&method=hybrid&model=main) | SVM | NEGATIVE | -0.37884433 |
| 178 | 111-126 | [GWPYIGSRSGPGPGA](http://crdd.osdd.net/raghava/ifnepitope/pep_design.php?sequence=GWPYIGSRSGPGPGA&method=hybrid&model=main) | SVM | NEGATIVE | -0.26505028 |
| 179 | 112-127 | [WPYIGSRSGPGPGAL](http://crdd.osdd.net/raghava/ifnepitope/pep_design.php?sequence=WPYIGSRSGPGPGAL&method=hybrid&model=main) | SVM | NEGATIVE | -0.49608855 |
| 180 | 113-128 | [PYIGSRSGPGPGALT](http://crdd.osdd.net/raghava/ifnepitope/pep_design.php?sequence=PYIGSRSGPGPGALT&method=hybrid&model=main) | SVM | NEGATIVE | -0.42239921 |
| 181 | 114-129 | [YIGSRSGPGPGALTD](http://crdd.osdd.net/raghava/ifnepitope/pep_design.php?sequence=YIGSRSGPGPGALTD&method=hybrid&model=main) | SVM | NEGATIVE | -0.14768995 |
| 182 | 115-130 | [IGSRSGPGPGALTDL](http://crdd.osdd.net/raghava/ifnepitope/pep_design.php?sequence=IGSRSGPGPGALTDL&method=hybrid&model=main) | SVM | NEGATIVE | -0.21274593 |
| 183 | 116-131 | [GSRSGPGPGALTDLG](http://crdd.osdd.net/raghava/ifnepitope/pep_design.php?sequence=GSRSGPGPGALTDLG&method=hybrid&model=main) | SVM | NEGATIVE | -0.34932932 |
| 184 | 117-132 | [SRSGPGPGALTDLGL](http://crdd.osdd.net/raghava/ifnepitope/pep_design.php?sequence=SRSGPGPGALTDLGL&method=hybrid&model=main) | SVM | POSITIVE | 0.061270902 |
| 185 | 118-133 | [RSGPGPGALTDLGLL](http://crdd.osdd.net/raghava/ifnepitope/pep_design.php?sequence=RSGPGPGALTDLGLL&method=hybrid&model=main) | SVM | POSITIVE | 0.0018756363 |
| 186 | 119-134 | [SGPGPGALTDLGLLY](http://crdd.osdd.net/raghava/ifnepitope/pep_design.php?sequence=SGPGPGALTDLGLLY&method=hybrid&model=main) | SVM | POSITIVE | 0.02455866 |
| 187 | 120-135 | [GPGPGALTDLGLLYT](http://crdd.osdd.net/raghava/ifnepitope/pep_design.php?sequence=GPGPGALTDLGLLYT&method=hybrid&model=main) | SVM | POSITIVE | 0.20604552 |
| 188 | 129-144 | [LGLLYTVKYPNGPGP](http://crdd.osdd.net/raghava/ifnepitope/pep_design.php?sequence=LGLLYTVKYPNGPGP&method=hybrid&model=main) | SVM | POSITIVE | 0.15974818 |
| 189 | 130-145 | [GLLYTVKYPNGPGPG](http://crdd.osdd.net/raghava/ifnepitope/pep_design.php?sequence=GLLYTVKYPNGPGPG&method=hybrid&model=main) | SVM | POSITIVE | 0.118329 |
| 190 | 131-146 | [LLYTVKYPNGPGPGN](http://crdd.osdd.net/raghava/ifnepitope/pep_design.php?sequence=LLYTVKYPNGPGPGN&method=hybrid&model=main) | SVM | NEGATIVE | -0.14813211 |
| 191 | 132-147 | [LYTVKYPNGPGPGNN](http://crdd.osdd.net/raghava/ifnepitope/pep_design.php?sequence=LYTVKYPNGPGPGNN&method=hybrid&model=main) | SVM | NEGATIVE | -0.14681072 |
| 192 | 133-148 | [YTVKYPNGPGPGNNQ](http://crdd.osdd.net/raghava/ifnepitope/pep_design.php?sequence=YTVKYPNGPGPGNNQ&method=hybrid&model=main) | SVM | NEGATIVE | -0.35594363 |
| 193 | 134-149 | [TVKYPNGPGPGNNQF](http://crdd.osdd.net/raghava/ifnepitope/pep_design.php?sequence=TVKYPNGPGPGNNQF&method=hybrid&model=main) | SVM | NEGATIVE | -0.36651667 |
| 194 | 135-150 | [VKYPNGPGPGNNQFG](http://crdd.osdd.net/raghava/ifnepitope/pep_design.php?sequence=VKYPNGPGPGNNQFG&method=hybrid&model=main) | SVM | NEGATIVE | -0.46967402 |
| 195 | 136-151 | [KYPNGPGPGNNQFGT](http://crdd.osdd.net/raghava/ifnepitope/pep_design.php?sequence=KYPNGPGPGNNQFGT&method=hybrid&model=main) | SVM | NEGATIVE | -0.63726271 |
| 196 | 137-152 | [YPNGPGPGNNQFGTM](http://crdd.osdd.net/raghava/ifnepitope/pep_design.php?sequence=YPNGPGPGNNQFGTM&method=hybrid&model=main) | SVM | NEGATIVE | -0.7910627 |
| 197 | 138-153 | [PNGPGPGNNQFGTMP](http://crdd.osdd.net/raghava/ifnepitope/pep_design.php?sequence=PNGPGPGNNQFGTMP&method=hybrid&model=main) | SVM | NEGATIVE | -0.70644064 |
| 198 | 139-154 | [NGPGPGNNQFGTMPS](http://crdd.osdd.net/raghava/ifnepitope/pep_design.php?sequence=NGPGPGNNQFGTMPS&method=hybrid&model=main) | SVM | NEGATIVE | -0.56721031 |
| 199 | 140-155 | [GPGPGNNQFGTMPSL](http://crdd.osdd.net/raghava/ifnepitope/pep_design.php?sequence=GPGPGNNQFGTMPSL&method=hybrid&model=main) | SVM | NEGATIVE | -0.63370584 |
| 200 | 141-156 | [PGPGNNQFGTMPSLT](http://crdd.osdd.net/raghava/ifnepitope/pep_design.php?sequence=PGPGNNQFGTMPSLT&method=hybrid&model=main) | SVM | NEGATIVE | -0.50517736 |
| 201 | 142-157 | [GPGNNQFGTMPSLTL](http://crdd.osdd.net/raghava/ifnepitope/pep_design.php?sequence=GPGNNQFGTMPSLTL&method=hybrid&model=main) | SVM | NEGATIVE | -0.8805301 |
| 202 | 143-158 | [PGNNQFGTMPSLTLA](http://crdd.osdd.net/raghava/ifnepitope/pep_design.php?sequence=PGNNQFGTMPSLTLA&method=hybrid&model=main) | SVM | NEGATIVE | -0.76835754 |
| 203 | 144-159 | [GNNQFGTMPSLTLAC](http://crdd.osdd.net/raghava/ifnepitope/pep_design.php?sequence=GNNQFGTMPSLTLAC&method=hybrid&model=main) | SVM | NEGATIVE | -1.0125524 |
| 204 | 145-160 | [NNQFGTMPSLTLACL](http://crdd.osdd.net/raghava/ifnepitope/pep_design.php?sequence=NNQFGTMPSLTLACL&method=hybrid&model=main) | SVM | NEGATIVE | -1.1920569 |
| 205 | 146-161 | [NQFGTMPSLTLACLG](http://crdd.osdd.net/raghava/ifnepitope/pep_design.php?sequence=NQFGTMPSLTLACLG&method=hybrid&model=main) | SVM | NEGATIVE | -1.100651 |
| 206 | 147-162 | [QFGTMPSLTLACLGP](http://crdd.osdd.net/raghava/ifnepitope/pep_design.php?sequence=QFGTMPSLTLACLGP&method=hybrid&model=main) | SVM | NEGATIVE | -1.1699767 |
| 207 | 148-163 | [FGTMPSLTLACLGPG](http://crdd.osdd.net/raghava/ifnepitope/pep_design.php?sequence=FGTMPSLTLACLGPG&method=hybrid&model=main) | SVM | NEGATIVE | -1.0188869 |
| 208 | 149-164 | [GTMPSLTLACLGPGP](http://crdd.osdd.net/raghava/ifnepitope/pep_design.php?sequence=GTMPSLTLACLGPGP&method=hybrid&model=main) | SVM | NEGATIVE | -0.82375517 |
| 209 | 150-165 | [TMPSLTLACLGPGPG](http://crdd.osdd.net/raghava/ifnepitope/pep_design.php?sequence=TMPSLTLACLGPGPG&method=hybrid&model=main) | SVM | NEGATIVE | -0.38842866 |
| 210 | 151-166 | [MPSLTLACLGPGPGV](http://crdd.osdd.net/raghava/ifnepitope/pep_design.php?sequence=MPSLTLACLGPGPGV&method=hybrid&model=main) | SVM | NEGATIVE | -0.16602537 |
| 211 | 152-167 | [PSLTLACLGPGPGVW](http://crdd.osdd.net/raghava/ifnepitope/pep_design.php?sequence=PSLTLACLGPGPGVW&method=hybrid&model=main) | SVM | NEGATIVE | -0.14563912 |
| 212 | 153-168 | [SLTLACLGPGPGVWD](http://crdd.osdd.net/raghava/ifnepitope/pep_design.php?sequence=SLTLACLGPGPGVWD&method=hybrid&model=main) | SVM | NEGATIVE | -0.45820579 |
| 213 | 154-169 | [LTLACLGPGPGVWDV](http://crdd.osdd.net/raghava/ifnepitope/pep_design.php?sequence=LTLACLGPGPGVWDV&method=hybrid&model=main) | SVM | NEGATIVE | -0.37802932 |
| 214 | 155-170 | [TLACLGPGPGVWDVK](http://crdd.osdd.net/raghava/ifnepitope/pep_design.php?sequence=TLACLGPGPGVWDVK&method=hybrid&model=main) | SVM | NEGATIVE | -0.5063164 |
| 215 | 156-171 | [LACLGPGPGVWDVKD](http://crdd.osdd.net/raghava/ifnepitope/pep_design.php?sequence=LACLGPGPGVWDVKD&method=hybrid&model=main) | SVM | NEGATIVE | -0.9180339 |
| 216 | 157-172 | [ACLGPGPGVWDVKDS](http://crdd.osdd.net/raghava/ifnepitope/pep_design.php?sequence=ACLGPGPGVWDVKDS&method=hybrid&model=main) | SVM | NEGATIVE | -0.93657397 |
| 217 | 158-173 | [CLGPGPGVWDVKDSS](http://crdd.osdd.net/raghava/ifnepitope/pep_design.php?sequence=CLGPGPGVWDVKDSS&method=hybrid&model=main) | SVM | NEGATIVE | -0.98100899 |
| 218 | 159-174 | [LGPGPGVWDVKDSSL](http://crdd.osdd.net/raghava/ifnepitope/pep_design.php?sequence=LGPGPGVWDVKDSSL&method=hybrid&model=main) | SVM | NEGATIVE | -0.89917096 |
| 219 | 160-175 | [GPGPGVWDVKDSSLL](http://crdd.osdd.net/raghava/ifnepitope/pep_design.php?sequence=GPGPGVWDVKDSSLL&method=hybrid&model=main) | SVM | NEGATIVE | -0.78916948 |
| 220 | 161-176 | [PGPGVWDVKDSSLLN](http://crdd.osdd.net/raghava/ifnepitope/pep_design.php?sequence=PGPGVWDVKDSSLLN&method=hybrid&model=main) | SVM | NEGATIVE | -0.82096238 |
| 221 | 162-177 | [GPGVWDVKDSSLLNN](http://crdd.osdd.net/raghava/ifnepitope/pep_design.php?sequence=GPGVWDVKDSSLLNN&method=hybrid&model=main) | SVM | NEGATIVE | -1.1533179 |
| 222 | 163-178 | [PGVWDVKDSSLLNNQ](http://crdd.osdd.net/raghava/ifnepitope/pep_design.php?sequence=PGVWDVKDSSLLNNQ&method=hybrid&model=main) | SVM | NEGATIVE | -1.0111686 |
| 223 | 164-179 | [GVWDVKDSSLLNNQF](http://crdd.osdd.net/raghava/ifnepitope/pep_design.php?sequence=GVWDVKDSSLLNNQF&method=hybrid&model=main) | SVM | NEGATIVE | -1.0856002 |
| 224 | 165-180 | [VWDVKDSSLLNNQFG](http://crdd.osdd.net/raghava/ifnepitope/pep_design.php?sequence=VWDVKDSSLLNNQFG&method=hybrid&model=main) | SVM | NEGATIVE | -1.3050832 |
| 225 | 166-181 | [WDVKDSSLLNNQFGG](http://crdd.osdd.net/raghava/ifnepitope/pep_design.php?sequence=WDVKDSSLLNNQFGG&method=hybrid&model=main) | SVM | NEGATIVE | -1.0893547 |
| 226 | 167-182 | [DVKDSSLLNNQFGGP](http://crdd.osdd.net/raghava/ifnepitope/pep_design.php?sequence=DVKDSSLLNNQFGGP&method=hybrid&model=main) | SVM | NEGATIVE | -1.1860253 |
| 227 | 168-183 | [VKDSSLLNNQFGGPG](http://crdd.osdd.net/raghava/ifnepitope/pep_design.php?sequence=VKDSSLLNNQFGGPG&method=hybrid&model=main) | SVM | NEGATIVE | -1.1902899 |
| 228 | 169-184 | [KDSSLLNNQFGGPGP](http://crdd.osdd.net/raghava/ifnepitope/pep_design.php?sequence=KDSSLLNNQFGGPGP&method=hybrid&model=main) | SVM | NEGATIVE | -1.2574122 |
| 229 | 170-185 | [DSSLLNNQFGGPGPG](http://crdd.osdd.net/raghava/ifnepitope/pep_design.php?sequence=DSSLLNNQFGGPGPG&method=hybrid&model=main) | SVM | NEGATIVE | -0.85763309 |
| 230 | 171-186 | [SSLLNNQFGGPGPGL](http://crdd.osdd.net/raghava/ifnepitope/pep_design.php?sequence=SSLLNNQFGGPGPGL&method=hybrid&model=main) | SVM | NEGATIVE | -0.67634606 |
| 231 | 172-187 | [SLLNNQFGGPGPGLT](http://crdd.osdd.net/raghava/ifnepitope/pep_design.php?sequence=SLLNNQFGGPGPGLT&method=hybrid&model=main) | SVM | NEGATIVE | -0.38719531 |
| 232 | 173-188 | [LLNNQFGGPGPGLTS](http://crdd.osdd.net/raghava/ifnepitope/pep_design.php?sequence=LLNNQFGGPGPGLTS&method=hybrid&model=main) | SVM | NEGATIVE | -0.4258004 |
| 233 | 174-189 | [LNNQFGGPGPGLTSL](http://crdd.osdd.net/raghava/ifnepitope/pep_design.php?sequence=LNNQFGGPGPGLTSL&method=hybrid&model=main) | SVM | NEGATIVE | -0.43241951 |
| 234 | 175-190 | [NNQFGGPGPGLTSLG](http://crdd.osdd.net/raghava/ifnepitope/pep_design.php?sequence=NNQFGGPGPGLTSLG&method=hybrid&model=main) | SVM | NEGATIVE | -0.52081835 |
| 235 | 176-191 | [NQFGGPGPGLTSLGL](http://crdd.osdd.net/raghava/ifnepitope/pep_design.php?sequence=NQFGGPGPGLTSLGL&method=hybrid&model=main) | SVM | NEGATIVE | -0.2463646 |
| 236 | 177-192 | [QFGGPGPGLTSLGLL](http://crdd.osdd.net/raghava/ifnepitope/pep_design.php?sequence=QFGGPGPGLTSLGLL&method=hybrid&model=main) | SVM | NEGATIVE | -0.030639337 |
| 237 | 178-193 | [FGGPGPGLTSLGLLY](http://crdd.osdd.net/raghava/ifnepitope/pep_design.php?sequence=FGGPGPGLTSLGLLY&method=hybrid&model=main) | SVM | POSITIVE | 0.085797731 |
| 238 | 179-194 | [GGPGPGLTSLGLLYT](http://crdd.osdd.net/raghava/ifnepitope/pep_design.php?sequence=GGPGPGLTSLGLLYT&method=hybrid&model=main) | SVM | POSITIVE | 0.3422808 |
| 239 | 188-203 | [LGLLYTVKYPNLKKS](http://crdd.osdd.net/raghava/ifnepitope/pep_design.php?sequence=LGLLYTVKYPNLKKS&method=hybrid&model=main) | SVM | POSITIVE | 0.3651326 |
| 240 | 189-204 | [GLLYTVKYPNLKKSE](http://crdd.osdd.net/raghava/ifnepitope/pep_design.php?sequence=GLLYTVKYPNLKKSE&method=hybrid&model=main) | SVM | NEGATIVE | -0.14041277 |
| 241 | 190-205 | [LLYTVKYPNLKKSER](http://crdd.osdd.net/raghava/ifnepitope/pep_design.php?sequence=LLYTVKYPNLKKSER&method=hybrid&model=main) | SVM | NEGATIVE | -0.70875788 |
| 242 | 191-206 | [LYTVKYPNLKKSERP](http://crdd.osdd.net/raghava/ifnepitope/pep_design.php?sequence=LYTVKYPNLKKSERP&method=hybrid&model=main) | SVM | NEGATIVE | -0.49938045 |
| 243 | 192-207 | [YTVKYPNLKKSERPQ](http://crdd.osdd.net/raghava/ifnepitope/pep_design.php?sequence=YTVKYPNLKKSERPQ&method=hybrid&model=main) | SVM | NEGATIVE | -0.83741974 |
| 244 | 193-208 | [TVKYPNLKKSERPQA](http://crdd.osdd.net/raghava/ifnepitope/pep_design.php?sequence=TVKYPNLKKSERPQA&method=hybrid&model=main) | SVM | NEGATIVE | -0.56201175 |
| 245 | 194-209 | [VKYPNLKKSERPQAS](http://crdd.osdd.net/raghava/ifnepitope/pep_design.php?sequence=VKYPNLKKSERPQAS&method=hybrid&model=main) | SVM | NEGATIVE | -0.57850041 |
| 246 | 195-210 | [KYPNLKKSERPQASG](http://crdd.osdd.net/raghava/ifnepitope/pep_design.php?sequence=KYPNLKKSERPQASG&method=hybrid&model=main) | SVM | NEGATIVE | -0.48208871 |
| 247 | 196-211 | [YPNLKKSERPQASGV](http://crdd.osdd.net/raghava/ifnepitope/pep_design.php?sequence=YPNLKKSERPQASGV&method=hybrid&model=main) | SVM | NEGATIVE | -0.20328235 |
| 248 | 197-212 | [PNLKKSERPQASGVK](http://crdd.osdd.net/raghava/ifnepitope/pep_design.php?sequence=PNLKKSERPQASGVK&method=hybrid&model=main) | SVM | NEGATIVE | -0.16561613 |
| 249 | 198-213 | [NLKKSERPQASGVKK](http://crdd.osdd.net/raghava/ifnepitope/pep_design.php?sequence=NLKKSERPQASGVKK&method=hybrid&model=main) | SVM | NEGATIVE | -0.21516333 |
| 250 | 199-214 | [LKKSERPQASGVKKI](http://crdd.osdd.net/raghava/ifnepitope/pep_design.php?sequence=LKKSERPQASGVKKI&method=hybrid&model=main) | SVM | NEGATIVE | -0.33209764 |
| 251 | 200-215 | [KKSERPQASGVKKIM](http://crdd.osdd.net/raghava/ifnepitope/pep_design.php?sequence=KKSERPQASGVKKIM&method=hybrid&model=main) | SVM | NEGATIVE | -0.41645806 |
| 252 | 201-216 | [KSERPQASGVKKIMR](http://crdd.osdd.net/raghava/ifnepitope/pep_design.php?sequence=KSERPQASGVKKIMR&method=hybrid&model=main) | SVM | NEGATIVE | -0.67040274 |
| 253 | 202-217 | [SERPQASGVKKIMRK](http://crdd.osdd.net/raghava/ifnepitope/pep_design.php?sequence=SERPQASGVKKIMRK&method=hybrid&model=main) | SVM | NEGATIVE | -0.65483794 |
| 254 | 203-218 | [ERPQASGVKKIMRKE](http://crdd.osdd.net/raghava/ifnepitope/pep_design.php?sequence=ERPQASGVKKIMRKE&method=hybrid&model=main) | SVM | NEGATIVE | -0.65843342 |
| 255 | 204-219 | [RPQASGVKKIMRKEK](http://crdd.osdd.net/raghava/ifnepitope/pep_design.php?sequence=RPQASGVKKIMRKEK&method=hybrid&model=main) | SVM | NEGATIVE | -0.30360993 |
| 256 | 205-220 | [PQASGVKKIMRKEKR](http://crdd.osdd.net/raghava/ifnepitope/pep_design.php?sequence=PQASGVKKIMRKEKR&method=hybrid&model=main) | SVM | NEGATIVE | -0.49531914 |
| 257 | 206-221 | [QASGVKKIMRKEKRD](http://crdd.osdd.net/raghava/ifnepitope/pep_design.php?sequence=QASGVKKIMRKEKRD&method=hybrid&model=main) | SVM | NEGATIVE | -0.50801916 |
| 258 | 207-222 | [ASGVKKIMRKEKRDD](http://crdd.osdd.net/raghava/ifnepitope/pep_design.php?sequence=ASGVKKIMRKEKRDD&method=hybrid&model=main) | SVM | NEGATIVE | -0.77950732 |
| 259 | 208-223 | [SGVKKIMRKEKRDDK](http://crdd.osdd.net/raghava/ifnepitope/pep_design.php?sequence=SGVKKIMRKEKRDDK&method=hybrid&model=main) | SVM | NEGATIVE | -0.79799186 |
| 260 | 209-224 | [GVKKIMRKEKRDDKD](http://crdd.osdd.net/raghava/ifnepitope/pep_design.php?sequence=GVKKIMRKEKRDDKD&method=hybrid&model=main) | SVM | NEGATIVE | -1.0143722 |
| 261 | 210-225 | [VKKIMRKEKRDDKDL](http://crdd.osdd.net/raghava/ifnepitope/pep_design.php?sequence=VKKIMRKEKRDDKDL&method=hybrid&model=main) | SVM | NEGATIVE | -1.0068656 |
| 262 | 211-226 | [KKIMRKEKRDDKDLQ](http://crdd.osdd.net/raghava/ifnepitope/pep_design.php?sequence=KKIMRKEKRDDKDLQ&method=hybrid&model=main) | SVM | NEGATIVE | -1.1358756 |
| 263 | 212-227 | [KIMRKEKRDDKDLQR](http://crdd.osdd.net/raghava/ifnepitope/pep_design.php?sequence=KIMRKEKRDDKDLQR&method=hybrid&model=main) | SVM | NEGATIVE | -0.88827454 |
| 264 | 213-228 | [IMRKEKRDDKDLQRL](http://crdd.osdd.net/raghava/ifnepitope/pep_design.php?sequence=IMRKEKRDDKDLQRL&method=hybrid&model=main) | SVM | NEGATIVE | -0.81481995 |
| 265 | 214-229 | [MRKEKRDDKDLQRLR](http://crdd.osdd.net/raghava/ifnepitope/pep_design.php?sequence=MRKEKRDDKDLQRLR&method=hybrid&model=main) | SVM | NEGATIVE | -0.68274153 |
| 266 | 215-230 | [RKEKRDDKDLQRLRK](http://crdd.osdd.net/raghava/ifnepitope/pep_design.php?sequence=RKEKRDDKDLQRLRK&method=hybrid&model=main) | SVM | NEGATIVE | -0.21101486 |
| 267 | 216-231 | [KEKRDDKDLQRLRKK](http://crdd.osdd.net/raghava/ifnepitope/pep_design.php?sequence=KEKRDDKDLQRLRKK&method=hybrid&model=main) | SVM | NEGATIVE | -0.55956706 |
| 268 | 217-232 | [EKRDDKDLQRLRKKP](http://crdd.osdd.net/raghava/ifnepitope/pep_design.php?sequence=EKRDDKDLQRLRKKP&method=hybrid&model=main) | SVM | NEGATIVE | -0.34204297 |
| 269 | 218-233 | [KRDDKDLQRLRKKPP](http://crdd.osdd.net/raghava/ifnepitope/pep_design.php?sequence=KRDDKDLQRLRKKPP&method=hybrid&model=main) | SVM | NEGATIVE | -0.53242643 |
| 270 | 219-234 | [RDDKDLQRLRKKPPQ](http://crdd.osdd.net/raghava/ifnepitope/pep_design.php?sequence=RDDKDLQRLRKKPPQ&method=hybrid&model=main) | SVM | NEGATIVE | -0.86546016 |
| 271 | 220-235 | [DDKDLQRLRKKPPQV](http://crdd.osdd.net/raghava/ifnepitope/pep_design.php?sequence=DDKDLQRLRKKPPQV&method=hybrid&model=main) | SVM | NEGATIVE | -0.69551984 |
| 272 | 221-236 | [DKDLQRLRKKPPQVG](http://crdd.osdd.net/raghava/ifnepitope/pep_design.php?sequence=DKDLQRLRKKPPQVG&method=hybrid&model=main) | SVM | NEGATIVE | -0.56222148 |
| 273 | 222-237 | [KDLQRLRKKPPQVGL](http://crdd.osdd.net/raghava/ifnepitope/pep_design.php?sequence=KDLQRLRKKPPQVGL&method=hybrid&model=main) | SVM | NEGATIVE | -0.23643545 |
| 274 | 223-238 | [DLQRLRKKPPQVGLS](http://crdd.osdd.net/raghava/ifnepitope/pep_design.php?sequence=DLQRLRKKPPQVGLS&method=hybrid&model=main) | SVM | NEGATIVE | -0.25040527 |
| 275 | 224-239 | [LQRLRKKPPQVGLSY](http://crdd.osdd.net/raghava/ifnepitope/pep_design.php?sequence=LQRLRKKPPQVGLSY&method=hybrid&model=main) | SVM | NEGATIVE | -0.23637785 |
| 276 | 225-240 | [QRLRKKPPQVGLSYS](http://crdd.osdd.net/raghava/ifnepitope/pep_design.php?sequence=QRLRKKPPQVGLSYS&method=hybrid&model=main) | SVM | NEGATIVE | -0.34916935 |
| 277 | 226-241 | [RLRKKPPQVGLSYSQ](http://crdd.osdd.net/raghava/ifnepitope/pep_design.php?sequence=RLRKKPPQVGLSYSQ&method=hybrid&model=main) | SVM | NEGATIVE | -0.59201315 |
| 278 | 227-242 | [LRKKPPQVGLSYSQT](http://crdd.osdd.net/raghava/ifnepitope/pep_design.php?sequence=LRKKPPQVGLSYSQT&method=hybrid&model=main) | SVM | NEGATIVE | -0.62110986 |
| 279 | 228-243 | [RKKPPQVGLSYSQTM](http://crdd.osdd.net/raghava/ifnepitope/pep_design.php?sequence=RKKPPQVGLSYSQTM&method=hybrid&model=main) | SVM | NEGATIVE | -0.5665652 |
| 280 | 229-244 | [KKPPQVGLSYSQTMK](http://crdd.osdd.net/raghava/ifnepitope/pep_design.php?sequence=KKPPQVGLSYSQTMK&method=hybrid&model=main) | SVM | NEGATIVE | -0.46780896 |
| 281 | 230-245 | [KPPQVGLSYSQTMKK](http://crdd.osdd.net/raghava/ifnepitope/pep_design.php?sequence=KPPQVGLSYSQTMKK&method=hybrid&model=main) | SVM | NEGATIVE | -0.46780896 |
| 282 | 231-246 | [PPQVGLSYSQTMKKP](http://crdd.osdd.net/raghava/ifnepitope/pep_design.php?sequence=PPQVGLSYSQTMKKP&method=hybrid&model=main) | SVM | NEGATIVE | -0.46780896 |
| 283 | 232-247 | [PQVGLSYSQTMKKPN](http://crdd.osdd.net/raghava/ifnepitope/pep_design.php?sequence=PQVGLSYSQTMKKPN&method=hybrid&model=main) | SVM | NEGATIVE | -0.16947114 |
| 284 | 233-248 | [QVGLSYSQTMKKPNA](http://crdd.osdd.net/raghava/ifnepitope/pep_design.php?sequence=QVGLSYSQTMKKPNA&method=hybrid&model=main) | SVM | POSITIVE | 0.085972778 |
| 285 | 234-249 | [VGLSYSQTMKKPNAK](http://crdd.osdd.net/raghava/ifnepitope/pep_design.php?sequence=VGLSYSQTMKKPNAK&method=hybrid&model=main) | SVM | POSITIVE | 0.15417348 |
| 286 | 235-250 | [GLSYSQTMKKPNAKT](http://crdd.osdd.net/raghava/ifnepitope/pep_design.php?sequence=GLSYSQTMKKPNAKT&method=hybrid&model=main) | SVM | POSITIVE | 0.073374005 |
| 287 | 236-251 | [LSYSQTMKKPNAKTW](http://crdd.osdd.net/raghava/ifnepitope/pep_design.php?sequence=LSYSQTMKKPNAKTW&method=hybrid&model=main) | SVM | NEGATIVE | -0.32920102 |
| 288 | 237-252 | [SYSQTMKKPNAKTWM](http://crdd.osdd.net/raghava/ifnepitope/pep_design.php?sequence=SYSQTMKKPNAKTWM&method=hybrid&model=main) | SVM | NEGATIVE | -0.27281234 |
| 289 | 238-253 | [YSQTMKKPNAKTWMD](http://crdd.osdd.net/raghava/ifnepitope/pep_design.php?sequence=YSQTMKKPNAKTWMD&method=hybrid&model=main) | SVM | NEGATIVE | -0.42213072 |
| 290 | 239-254 | [SQTMKKPNAKTWMDI](http://crdd.osdd.net/raghava/ifnepitope/pep_design.php?sequence=SQTMKKPNAKTWMDI&method=hybrid&model=main) | SVM | NEGATIVE | -0.42051117 |
| 291 | 240-255 | [QTMKKPNAKTWMDIE](http://crdd.osdd.net/raghava/ifnepitope/pep_design.php?sequence=QTMKKPNAKTWMDIE&method=hybrid&model=main) | SVM | NEGATIVE | -0.59674498 |
| 292 | 241-256 | [TMKKPNAKTWMDIEG](http://crdd.osdd.net/raghava/ifnepitope/pep_design.php?sequence=TMKKPNAKTWMDIEG&method=hybrid&model=main) | SVM | NEGATIVE | -0.428234 |
| 293 | 242-257 | [MKKPNAKTWMDIEGR](http://crdd.osdd.net/raghava/ifnepitope/pep_design.php?sequence=MKKPNAKTWMDIEGR&method=hybrid&model=main) | SVM | NEGATIVE | -0.49229376 |
| 294 | 243-258 | [KKPNAKTWMDIEGRP](http://crdd.osdd.net/raghava/ifnepitope/pep_design.php?sequence=KKPNAKTWMDIEGRP&method=hybrid&model=main) | SVM | NEGATIVE | -0.10425893 |
| 295 | 244-259 | [KPNAKTWMDIEGRPE](http://crdd.osdd.net/raghava/ifnepitope/pep_design.php?sequence=KPNAKTWMDIEGRPE&method=hybrid&model=main) | SVM | NEGATIVE | -0.18122653 |
| 296 | 245-260 | [PNAKTWMDIEGRPED](http://crdd.osdd.net/raghava/ifnepitope/pep_design.php?sequence=PNAKTWMDIEGRPED&method=hybrid&model=main) | SVM | NEGATIVE | -0.13863731 |
| 297 | 246-261 | [NAKTWMDIEGRPEDP](http://crdd.osdd.net/raghava/ifnepitope/pep_design.php?sequence=NAKTWMDIEGRPEDP&method=hybrid&model=main) | SVM | NEGATIVE | -0.10307564 |
| 298 | 247-262 | [AKTWMDIEGRPEDPV](http://crdd.osdd.net/raghava/ifnepitope/pep_design.php?sequence=AKTWMDIEGRPEDPV&method=hybrid&model=main) | SVM | POSITIVE | 0.0090228769 |
| 299 | 248-263 | [KTWMDIEGRPEDPVE](http://crdd.osdd.net/raghava/ifnepitope/pep_design.php?sequence=KTWMDIEGRPEDPVE&method=hybrid&model=main) | SVM | NEGATIVE | -0.0034733879 |
| 300 | 249-264 | [TWMDIEGRPEDPVEK](http://crdd.osdd.net/raghava/ifnepitope/pep_design.php?sequence=TWMDIEGRPEDPVEK&method=hybrid&model=main) | SVM | POSITIVE | 0.23372861 |
| 301 | 250-265 | [WMDIEGRPEDPVEKK](http://crdd.osdd.net/raghava/ifnepitope/pep_design.php?sequence=WMDIEGRPEDPVEKK&method=hybrid&model=main) | SVM | POSITIVE | 0.43524288 |
| 302 | 251-266 | [MDIEGRPEDPVEKKL](http://crdd.osdd.net/raghava/ifnepitope/pep_design.php?sequence=MDIEGRPEDPVEKKL&method=hybrid&model=main) | SVM | POSITIVE | 0.61835908 |
| 303 | 252-267 | [DIEGRPEDPVEKKLM](http://crdd.osdd.net/raghava/ifnepitope/pep_design.php?sequence=DIEGRPEDPVEKKLM&method=hybrid&model=main) | SVM | POSITIVE | 0.72950255 |
| 304 | 253-268 | [IEGRPEDPVEKKLMR](http://crdd.osdd.net/raghava/ifnepitope/pep_design.php?sequence=IEGRPEDPVEKKLMR&method=hybrid&model=main) | SVM | POSITIVE | 0.64375623 |
| 305 | 254-269 | [EGRPEDPVEKKLMRK](http://crdd.osdd.net/raghava/ifnepitope/pep_design.php?sequence=EGRPEDPVEKKLMRK&method=hybrid&model=main) | SVM | POSITIVE | 0.81888234 |
| 306 | 255-270 | [GRPEDPVEKKLMRKE](http://crdd.osdd.net/raghava/ifnepitope/pep_design.php?sequence=GRPEDPVEKKLMRKE&method=hybrid&model=main) | SVM | POSITIVE | 0.45967125 |
| 307 | 256-271 | [RPEDPVEKKLMRKER](http://crdd.osdd.net/raghava/ifnepitope/pep_design.php?sequence=RPEDPVEKKLMRKER&method=hybrid&model=main) | SVM | POSITIVE | 0.16687365 |
| 308 | 257-272 | [PEDPVEKKLMRKERR](http://crdd.osdd.net/raghava/ifnepitope/pep_design.php?sequence=PEDPVEKKLMRKERR&method=hybrid&model=main) | SVM | NEGATIVE | -0.31402559 |
| 309 | 258-273 | [EDPVEKKLMRKERRD](http://crdd.osdd.net/raghava/ifnepitope/pep_design.php?sequence=EDPVEKKLMRKERRD&method=hybrid&model=main) | SVM | NEGATIVE | -0.43285653 |
| 310 | 259-274 | [DPVEKKLMRKERRDD](http://crdd.osdd.net/raghava/ifnepitope/pep_design.php?sequence=DPVEKKLMRKERRDD&method=hybrid&model=main) | SVM | NEGATIVE | -0.526048 |
| 311 | 260-275 | [PVEKKLMRKERRDDN](http://crdd.osdd.net/raghava/ifnepitope/pep_design.php?sequence=PVEKKLMRKERRDDN&method=hybrid&model=main) | SVM | NEGATIVE | -0.47709451 |
| 312 | 261-276 | [VEKKLMRKERRDDND](http://crdd.osdd.net/raghava/ifnepitope/pep_design.php?sequence=VEKKLMRKERRDDND&method=hybrid&model=main) | SVM | NEGATIVE | -0.69568393 |
| 313 | 262-277 | [EKKLMRKERRDDNDL](http://crdd.osdd.net/raghava/ifnepitope/pep_design.php?sequence=EKKLMRKERRDDNDL&method=hybrid&model=main) | SVM | NEGATIVE | -0.83074765 |
| 314 | 263-278 | [KKLMRKERRDDNDLK](http://crdd.osdd.net/raghava/ifnepitope/pep_design.php?sequence=KKLMRKERRDDNDLK&method=hybrid&model=main) | SVM | NEGATIVE | -0.96193051 |
| 315 | 264-279 | [KLMRKERRDDNDLKR](http://crdd.osdd.net/raghava/ifnepitope/pep_design.php?sequence=KLMRKERRDDNDLKR&method=hybrid&model=main) | SVM | NEGATIVE | -1.0362245 |
| 316 | 265-280 | [LMRKERRDDNDLKRL](http://crdd.osdd.net/raghava/ifnepitope/pep_design.php?sequence=LMRKERRDDNDLKRL&method=hybrid&model=main) | SVM | NEGATIVE | -0.92317743 |
| 317 | 266-281 | [MRKERRDDNDLKRLR](http://crdd.osdd.net/raghava/ifnepitope/pep_design.php?sequence=MRKERRDDNDLKRLR&method=hybrid&model=main) | SVM | NEGATIVE | -0.91188679 |
| 318 | 267-282 | [RKERRDDNDLKRLRD](http://crdd.osdd.net/raghava/ifnepitope/pep_design.php?sequence=RKERRDDNDLKRLRD&method=hybrid&model=main) | SVM | NEGATIVE | -0.55149143 |
| 319 | 268-283 | [KERRDDNDLKRLRDL](http://crdd.osdd.net/raghava/ifnepitope/pep_design.php?sequence=KERRDDNDLKRLRDL&method=hybrid&model=main) | SVM | NEGATIVE | -0.58599414 |
| 320 | 269-284 | [ERRDDNDLKRLRDLK](http://crdd.osdd.net/raghava/ifnepitope/pep_design.php?sequence=ERRDDNDLKRLRDLK&method=hybrid&model=main) | SVM | NEGATIVE | -0.20670896 |
| 321 | 270-285 | [RRDDNDLKRLRDLKK](http://crdd.osdd.net/raghava/ifnepitope/pep_design.php?sequence=RRDDNDLKRLRDLKK&method=hybrid&model=main) | SVM | NEGATIVE | -0.031762181 |
| 322 | 271-286 | [RDDNDLKRLRDLKKS](http://crdd.osdd.net/raghava/ifnepitope/pep_design.php?sequence=RDDNDLKRLRDLKKS&method=hybrid&model=main) | SVM | NEGATIVE | -0.10086332 |
| 323 | 272-287 | [DDNDLKRLRDLKKSL](http://crdd.osdd.net/raghava/ifnepitope/pep_design.php?sequence=DDNDLKRLRDLKKSL&method=hybrid&model=main) | SVM | NEGATIVE | -0.2875342 |
| 324 | 273-288 | [DNDLKRLRDLKKSLT](http://crdd.osdd.net/raghava/ifnepitope/pep_design.php?sequence=DNDLKRLRDLKKSLT&method=hybrid&model=main) | SVM | NEGATIVE | -0.065685604 |
| 325 | 274-289 | [NDLKRLRDLKKSLTL](http://crdd.osdd.net/raghava/ifnepitope/pep_design.php?sequence=NDLKRLRDLKKSLTL&method=hybrid&model=main) | SVM | POSITIVE | 0.0092401936 |
| 326 | 275-290 | [DLKRLRDLKKSLTLA](http://crdd.osdd.net/raghava/ifnepitope/pep_design.php?sequence=DLKRLRDLKKSLTLA&method=hybrid&model=main) | SVM | NEGATIVE | -0.08950424 |
| 327 | 276-291 | [LKRLRDLKKSLTLAC](http://crdd.osdd.net/raghava/ifnepitope/pep_design.php?sequence=LKRLRDLKKSLTLAC&method=hybrid&model=main) | SVM | NEGATIVE | -0.22798343 |
| 328 | 277-292 | [KRLRDLKKSLTLACL](http://crdd.osdd.net/raghava/ifnepitope/pep_design.php?sequence=KRLRDLKKSLTLACL&method=hybrid&model=main) | SVM | NEGATIVE | -0.61905293 |
| 329 | 278-293 | [RLRDLKKSLTLACLT](http://crdd.osdd.net/raghava/ifnepitope/pep_design.php?sequence=RLRDLKKSLTLACLT&method=hybrid&model=main) | SVM | NEGATIVE | -0.55332507 |
| 330 | 279-294 | [LRDLKKSLTLACLTK](http://crdd.osdd.net/raghava/ifnepitope/pep_design.php?sequence=LRDLKKSLTLACLTK&method=hybrid&model=main) | SVM | NEGATIVE | -0.66502019 |
| 331 | 280-295 | [RDLKKSLTLACLTKQ](http://crdd.osdd.net/raghava/ifnepitope/pep_design.php?sequence=RDLKKSLTLACLTKQ&method=hybrid&model=main) | SVM | NEGATIVE | -0.51526692 |
| 332 | 281-296 | [DLKKSLTLACLTKQK](http://crdd.osdd.net/raghava/ifnepitope/pep_design.php?sequence=DLKKSLTLACLTKQK&method=hybrid&model=main) | SVM | NEGATIVE | -0.45110138 |
| 333 | 282-297 | [LKKSLTLACLTKQKK](http://crdd.osdd.net/raghava/ifnepitope/pep_design.php?sequence=LKKSLTLACLTKQKK&method=hybrid&model=main) | SVM | NEGATIVE | -0.19697109 |
| 334 | 283-298 | [KKSLTLACLTKQKKK](http://crdd.osdd.net/raghava/ifnepitope/pep_design.php?sequence=KKSLTLACLTKQKKK&method=hybrid&model=main) | SVM | POSITIVE | 0.095193632 |
| 335 | 284-299 | [KSLTLACLTKQKKKV](http://crdd.osdd.net/raghava/ifnepitope/pep_design.php?sequence=KSLTLACLTKQKKKV&method=hybrid&model=main) | SVM | NEGATIVE | -0.3219944 |
| 336 | 285-300 | [SLTLACLTKQKKKVV](http://crdd.osdd.net/raghava/ifnepitope/pep_design.php?sequence=SLTLACLTKQKKKVV&method=hybrid&model=main) | SVM | NEGATIVE | -0.21138115 |
| 337 | 286-301 | [LTLACLTKQKKKVVK](http://crdd.osdd.net/raghava/ifnepitope/pep_design.php?sequence=LTLACLTKQKKKVVK&method=hybrid&model=main) | SVM | NEGATIVE | -0.28751111 |
| 338 | 287-302 | [TLACLTKQKKKVVKD](http://crdd.osdd.net/raghava/ifnepitope/pep_design.php?sequence=TLACLTKQKKKVVKD&method=hybrid&model=main) | SVM | NEGATIVE | -0.64503783 |
| 339 | 288-303 | [LACLTKQKKKVVKDA](http://crdd.osdd.net/raghava/ifnepitope/pep_design.php?sequence=LACLTKQKKKVVKDA&method=hybrid&model=main) | SVM | NEGATIVE | -0.53165729 |
| 340 | 297-312 | [KVVKDAVSLINGLDF](http://crdd.osdd.net/raghava/ifnepitope/pep_design.php?sequence=KVVKDAVSLINGLDF&method=hybrid&model=main) | SVM | NEGATIVE | -0.078536871 |
| 341 | 298-313 | [VVKDAVSLINGLDFS](http://crdd.osdd.net/raghava/ifnepitope/pep_design.php?sequence=VVKDAVSLINGLDFS&method=hybrid&model=main) | SVM | POSITIVE | 0.028658313 |
| 342 | 299-314 | [VKDAVSLINGLDFSM](http://crdd.osdd.net/raghava/ifnepitope/pep_design.php?sequence=VKDAVSLINGLDFSM&method=hybrid&model=main) | SVM | NEGATIVE | -0.34721461 |
| 343 | 300-315 | [KDAVSLINGLDFSMV](http://crdd.osdd.net/raghava/ifnepitope/pep_design.php?sequence=KDAVSLINGLDFSMV&method=hybrid&model=main) | SVM | NEGATIVE | -0.53887315 |
| 344 | 301-316 | [DAVSLINGLDFSMVK](http://crdd.osdd.net/raghava/ifnepitope/pep_design.php?sequence=DAVSLINGLDFSMVK&method=hybrid&model=main) | SVM | NEGATIVE | -0.47234307 |
| 345 | 302-317 | [AVSLINGLDFSMVKK](http://crdd.osdd.net/raghava/ifnepitope/pep_design.php?sequence=AVSLINGLDFSMVKK&method=hybrid&model=main) | SVM | NEGATIVE | -0.48331243 |
| 346 | 303-318 | [VSLINGLDFSMVKKP](http://crdd.osdd.net/raghava/ifnepitope/pep_design.php?sequence=VSLINGLDFSMVKKP&method=hybrid&model=main) | SVM | NEGATIVE | -0.75396695 |
| 347 | 304-319 | [SLINGLDFSMVKKPN](http://crdd.osdd.net/raghava/ifnepitope/pep_design.php?sequence=SLINGLDFSMVKKPN&method=hybrid&model=main) | SVM | NEGATIVE | -0.69015274 |
| 348 | 305-320 | [LINGLDFSMVKKPNM](http://crdd.osdd.net/raghava/ifnepitope/pep_design.php?sequence=LINGLDFSMVKKPNM&method=hybrid&model=main) | SVM | NEGATIVE | -0.70430015 |
| 349 | 306-321 | [INGLDFSMVKKPNMD](http://crdd.osdd.net/raghava/ifnepitope/pep_design.php?sequence=INGLDFSMVKKPNMD&method=hybrid&model=main) | SVM | NEGATIVE | -0.93381325 |
| 350 | 307-322 | [NGLDFSMVKKPNMDD](http://crdd.osdd.net/raghava/ifnepitope/pep_design.php?sequence=NGLDFSMVKKPNMDD&method=hybrid&model=main) | SVM | NEGATIVE | -0.97031073 |
| 351 | 308-323 | [GLDFSMVKKPNMDDL](http://crdd.osdd.net/raghava/ifnepitope/pep_design.php?sequence=GLDFSMVKKPNMDDL&method=hybrid&model=main) | SVM | NEGATIVE | -0.93269133 |
| 352 | 309-324 | [LDFSMVKKPNMDDLD](http://crdd.osdd.net/raghava/ifnepitope/pep_design.php?sequence=LDFSMVKKPNMDDLD&method=hybrid&model=main) | SVM | NEGATIVE | -1.0113018 |
| 353 | 310-325 | [DFSMVKKPNMDDLDK](http://crdd.osdd.net/raghava/ifnepitope/pep_design.php?sequence=DFSMVKKPNMDDLDK&method=hybrid&model=main) | SVM | NEGATIVE | -1.1340963 |
| 354 | 311-326 | [FSMVKKPNMDDLDKL](http://crdd.osdd.net/raghava/ifnepitope/pep_design.php?sequence=FSMVKKPNMDDLDKL&method=hybrid&model=main) | SVM | NEGATIVE | -1.2206639 |
| 355 | 312-327 | [SMVKKPNMDDLDKLK](http://crdd.osdd.net/raghava/ifnepitope/pep_design.php?sequence=SMVKKPNMDDLDKLK&method=hybrid&model=main) | SVM | NEGATIVE | -1.198548 |
| 356 | 313-328 | [MVKKPNMDDLDKLKN](http://crdd.osdd.net/raghava/ifnepitope/pep_design.php?sequence=MVKKPNMDDLDKLKN&method=hybrid&model=main) | SVM | NEGATIVE | -1.0729094 |
| 357 | 314-329 | [VKKPNMDDLDKLKNK](http://crdd.osdd.net/raghava/ifnepitope/pep_design.php?sequence=VKKPNMDDLDKLKNK&method=hybrid&model=main) | SVM | NEGATIVE | -0.83839835 |
| 358 | 315-330 | [KKPNMDDLDKLKNKK](http://crdd.osdd.net/raghava/ifnepitope/pep_design.php?sequence=KKPNMDDLDKLKNKK&method=hybrid&model=main) | SVM | NEGATIVE | -0.42674519 |
| 359 | 334-349 | [YKICLSGKKDERPGN](http://crdd.osdd.net/raghava/ifnepitope/pep_design.php?sequence=YKICLSGKKDERPGN&method=hybrid&model=main) | SVM | NEGATIVE | -0.31240167 |
| 360 | 335-350 | [KICLSGKKDERPGNR](http://crdd.osdd.net/raghava/ifnepitope/pep_design.php?sequence=KICLSGKKDERPGNR&method=hybrid&model=main) | SVM | NEGATIVE | -0.17148308 |
| 361 | 336-351 | [ICLSGKKDERPGNRN](http://crdd.osdd.net/raghava/ifnepitope/pep_design.php?sequence=ICLSGKKDERPGNRN&method=hybrid&model=main) | SVM | NEGATIVE | -0.22871153 |
| 362 | 337-352 | [CLSGKKDERPGNRNP](http://crdd.osdd.net/raghava/ifnepitope/pep_design.php?sequence=CLSGKKDERPGNRNP&method=hybrid&model=main) | SVM | NEGATIVE | -0.31276267 |
| 363 | 338-353 | [LSGKKDERPGNRNPY](http://crdd.osdd.net/raghava/ifnepitope/pep_design.php?sequence=LSGKKDERPGNRNPY&method=hybrid&model=main) | SVM | NEGATIVE | -0.49050976 |
| 364 | 339-354 | [SGKKDERPGNRNPYK](http://crdd.osdd.net/raghava/ifnepitope/pep_design.php?sequence=SGKKDERPGNRNPYK&method=hybrid&model=main) | SVM | NEGATIVE | -0.48674638 |
| 365 | 340-355 | [GKKDERPGNRNPYKK](http://crdd.osdd.net/raghava/ifnepitope/pep_design.php?sequence=GKKDERPGNRNPYKK&method=hybrid&model=main) | SVM | NEGATIVE | -0.23919071 |
| 366 | 341-356 | [KKDERPGNRNPYKKP](http://crdd.osdd.net/raghava/ifnepitope/pep_design.php?sequence=KKDERPGNRNPYKKP&method=hybrid&model=main) | SVM | NEGATIVE | -0.41038476 |
| 367 | 342-357 | [KDERPGNRNPYKKPG](http://crdd.osdd.net/raghava/ifnepitope/pep_design.php?sequence=KDERPGNRNPYKKPG&method=hybrid&model=main) | SVM | NEGATIVE | -0.14387479 |
| 368 | 343-358 | [DERPGNRNPYKKPGL](http://crdd.osdd.net/raghava/ifnepitope/pep_design.php?sequence=DERPGNRNPYKKPGL&method=hybrid&model=main) | SVM | POSITIVE | 0.075038335 |
| 369 | 344-359 | [ERPGNRNPYKKPGLL](http://crdd.osdd.net/raghava/ifnepitope/pep_design.php?sequence=ERPGNRNPYKKPGLL&method=hybrid&model=main) | SVM | POSITIVE | 0.10493205 |
| 370 | 345-360 | [RPGNRNPYKKPGLLS](http://crdd.osdd.net/raghava/ifnepitope/pep_design.php?sequence=RPGNRNPYKKPGLLS&method=hybrid&model=main) | SVM | POSITIVE | 0.24442423 |
| 371 | 346-361 | [PGNRNPYKKPGLLSY](http://crdd.osdd.net/raghava/ifnepitope/pep_design.php?sequence=PGNRNPYKKPGLLSY&method=hybrid&model=main) | SVM | NEGATIVE | -0.20721056 |
| 372 | 347-362 | [GNRNPYKKPGLLSYV](http://crdd.osdd.net/raghava/ifnepitope/pep_design.php?sequence=GNRNPYKKPGLLSYV&method=hybrid&model=main) | SVM | NEGATIVE | -0.60431852 |
| 373 | 365-380 | [LPQGSVITVQKKAKF](http://crdd.osdd.net/raghava/ifnepitope/pep_design.php?sequence=LPQGSVITVQKKAKF&method=hybrid&model=main) | SVM | NEGATIVE | -0.42672562 |
| 374 | 366-381 | [PQGSVITVQKKAKFV](http://crdd.osdd.net/raghava/ifnepitope/pep_design.php?sequence=PQGSVITVQKKAKFV&method=hybrid&model=main) | SVM | NEGATIVE | -0.45255155 |
| 375 | 367-382 | [QGSVITVQKKAKFVA](http://crdd.osdd.net/raghava/ifnepitope/pep_design.php?sequence=QGSVITVQKKAKFVA&method=hybrid&model=main) | SVM | NEGATIVE | -0.10420553 |
| 376 | 375-390 | [KKAKFVAAWTLKAAA](http://crdd.osdd.net/raghava/ifnepitope/pep_design.php?sequence=KKAKFVAAWTLKAAA&method=hybrid&model=main) | SVM | POSITIVE | 1.2094703 |
| 377 | 376-391 | [KAKFVAAWTLKAAAG](http://crdd.osdd.net/raghava/ifnepitope/pep_design.php?sequence=KAKFVAAWTLKAAAG&method=hybrid&model=main) | SVM | POSITIVE | 0.79460947 |
| 378 | 377-392 | [AKFVAAWTLKAAAGG](http://crdd.osdd.net/raghava/ifnepitope/pep_design.php?sequence=AKFVAAWTLKAAAGG&method=hybrid&model=main) | SVM | POSITIVE | 0.61623789 |
| 379 | 378-393 | [KFVAAWTLKAAAGGG](http://crdd.osdd.net/raghava/ifnepitope/pep_design.php?sequence=KFVAAWTLKAAAGGG&method=hybrid&model=main) | SVM | POSITIVE | 0.67221404 |
| 380 | 379-394 | [FVAAWTLKAAAGGGS](http://crdd.osdd.net/raghava/ifnepitope/pep_design.php?sequence=FVAAWTLKAAAGGGS&method=hybrid&model=main) | SVM | POSITIVE | 0.52759093 |
| 381 | 380-394 | [VAAWTLKAAAGGGS](http://crdd.osdd.net/raghava/ifnepitope/pep_design.php?sequence=VAAWTLKAAAGGGS&method=hybrid&model=main) | SVM | POSITIVE | 0.51642487 |
| 382 | 381-394 | [AAWTLKAAAGGGS](http://crdd.osdd.net/raghava/ifnepitope/pep_design.php?sequence=AAWTLKAAAGGGS&method=hybrid&model=main) | SVM | POSITIVE | 0.4685247 |
| 383 | 382-394 | [AWTLKAAAGGGS](http://crdd.osdd.net/raghava/ifnepitope/pep_design.php?sequence=AWTLKAAAGGGS&method=hybrid&model=main) | SVM | POSITIVE | 0.29279818 |
| 384 | 383-394 | [WTLKAAAGGGS](http://crdd.osdd.net/raghava/ifnepitope/pep_design.php?sequence=WTLKAAAGGGS&method=hybrid&model=main) | SVM | POSITIVE | 0.26331446 |
| 385 | 384-394 | [TLKAAAGGGS](http://crdd.osdd.net/raghava/ifnepitope/pep_design.php?sequence=TLKAAAGGGS&method=hybrid&model=main) | SVM | POSITIVE | 0.15651694 |
| 386 | 385-394 | [LKAAAGGGS](http://crdd.osdd.net/raghava/ifnepitope/pep_design.php?sequence=LKAAAGGGS&method=hybrid&model=main) | SVM | POSITIVE | 0.20420466 |
